# Macroscopic Vibronic Coherence and Non-Dispersive Soliton Propagation Across a Protein-Solvent Interface

**DOI:** 10.64898/2026.10.01.755905

**Authors:** Kristina N. Woods

## Abstract

A foundational tenet of modern biophysics dictates that protein-solvent interactions and charge-transport dynamics operate strictly within microscopic, sub-nanometer boundaries where long-range coordination is natively suppressed by rapid room-temperature thermal dephasing. Here, we report an unexpected phenomenon that fundamentally challenges this paradigm: the long-distance, quantum-coherent communication between isolated protein samples in aqueous solution across macroscopic spatial separations. By mapping the distance-dependence of a weak, protein-emitted electromagnetic field on the response dynamics of an adjacent complex, we demonstrate that this inter-protein communication establishes a stable, intermolecular vibronic coupling operating across millimeter-scale distances up to at least 40 mm. This field interference induces a robust quantum coherence that lowers the local potential energy surface, transforming the coupled protein-solvent network into an un-attenuated, low-barrier energy transfer conduit. This stabilized pathway launches a topologically protected, non-dispersive soliton that exhibits exceptional phase stability against room-temperature thermal dephasing. These findings provide definitive empirical evidence of a sophisticated, field-driven quantum infrastructure embedded within a biological macromolecule. More explicitly, they reveal that a protein can possess an inherent capacity to preserve coherent wave propagation over macroscopic scales previously deemed impossible in room-temperature aqueous environments.

## 1. Introduction

Long-range intermolecular communication across protein-solvent interfaces remains a fundamental yet elusive frontier in quantum biophysics. While standard macromolecular energy transport is notoriously restricted by rapid thermal dephasing (*1*, *2*) (T2 ≤ 2 ps) and classical dielectric friction, certain inherent cytokine network interactions appear evolutionarily optimized to preserve signaling fidelity over extended spatiotemporal distances. Here, we demonstrate that the Interferon-gamma (IFN-γ) dimer functions as a resonant quantum waveguide capable of achieving nonlinear, long-distance wave propagation out to at least a macroscopic constraint of 40 mm. Rather than relying on high-energy external pump inputs, this macromolecular communication channel is initialized via nonlinear excitation driven by the endogenous electromagnetic field emitted from an adjacent protein complex. Ultra-broadband time-domain spectroscopy (UB-TDS) reveals that the system achieves long-lived vibronic coherences by re-tuning the intrinsic collective intermolecular vibrations that form within the native protein structure. Under spatial resonance, the potential energy surface (PES) flattens to create a delocalized, low-barrier Zundel-motif. This framework initiates a topologically protected, non-dispersive soliton wave packet that propagates ballistically out to a 10 picosecond (ps) time scale. By vibronically coupling the fast-moving charge carrier to the fourth harmonic overtone (6488 cm^-1^ = 4 x 1622 cm^-1^) of the collective protein dynamics, the system removes boundary drag and prevents classical near-infrared dissipation - forming an elegant mechanism for long-range quantum coherence in room-temperature biological systems.

Efficient, long-range charge and energy transport across protein-solvent interfaces is a foundational mechanism underpinning biological energy conversion, enzymatic catalysis, and cellular signaling networks (*3*, *4*). Within these macromolecular interactions, dynamically structured networks of hydrogen bonds (H-bonds) at the immediate hydration interface act as dedicated conduits for rapid proton translocation via a cooperative Grotthuss-type shuttling mechanism (*5*). In standard room-temperature aqueous environments, however, the structural stability of these charge-transport pathways is heavily limited by thermal fluctuations. The continuous bending, twisting, and breaking of H-bonds create a rugged PES, characterized by deep potential energy wells separated by high activation barriers. Consequently, an escaping charge carrier is typically subjected to intense local dielectric friction and conformational trapping, restricting transport to a slow, sequential hopping regime. Under these standard ambient conditions, any initial quantum phase coherence natively undergoes rapid quantum decoherence (T_2_ ≤ 2 ps), scattering its forward kinetic momentum into the surrounding thermal reservoir as random heat.

To circumvent the impediment of rapid thermal dephasing, nonlinear wave mechanics and Davydov-type soliton frameworks (*6*) have been theoretically proposed as alternative mechanisms for lossless (*7*) biological energy transport. In these nonlinear models, a traveling wavepacket avoids dispersion and dephasing by effecting a coupled energy exchange with the surrounding macromolecular network of interactions. If the fast-moving charge carrier can successfully synchronize its charge-migration frequency with the slow, collective low-frequency vibrations of the protein, the system achieves a state of sustained quantum coherence via stable vibronic coupling. This mechanical synchronization dynamically lowers the PES ahead of the wave, pulling down the local activation barriers to eliminate boundary drag. While synthetic model systems (*8*) have demonstrated quantum-tunneling via localized, low-barrier H-bonding chains of water molecules confined in nanometer-scale channels - observing long-lived quantum phase stability across large-scale, room-temperature biopolymer interfaces has remained a profound experimental challenge.

In this work, we investigate the signaling response dynamics of IFN-γ in the presence of a protein-emitted electromagnetic field (*9*) of an adjacent protein complex. Specifically, we follow the nonlinear energy transport dynamics operating across the protein-solvent interface of the IFN-γ cytokine homodimer in aqueous solution. Rather than utilizing an external laser pump input to drive the system, we initialize the interfacial charge-migration process in the target protein by using the weak, endogenous electromagnetic field emitted natively by an adjacent IFN-γ complex. To systematically evaluate this coupling, we implement a discrete spatial matrix, varying the physical distance between the field emitter and the target macromolecule. We employ UB-TDS (*10*) to directly capture the continuous electric field wave of the resulting charge carriers across a continuous 10 to 10,000 cm^-1^ spectral region, focusing our analyses systematically on the extracted phase spectra. Importantly, because this field-resolved approach follows the real-time temporal wave of the infrared electric field rather than its square-law intensity, it preserves the phase information natively. This phase preservation establishes a unique methodology for identifying quantum correlations and long-lived vibronic coherences that are otherwise buried under bulk solvent noise. Two-dimensional (2D) correlation spectroscopy is utilized on the spectral datasets to map the cross-peak frequency associations, as well as the intramolecular and intermolecular frequency shifts modulating the protein-solvent interface. This ultrafast spectroscopic phase mapping is complemented by a phase-sensitive heterodyne detection method that traces the real-time interference of the emitted electromagnetic fields of the two protein samples at defined distances – allowing us to directly extract the field-induced electrodynamic coupling profile of the protein samples. By monitoring the phase variations driven by the coherent spatial interference of the overlapping fields, this dual (phase) approach identifies a nonlinear coupling mechanism that supports a vibronically coherent energy transfer pathway that operates across macroscopic, millimeter-scale distances. Explicitly, it identifies an unprecedented biological communication channel that preserves the propagation of a delocalized, non-dispersive soliton over scales previously assumed to be unfeasible in room-temperature aqueous environments.

## 2. Results

### 2.1 Inter-Cavity Phase Confinement and Coherent Electrodynamic Trajectories

We have mapped the macroscopic electrodynamic coupling profile of a cavity formed between two IFN-γ samples, as illustrated in Figure 1f. The primary empirical observables mapped in these experiments are the time-dependent phase velocity (Θ(*t*)) and amplitude (*R*(*t*)) fields of the electromagnetic boundary-layer interference generated by the emission of the two protein samples forming the cavity, which we measure via a heterodyne lock-in detection method (Supplementary Figures S13 a - b). Across all experimental spatial coordinates, the instantaneous frequency deviation (Δ*f*) relative to the mechanical modulation reference remains below 0.6 Hz (Table I). This constraint establishes that the coupled system functions within a regime of passive, modulation-locked phase confinement, completely decoupled from external driving fields or autonomous high-frequency macroscopic current generation.

**Figure 1.**
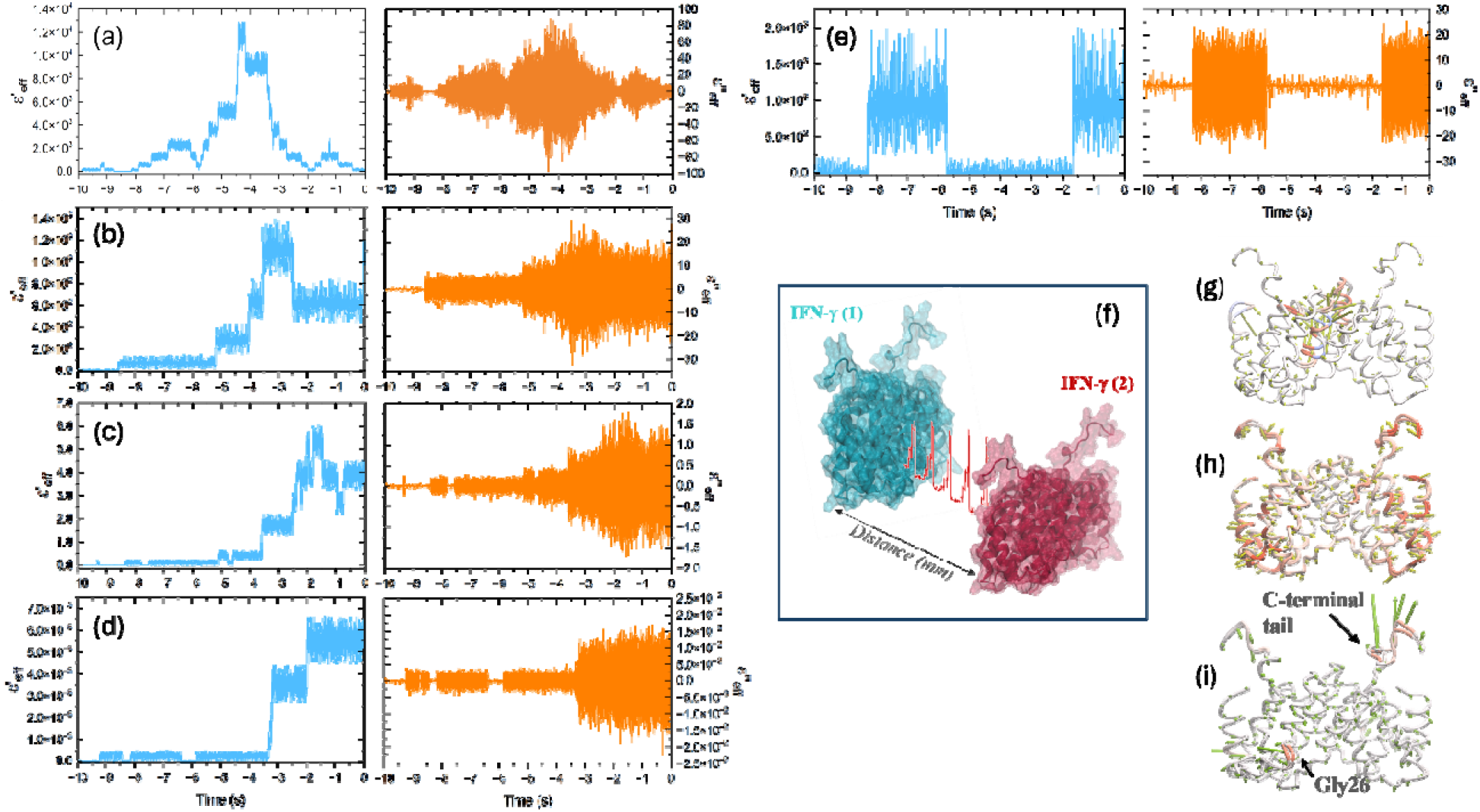
Experimental detection of the effective complex dielectric function of emitted fields and frequency-dependent macromolecular fluctuations in IFN-γ ensembles. The time-resolved real (cyan) and imaginary (orange) part of the effective complex dielectric function of the **(a)** 2.0 mm distance separation sample, **(b)** 3.0 mm distance separation sample, **(c)** 15.0 mm distance separation sample, **(d)** 40.0 mm distance separation sample of IFN-γ and **(e)** the single IFN-γ sample determined by heterodyne lock-in detection. **(f)** Schematic representation of the heterodyne detection configuration mapping the inter-protein electromagnetic field coupling across millimeter-scale spatial separation distances (*d*) between adjacent IFN-γ solution samples. **(g)** Molecular representation of the dominant equilibrium conformational mode from the MD simulation control protein trajectory (no external electric field) via Full Correlation Analysis (FCA). **(h)** Solvent-exposed helical fluctuation mode resolved via FCA under the 150 GHz external electric field (associated with the 2.0 mm close-range sample boundary). **(i)** Conformation mapping of the localized compression and hinge fluctuations of the flexible Gly26 loop and C-terminal tail residue cluster resolved via FCA. This structural coordinate maps a secondary bifurcated fluctuation under the 150 GHz field and becomes the single collective transition under the 7.5 GHz external field (associated with the 40.0 mm far-field node).

**Table I:**
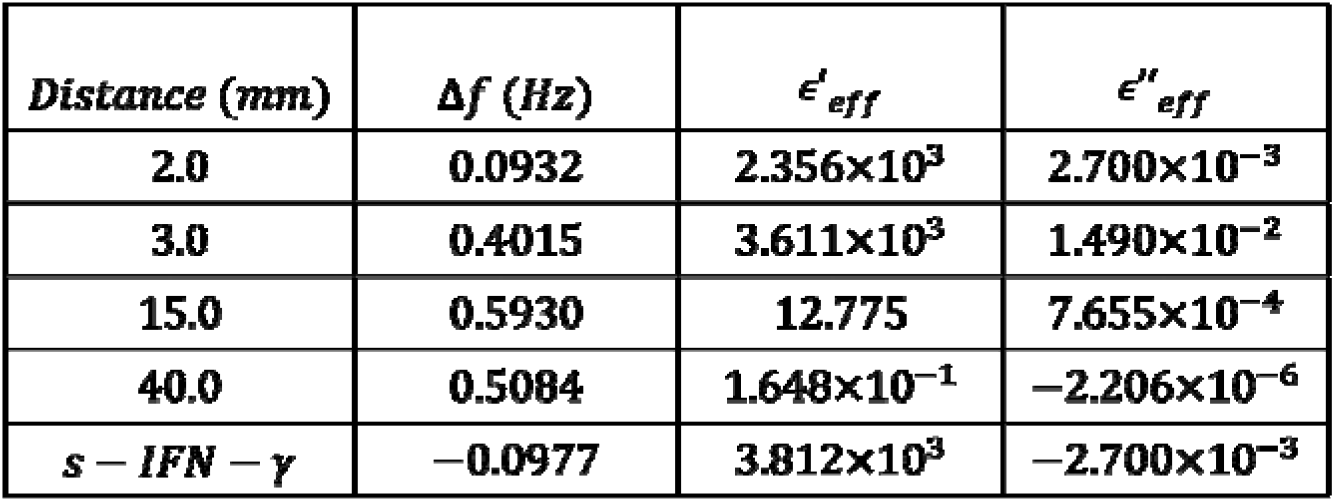
The extracted parameters of the emitted IFN-γ fields as a function of sample cavity separation distance. The frequency offset () represents the time-resolved phase rotation rate, which follows the dynamic frequency shift introduced by the sample within the distance-dependent cavity resonator. The effective real permittivity () and effective imaginary loss () denote the dielectric storage and loss components extracted from the demodulated signal across the respective protein-protein separation distances. Further, we also include the parameters for the single protein (s – IFN-γ) for comparison.

| *Distance* (*mm*) | Δ*f* (*Hz*) | ϵ′<sub>*eff*</sub> | ϵ″<sub>*eff*</sub> |
|---|---|---|---|
| 2.0 | 0.0932 | 2.356×10<sup>3</sup> | 2.700×10<sup>−3</sup> |
| 3.0 | 0.4015 | 3.611×10<sup>3</sup> | 1.490×10<sup>−2</sup> |
| 15.0 | 0.5930 | 12.775 | 7.655×10<sup>−4</sup> |
| 40.0 | 0.5084 | 1.648×10<sup>−1</sup> | −2.206×10<sup>−6</sup> |
| *s* − *IFN* − *γ* | −0.0977 | 3.812×10<sup>3</sup> | −2.700×10<sup>−3</sup> |

Further, to resolve the polar field components into fundamental thermodynamic coordinates, the time-series arrays were converted into a time-resolved effective complex dielectric function evaluated at each discrete separation distance:

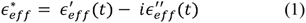

where the real part 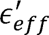 is derived from the spatial phase gradient, mapping the effective field-induced dipole polarization state and localized electric displacement field storage capacity (***D*** = *ɛ*_0_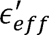***E***) within the intervening cavity volume. The imaginary part 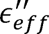 is extracted from the time derivative of the measured logarithmic amplitude of the field, defining the localized dielectric dissipation or reactive absorption profile.

#### 2.1.1 Spatial geometric filtering of collective macromolecular motions

The specific free-space separation distances (*d*) established between the independent samples function as a spatial geometric filter. According to classical wave boundaries, a spatial distance separation acts as a resonant cavity that selectively confines high-frequency electromagnetic carrier fields whose wavelengths (*λ*) match the half-wavelength constraint (*d* = *mλ*/2). The slow temporal modulations captured by our detection system monitor the macroscopic phase stability of these modes over time.

Explicitly, the high-frequency electromagnetic carrier continuum bounded within the cavity is directly associated with the large-amplitude collective modes of the protein, where the macromolecular network interacts directly with the local solvent environment. Within this un-driven operational regime, the emergence of highly structured, time-dependent permittivity profiles unique to the protein structure (Supplementary Figure S1) indicates that the global structural fluctuations undergo organized, non-linear state-switching and structural latching events rather than dispersing randomly into the thermal background (*k_B_T*). The coupled architecture establishes a self-sustaining macro-dipole state that actively manipulates local dielectric coherence across the separation distance.

#### 2.1.2 Distance-dependent analysis of coupled protein interactions

The spatial progression of the complex permittivity parameters across the free-space protein separation distances resolves into distinct regimes, demonstrating that the inter-protein field interactions are governed by nonlinear, cooperative macroscopic electrodynamics:

##### 2.1.2.1 The 2.0 mm near-field resonance condition

At a separation distance of 2.0 mm, the system exhibits the classic signatures of a macroscopic resonance condition, wherein the protein macro-dipoles undergo a highly coordinated, time-delimited cooperative alignment. As shown in Figure 1a, the effective real permittivity (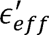) maps out a globally symmetric, peaked resonant envelope that opens at approximately *t* = −8.0 *s*, reaches its maximum alignment apex at the midpoint *t* = ∼ − 4.5 *s t* − 4.2 *s*, and returns symmetrically to its initial disordered baseline by *t* = 0 *s*. This macro-envelope represents a transient, highly structured surge in the macroscopic electric susceptibility of the medium. At this narrow distance separation, the near-field dipole-dipole interactions between the protein samples are optimized to establish a transient cooperative state, temporarily storing electrodynamic energy as the system ascends the resonant curve.

The fine structure of this resonant curve is outlined by a sequence of step-wise transitions. The system ascends the leading edge of the resonance via a series of discrete upward step-jumps at *t* = −8.1 *s*, −6.8 *s*, and −5.1 *s*, before undergoing a sharp, near-vertical climb into its maximum permittivity plateau state (centered around 1.3 × 10^4^). After passing the symmetric apex of the resonance at *t* = −4.2 *s*, the system descends the trailing edge of the curve through an equally structured, cascading sequence of downward jumps at *t* = −3.3 *s*, −2.7 *s*, and −1.8 *s*. The symmetry between the ascending and descending pathways suggests that this interference-driven polarization and subsequent depolarization process is globally balanced and thermodynamically reversible.

Concurrently with the real part, the effective imaginary permittivity (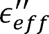) forms a localized, globally symmetric envelope of high-frequency fluctuations centered at the resonant midpoint (*t* ∼ − 4.5 *s t* − 4.2 *s*). As shown in Figure 1a, the amplitude of these fluctuations opens from a narrow baseline at approximately *t* = −8.0 *s*, expands proportionally with the magnitude of 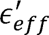 to its maximum breadth (oscillating between approximately −80 and +80), and decays symmetrically back to a narrow baseline as the timeline approaches *t* = 0 *s*. The signal continuously oscillates symmetrically across the zero baseline throughout the active window. According to the fluctuation-dissipation theorem (*11*), the imaginary component directly maps the dissipative processes and out-of-phase fluctuations of the system. The symmetric expansion around zero indicates that the system is not undergoing a net directional energy loss, but rather a transient surge in high-frequency, phase exchange between the self-emitted field and the localized macro-dipoles. As the protein dipoles cooperatively reorient (driving the peak in 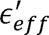), they undergo a temporary period of heightened dielectric friction and stochastic phase fluctuations. Once the system passes the resonant apex and decouples, these dissipative fluctuations are quenched, and the medium returns to its baseline uncoupled state by *t* = 0 *s*, where the protein macro-dipoles oscillate independently and thermally dissipate into the surrounding environment.

In summary, the 2.0 mm sample distance interaction is governed by short-range, near-field electrodynamic coupling. The proximity allows the two samples to act as a weak, localized perturbation that co-aligns the protein macro-dipoles temporarily. However, the spatial boundary conditions at 2.0 mm do not support a sustained interference field capable of phase-locking the macro-dipolar network, causing the system to relax entirely back to its initial disordered equilibrium via a thermodynamically reversible pathway.

##### 2.1.2.2 3.0 mm cavity configuration: structural polarization hysteresis and metastable state latching

Increasing the separation distance to 3.0 mm induces a fundamental qualitative shift in the system’s thermodynamics. The system transitions from the reversible, transient resonance observed at 2.0 mm into a nonlinear, multi-stable polarization state, where the medium becomes structurally trapped in a sustained, higher-ordered configuration.

The effective real permittivity (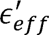) remains at its minimum baseline until approximately *t* = −8.5 *s* (Figure 1b), where it initiates a series of discrete, multi-step discontinuous jumps. Rather than transitioning via a single, simple threshold, the system cascades through a sequence of distinct intermediate plateaus at *t* = −5.2 *s* and *t* = −4.1 *s*, before undergoing an abrupt upward step at *t* = −3.6 *s* into a maximum alignment state. This step-function behavior is a signature of a multi-stage, first-order-like structural phase transition driven by electrodynamic feedback. At 3.0 mm, the spatial separation distance alters the boundary conditions of the cavity, allowing the self-emitted field to build up and cross a succession of critical thresholds. Once these thresholds are attained, the protein macro-dipoles and their surrounding hydration networks undergo a cooperative, nonlinear reorganization. The peak state is sustained for approximately one second until *t* = −2.5 *s*, where the system undergoes a sudden, discontinuous downward step. Rather than returning to equilibrium, the medium stabilizes into an elevated plateau between *t* = −2.5 *s* and *t* = 0 *s*. This sustained state marks the transition into a long-lived metastable configuration characterized by enhanced macroscopic electric susceptibility and a persistent collective polarization.

Initially, the imaginary component exhibits a quiescent baseline centered around zero. Coincident with the first step-jump in 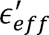 at *t* = −8.5 *s*, this baseline abruptly expands into a dense, high-frequency envelope that oscillates symmetrically about zero. As the real component cascades through subsequent plateaus, the dissipation envelope mirrors these transitions, expanding further at *t* = −5.2 *s* and reaching its maximum amplitude between *t* = −3.6 *s* and *t* = −2.5 *s*, which is synchronized with the peak alignment state of 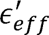. Following the partial downward relaxation at *t* = −2.5 *s*, the envelope narrows slightly but remains wide, highly active, and sustained up to *t* = 0 *s*. The step-wise widening of the 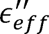 envelope signifies a multi-stage shift in the system’s dissipative pathway. The symmetric, bipolar oscillation around zero signifies that the newly formed macroscopic state is continuously and dynamically exchanging phase and momentum with the local electromagnetic field (*11*). Rather than a brief burst of dielectric friction followed by rapid quenching (as seen at 2.0 mm distance sample), the 3.0 mm cavity configuration maintains a state of continuous, nonequilibrium energy exchange. The system is actively driven by its own self-induced field interference, utilizing the localized electromagnetic feedback to continuously sustain its ordered, highly polarizable molecular configuration against background thermal dissipation forces.

To summarize, at 3.0 mm separation the system exhibits structural polarization hysteresis. The geometric boundary conditions of the cavity allow the superimposed, self-emitted fields to achieve critical nonlinear thresholds that override local thermal dissipation. The protein sample effectively self-organizes into an active, multi-stable dielectric phase, demonstrating that small adjustments in macroscopic spatial geometry can fundamentally alter the collective thermodynamic behavior of the macromolecular network.

##### 2.1.2.3 15.0 mm cavity configuration: threshold-driven polarization screening and interference-driven phase transitions

Increasing the separation distance to 15.0 mm induces a fundamental qualitative shift in the system’s thermodynamics. The system transitions from standard isolated dielectric behavior into a nonlinear, multi-stable polarization phase, where the medium becomes structurally trapped in a sustained, higher-ordered configuration via spatial electric field interference. As illustrated in Figure 1c, the effective real permittivity (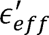) undergoes an extended incubation phase, remaining flat along its baseline until approximately *t* = −5.2 *s*, where a cascading series of discrete, multi-step discontinuous jumps are initiated. The system implements an initial minor upward step at *t* = −5.2 *s*, followed by a second distinct step-jump at *tt* = −3.6 *ss* to an intermediate plateau. At *t* = 2.5 *s*, a third transition accelerates into a sharp, near-vertical upward jump at *tt* = −2.0 *s*, promoting the medium into its maximum alignment state. This intricate, stepped response establishes a multi-phase, discontinuous structural reorganization propagated by self-induced electromagnetic feedback. This maximum alignment state is sustained for precisely half a second until *t* = −1.5 *s*, where the system undergoes a sudden, discontinuous downward step. Following a brief, transient descent between *t* = −1.0 *s* and *t* = −0.8 *s*, the medium stabilizes and latches into a sustained, intermediate elevated plateau that remains locked through the *t* = 0 *s* timeline. Interestingly, the multi-staged, threshold-driven jump behavior observed in the time-profile suggests that when the self-emitted waves constructively interfere within the sample volume, the localized field intensity triggers a sequence of critical nonlinear transitions. This promotes a collective, step-wise reorientation of the macro-dipoles into highly aligned, long-lived metastable states that sustain significantly higher gains in macroscopic electric susceptibility.

The imaginary component (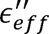) exhibits a latent baseline centered at zero preceding the transition that was previously observed in the real component. Coincident with the initial structural activation of the system in 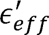, the imaginary component profile initiates a series of step-wise expansions. A minor onset of activity occurs between *t* = −8.5 *s* and *t* = −7.8 *s*, followed by a distinct widening of the fluctuation envelope at *t* = −5.2 *s*, matching the first step-jump in 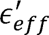. At *t* = −3.6 *s*, the imaginary channel expands significantly into a broad, dense, high-frequency envelope that oscillates symmetrically across the zero baseline. The amplitude of these fluctuations continues to step upward, reaching the maximum of the dielectric loss factor between *t* = −2.0 *s* and *t* = −1.5 *s*, which is coordinated with the maximum alignment state of the effective real permittivity. Following the downward step-jump at *t* = −1.5 *s*, the envelope narrows slightly but remains highly dynamic and sustained up to *t* = 0 *s*. The stepwise expansion of the 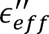 profile represents a restructuring of the system’s dissipative channels, establishing a multi-tiered landscape for energy management. The symmetric, zero-centered bipolar oscillations denote that this emergent macroscopic state undergoes a continuous, coherent exchange of phase and momentum with the local electromagnetic environment. Consequently, the 15.0 mm cavity circumvents the typical trajectory of transient dielectric friction and abrupt quenching. Rather, it sustains a persistent, nonequilibrium energetic steady state where self-generated field interference acts as a stabilizing feedback loop, shielding the highly polarizable molecular architecture against thermal randomization.

At 15.0 mm separation distance, the system does not experience a transient, self-limiting surge, but rather a sustained transition into an active, multi-stable dielectric phase. The geometric boundary conditions allow the superimposed, self-emitted fields to achieve successive nonlinear thresholds. Once locked into this configuration, the continuous high-amplitude dissipation envelope signifies that the macro-dipolar network remains driven by the field interference pattern, maintaining its phase-inverted screening structure through the end of the time-series.

##### 2.1.2.4 40.0 mm cavity configuration: long-range interference latching and critical incubation latency

At a macroscopic separation distance of 40.0 mm, the system reaches the farthest spatial limit of its collective behavior within this experimental geometry. At this distance, the free-space wavelength of ∼7.5 GHz matches the 40.0 mm boundary conditions, defining the spatial constraints required for wave superposition to cross the nonlinear threshold. The effective real permittivity (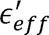) shown in Figure 1d resides at its minimum baseline until approximately *t* = −9.3 *s*, where it undergoes an initial, small discontinuous upward step into a low, fluctuating baseline state. It maintains this lower state for several seconds - displaying a transient dip between *t* = −8.2 *s* and *t* = −7.9 *s* until it reaches a major electrodynamic threshold at *t* = −3.3 *s*. At this point, the system implements a sharp, near-vertical step-jump into a distinct intermediate plateau with a permittivity value centered around 3.5 × 10^−3^. The system remains locked in this intermediate state until *t* = −2.0 *s*, where it undergoes a secondary sharp, discontinuous step-jump. Following this final transition, the medium latches into a maximum, highly active polarization plateau (5.4 × 10^−3^) that remains locked through to *t* = 0 *s*. These cascading transitions point to a multi-stage, first-order-like structural phase change driven by self-induced electromagnetic feedback. This staircase-like evolution demonstrates that even at macroscopic distances, the protein-solvent system undergoes a quantized, cooperative reorganization. As the local field intensity at the interference antinodes crosses successive critical thresholds, the system latches sequentially into discrete, higher-energy configurations

The imaginary component (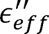) initially exhibits a minimum baseline centered about zero. Coincident with the multi-stage structural activation of the real component, the imaginary time profile undergoes a corresponding, step-wise expansion of its fluctuation envelope. The first onset of high-frequency fluctuations occurs between *t* = −9.3 *s* and *t* = −8.2 *s*, followed by a brief return to the quiescent baseline until *t* = −7.9 *s*. A second active envelope persists until*t* = −6.4 *s*, where the signal collapses back to zero baseline for a short window before re-activating at *t* = −5.9 *s*. At *t* = −3.3 *s* - coordinated with the first major upward step-jump in *ε*′ - the imaginary channel undergoes a large-scale expansion into a broad, dense envelope. This envelope steps upward a final time at *t* = −2.0 *s*, corresponding with the second major latching event in the real component. Unlike the 15.0 mm configuration, this fluctuation envelope does not quench or collapse; it continually expands to its maximum amplitude (oscillating between approximately −1.6 × 10^−2^ and +1.6 × 10^−2^) and remains fully active, dense, and sustained up to *t* = 0 *s*.

The progressive expansion and absence of thermal relaxation within the 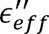 profile outlines the structural evolution of the system’s dissipative pathways under these extended boundary conditions. Again, the symmetric, zero-centered bipolar oscillations reveal that the medium participates in a continuous, high-amplitude exchange of phase and momentum with the local electromagnetic field. But in this case, in bypassing self-limiting transient decay - the 40.0 mm distance geometry functions effectively as a high-Q macroscopic resonator. Within this architecture, macro-dipoles undergo intense, continuous driving by a highly coherent, self-induced field interference pattern. This persistent, unattenuated envelope provides compelling evidence of sustained, non-equilibrium energy throughput, wherein the protein-solvent system remains actively stabilized at the anti-node of a stable, constructive wave superposition throughout the entire time-series.

##### 2.1.2.5 Isolated macromolecular kinetics: the single protein

Without a secondary source to generate a spatial interference pattern, the individual protein sample loses its stabilizing macroscopic feedback loop. As shown in Figure 1e, the effective real permittivity (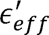) of the single IFN-γ sample remains at its minimum baseline from *t* = −10 *s* until approximately *t* = −8.3 *s*. At this point, the system undergoes an abrupt, near-vertical discontinuous jump into a dense, high-frequency fluctuating state centered around a permittivity value of 1.0 × 10^3^. This active, bistable state resembles an intense square-wave noise envelope that is sustained for nearly three seconds before undergoing a sudden, sharp collapse back to the minimum baseline at *t* = −5.7 *s*. Subsequently, the medium remains quiescent at this lower baseline for an extended window from *t* = −5.7 *s* to *t* = −1.7 *s*. At *t* = 1.7 *s*, the system experiences a second sudden, near-vertical activation, moving back into the dense, highly fluctuating envelope where it remains unconstrained through to *t* = 0 *s*.

This macro-step profile is a classic signature of stochastic bistability (*12*) and intermittent macro-state switching (*13*) in an open, driven system far from equilibrium. The IFN-γ molecules within the sample are actively generating their own internal RF fields; however, because there is no secondary sample to superimpose an external interference field, the protein-solvent network lacks the stable global boundary conditions required to stabilize a long-lived macro-state. Instead, the local macro-dipoles are caught in an intermittent thermodynamic “tug-of-war”. As the local field intensity builds, it drives the protein network into a highly fluctuating regime; however, because the field radiates into free space rather than forming a cavity-locked standing wave, the system remains structurally unstable - alternating between prolonged intervals of quiescent thermal equilibrium and bursts of high-frequency structural flipping.

The effective imaginary component (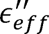) exhibits a standard, quiescent baseline centered at zero at the beginning of the time series. Mirroring the macro-state switching of the real component, the imaginary profile undergoes a synchronized, intermittent expansion of its fluctuation envelope. Specifically, the profile remains at its baseline value from *t* = −10 *s* until *t* = −8.3 *s*, where it abruptly expands into a large, dense block of high-frequency fluctuations. This symmetric envelope expands between approximately −20 and +20 across the zero baseline, maintaining its high-amplitude state for nearly three seconds before undergoing an abrupt collapse back to the near zero baseline value at *t* = −5.7 *s*. The system remains quiescent through an extended window until *t* = −1.7 *s*, where it undergoes a second sudden activation, expanding back into the dense, highly fluctuating symmetric envelope through to *t* = 0 *s*.

This intermittent bursting directly reflects non-stationary phase transients and localized dielectric friction within the unconfined medium. The broadened fluctuation envelopes match intervals where the effective real permittivity is driven into an elevated, fluctuating state. These symmetric bursting packets chart a temporary surge in the spectral density of the system’s internal, out-of-phase fluctuations (*11*). Lacking a cavity-stabilized energy exchange, the network alternates between quiet thermal equilibrium and windows of intense, bipolar phase exchange between the self-emitted field and the local macro-dipoles. As uncoordinated protein molecules reorient, they experience heightened dielectric friction, dissipating the excess energy generated by their uncorrelated macromolecular movements.

Explicitly, the isolated protein sample behaves as an unlocked, nonlinear oscillator. This demonstrates that IFN-γ natively possesses the energetic drive to self-organize and fluctuate between distinct polarization configurations at room temperature; however, it also underscores that an unconfined protein network cannot achieve long-range stabilization independently. Without a secondary sample to superimpose an external field and generate a stable spatial interference pattern, the system lacks a macroscopic anchor to suppress this stochastic flipping. It requires the geometric constraints of a configured cavity (such as the 15.0 mm or 40.0 mm cavity configuration) to arrest the system’s phase fluctuations, persistently locking the macro-dipolar network into a single, cohesive, and long-lived macroscopic phase.

Ultimately, these results demonstrate that IFN-γ does not behave as a passive protein-solvent system (Supplementary Figure S1), but rather acts as an active, tunable soft-matter bio-metamaterial. By utilizing the spatial electric field interference patterns generated by their own endogenous emissions, these protein networks successfully override room-temperature thermal noise. This self-induced electromagnetic feedback drives discontinuous macroscopic state-switching and structural latching events, transitioning the system out of an uncoupled equilibrium baseline and stabilizing it within highly persistent, phase-locked electrodynamic configurations.

### 2.2 Spatial Frequency Scaling and Mechanical Translation of Electromagnetic Carrier Fields

To further elucidate the molecular mechanisms driving the frequency-specific modulations observed experimentally at the spatial distances measured in Figures 1a – 1e, we have conducted molecular dynamics (MD) simulations of the protein homodimer under the influence of external electric fields. We have taken the two limiting distances of our experimental measurements at 2.0 mm and 40.0 mm and applied external fields associated with the carrier wave frequencies of these distances using the following relationship,

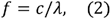

where *c* is the speed of light (2.998 ×10^11^ mm/s) and λ is the physical wavelength associated with the sample separation distance, parameterized based on the single-pass electrodynamic propagation time across the spatial distance. Subsequently, we have applied Full Correlation Analysis (FCA) on the MD simulation trajectories (see Supplementary text, *Atomistic mechanisms of field-driven dynamics via Full Correlation Analysi*s). By computing the nonlinear mutual information (*I*) across the principal functional modes (*14*), we have mapped the reorganization of the protein dimer’s conformational landscape under both control conditions and at the oscillating external electric field frequencies (150 GHz and 7.5 GHz).

We have found that the introduction of oscillating directional carrier fields dramatically alters the native network of protein interactions, transforming the structural pathways into frequency-dependent regimes. The application of these directional external fields does not merely perturb the native macromolecular dynamics but fundamentally reconfigures the underlying energy landscape by modifying the base dynamics of the control protein. In the control protein, the homodimer dynamics is dominated by an integrated network of interactions involving the nonlinear coupling between global collective core fluctuations and localized catalytic sites as shown in Figure 1g. Under the 150 GHz carrier field (associated with the 2.0 mm distance sample), the native core-to-proton-donor coupling (Figure 1g) is effectively overridden and instead the protein dynamics is almost equally split into a bifurcated network characterized by conformations within two main substates: the first mode involves high-amplitude fluctuations along the solvent-exposed outer alpha-helices of the protein (Figure 1h) and this is paired with a secondary mode with slightly higher transition probability involving the C-terminal loop compression elongation coupled with the adjacent flexible Gly26 hinge region (Figure 1i).

Conversely, shifting the external driving frequency to the 7.5 GHz carrier field (associated with the 40.0 mm far-field node) completely suppresses these multi-component modes, promoting a single collective transition involving an intense dynamical caging effect focused exclusively within the localized C-terminal and Gly26 cluster (Figure 1i). By constraining the flexible loop hinges into a single, field-synchronized pathway, the 7.5 GHz field effectively freezes out the decentralized background thermal fluctuations, organizing a highly rigid, low-dispersion macromolecular resonator capable of sustaining the phase-coherent dynamics recorded across macroscopic distances.

### 2.3 2D Spectral Mapping of the Proton Potential Energy Surface

To evaluate the immediate electronic and structural transitions driven by the inter-protein field interaction, we have analyzed the sub-picosecond transient spectral topographies captured via UB-TDS in the 10 - 10,000 cm^−1^ spectral region. The 2D contour maps in Figures 2a – d and 2f provide a direct visual mapping of how the emitted field interference modulates the nuclear reorganization and the solute-solvent interaction network of the protein as a function of distance. Within the 0.2 – 0.8 ps timescale visualized, all the protein separation distances exhibit active dynamics driven by an underdamped vibrational coupling. This coherent energy exchange oscillates periodically between the α-helical protein backbone spectral region (∼1700 cm^-1^) and the broad water combination bands (2500 – 5500 cm^-1^) that are associated with the solute-solvent mode mixing that underlie the proton-coupled electron transfer (PCET) reaction along the solvent collective coordinate that stabilizes the initial charge distribution. The prominent vertical black lines that transect the spectral maps represent the active vibrational coherences of the molecular ensemble as it moves through this coordinate. It is the visual topology of the contours from the spectral map in the < 0.3 ps region that provides some insight into the magnitude of (electron-proton) vibronic coupling at each protein distance and allows us to deduce if the mechanism is operating as a vibronically non-adiabatic or a vibronically adiabatic PCET reaction.

**Figure 2.**
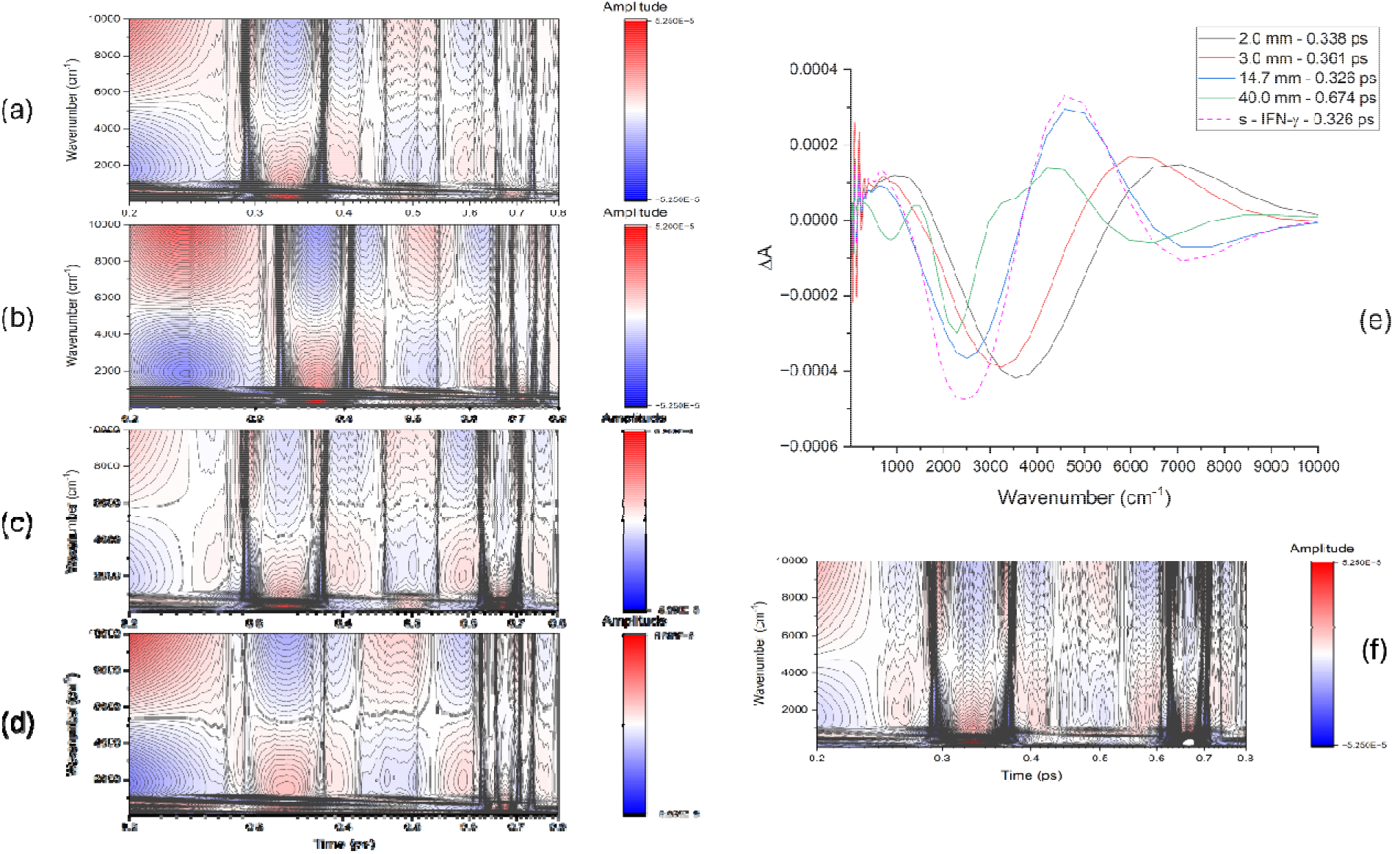
UB-TDS phase spectral maps and initial transient state spectra across the spatial matrix. (**a–d, f**) 2-D time-frequency spectral contour maps displaying the phase spectra of the UB-TDS dataset within the early time interval from 0.2 ps to 0.8 ps (200 fs to 800 fs) for **(a)** the 2.0 mm distance sample, **(b)** the 3.0 mm distance sample, **(c)** the 14.7 mm sample, **(d)** the 40.0 mm distance sample, and **(f)** the single protein control (s-IFN-γ). The contour matrices have been smoothed using a 2-D 50-point Savitzky–Golay digital spectral filter. The color scales denote positive absorbances as red and negative ground-state bleaches as blue. **(e)** Resolved transient phase spectra corresponding to the first time constant extracted from empirical mode decomposition (EMD) across the 10 to 10,000 cm⁻¹ spectral region. This initial time constant corresponds to the dynamics associated with the structural pre-organization required for the formation of the macroscale adiabatic pathway. The individual traces represent the 2.0 mm distance sample (black solid line, τ_1_ = 0.338 ps), the 3.0 mm distance sample (red solid line, τ_1_ = 0.361 ps), the 14.7 mm distance sample (blue solid line, τ_1_ = 0.326 ps), the 40.0 mm distance sample (green solid line, τ_1_ = 0.674 ps), and the single protein control (magenta dotted line, = τ_1_ 0.326 ps).

#### 2.3.1 The vibronically non-adiabatic single protein (control)

In the single protein (s – IFN-γ), the sub-300 fs region resolves four distinct, vibrational modes that comprise both transient absorbances and ground-state bleaches (GSBs) in Figure 2f. Here, GSBs are negative and colored blue in the spectral maps, while absorbances are positive and colored red in the spectral maps. The topography map shows the vibronic coupling between the slow collective movements of the protein structure and the high-frequency vibrations of the solvent H-bonding network. In the time period close to 250 fs, the spectral map clearly illustrates that in the single protein the low-frequency protein backbone reorganization modulates a series of higher vibrational overtones spanning from the fundamental Amide I (∼1650 cm^−1^) and Amide II (∼1550 cm^−1^) vibrational modes out to a near-infrared tail near 6500 cm^-1^. At very early times ≤ 200fs there is also a partially resolved absorbance band stretching from 2500 cm^−1^ up to 5500 cm^-1^ and a prominent bleach extending from 6000 out to at least 10,000 cm^-1^. These earlier time modes map the localized hydronium potentials and the bending/stretching combination modes of the interfacial water fluctuations. The four distinct spectral features representing the vibrational modes of the protein (donor) and solvent (acceptor) in the ≤ 300 fs region are consistent with a two-minima potential energy curve characteristic of a vibronically non-adiabatic PCET mechanism, where the reactant and product states remain separated by a high barrier. Interestingly, it also becomes apparent when examining the > 300 fs time region that there is a clear degradation in the “distinctness” of the vibrational modes that are visible in the spectral map. This blurring effect signifies that within this time period the system undergoes rapid vibrational dephasing under room-temperature conditions.

#### 2.3.2 The proximal field adiabatic profile (2.0 mm distance)

At a distance of 2.0 mm (Figure 2a) the protein field interaction triggers an immediate transition into a strongly coupled vibronically adiabatic PCET regime. Here, there is a deep GSB that expands broadly into the low-frequency region below 2000 cm^-1^ up to about 5000 cm^-1^. There is also a broad absorbance that extends from 6000 cm^-1^ out to the 10,000 cm^-1^ spectral limit of our experimental window. Unlike the single protein in Figure 2f, the transient absorbances and bleaches in the < 300 fs time region align vertically along the frequency axis across the entire 10 - 10,000 cm^-1^ spectral range. The fact that these phase-derived transient absorbances and GSBs span across the entire spectrum synchronously illustrates that there is a highly coordinated vibronic coupling network operating across the protein-solvent interface. It also demonstrates the proton wavepacket has delocalized from its initial localized coordinates by means of an underdamped cyclic energy exchange that now spreads smoothly between the α-helical backbone atoms and the solvent environment, which is consistent with a vibronically adiabatic mechanism. Vibronic coupling is strongly modulated by the proton donor-acceptor distance (***R***). We observed in Figure 1 that the emitted field interaction between the two proteins at a 2.0 mm distance was directly correlated with an increase in large-scale protein structural fluctuations. It is likely that the field-induced dynamics in the protein drastically alter (***R***) – greatly increasing the vibrational overlap and merging the two distinct states so that the proton and electron move together on a single, continuous energy path.

#### 2.3.3 The red-shifted adiabatic reaction (3.0 mm distance)

Shifting the spatial boundary to a 3.0 mm distance (Figure 2b) also shows an adiabatic profile but reveals a distinct spectral modification when compared to the 2.0 mm distance sample in Figure 2a. Specifically, the initial vibrational resonances exhibit a pronounced red-shift and a noticeable time delay in the < 300 fs time region. From our experimental measurements on the emitted fields in Figure 1, we have noted that at 3.0 mm the field amplitude is considerably attenuated when compared with that at 2.0 mm. For an adiabatic mechanism, the reaction is assumed to take place entirely on the lowest-energy electronic and vibrational ground state via strong vibronic coupling. Within this context, the observed red-shift and kinetic lag suggest that the underlying effect of the field is somehow muted at this distance – leading to a modified potential energy curve with higher solvent reorganization energy (**λ**) and weaker vibronic coupling. The solvent must spend more time rearranging before the reactant and product states can achieve vibronic degeneracy.

#### 2.3.4 The sharpened non-adiabatic profile (14.7 mm distance)

The 14.7 mm distance sample in Figure 2c reveals a spectral topography that closely resembles the single protein control. Both the shape and intensity of the transient absorbances and GSBs mirror that of the single protein in Figure 2f in the ≤ 300 fs region of the spectral map. However, a highly significant distinction arises past 300 fs: while the single protein experiences rapid vibrational dephasing, the 14.7 mm distance sample sustains vibrational modes with high structural clarity across the entire timeline. We initially surmised from our analysis in Figure 1 that the 14.7 mm distance position represented a node in the spatial phase interference effect of the protein emitted fields. However, the contour profile in the spectral maps shown here seem to indicate otherwise. At this distance the sample appears to adopt a two-minima, high-barrier non-adiabatic PCET framework like the single protein (Figure 2f). But unlike the single protein, the well-defined persistent sharpness of the spectral features across the timeline indicates that the field actively influences the environment to maintain the vibrational modes against room-temperature dephasing.

#### 2.3.5 The delayed far-field adiabatic reaction (40.0 mm distance)

In the extended 40.0 mm distance in Figure 2d, the sample recovers a strongly coupled adiabatic potential surface that closely resembles the spectral features of the 2.0 mm sample in Figure 2a. Although, there is one major temporal distinction. Early in the timeline, the dark, vertical lines signifying vibrational coherence in the spectral maps are visible but remain faint and low in contrast until about 600 fs. Once this initial delay window is surpassed, the dark vertical lines sharpen into the dark, prominent lines characteristic in the other samples. It is likely that this delay corresponds to an extended solvent reorganization phase that appears to be distinct in this long-distance spatial coherence matrix.

### 2.4 Formation of the Adiabatic Pathway

In Figure 2e, the first time-constant derived from the UB-TDS phase measurements captures the initiation and formation of the adiabatic pathway. It maps the system’s initial dielectric relaxation and the sub-picosecond molecular rearrangement that are necessary to reach the Franck-Condon transition state.

#### 2.4.1 The field-free baseline (single protein control)

At 326 fs, the single protein exhibits a deep, wide bleach centered at 2500 cm^-1^ that reflects the ground state bleach of a strongly hydrogen-bonded, O-H population. In this case it is likely associated with the donor aspartate (Asp41) carboxylate O–H bond in IFN-γ. This is combined with a positive absorbance at 4600 cm^-1^ that reflects a vibrational combination band populated by the initial field-driven excitation. This feature serves as a distinct diabatic signature characteristic of a non-adiabatic proton transfer regime, where the deep, narrow potential energy wells of the reactant and product states remain structurally isolated. In this uncoupled configuration, the non-equilibrium wavefunctions localized within the donor core mix only weakly through high-frequency combination states, confirming that in the absence of an external inter-protein field to modify the landscape, the baseline protein framework is restricted to a non-adiabatic transport pathway.

#### 2.4.2 The interfacial fusing mode (2.0 mm distance)

At 338 fs, the 2.0 mm distance sample is characterized by a broad positive absorption band centered at about 7000 cm^-1^. The 7000 cm^-1^ band represents an optical intervalence charge transfer (IVCT) state that is indicative of a concerted electronic and proton delocalization, driven by maximum interfacial electronic coupling across the homodimer interface. In this concerted regime, the electronic delocalization involves the tryptophan donor and its cross-interface electronic acceptor. While proton delocalization involves the proton donor (Asp41) and the network of solvent molecules in the residue vicinity. In addition to the IVCT band there is also a bleach at 3540 cm^-1^ and a broad absorbance extending from about 300 – 1400 cm^-1^. The 3540 cm^-1^ absorbance is the signature of free, non-hydrogen-bonded or weakly bound O–H stretching modes belonging to water molecules at the edge of the protein hydration layer. The broad vibrational Zundel-type continuum envelope extending from 300 – 1400 cm^-1^ signifies that the proton barrier has collapsed, allowing the proton to oscillate freely within the flat well of the shared vibronic wavefunction. Together, these spectral signatures show that the entire surface water layer has fused into the electronic transition. The system foregoes any distinct, separate step-by-step chemical steps. Instead, it uses the solvent molecules to build a continuous, delocalized environment where the electron and proton move smoothly as a single, unified wavepacket. Overall, these spectral features provide chemical and physical confirmation of a concerted, vibronically adiabatic PCET mechanism at this distance.

#### 2.4.3 The mechanically deformed channel (3.0 mm distance)

At 361 fs, the 3.0 mm distance spectrum shows a broad absorbance centered at 5950 cm^-1^. There is a prominent red-shift in absorbance band frequency when compared with the (∼7000 cm^-1^) IVCT band in the 2.0 mm distance spectrum (Figure 2e). The peak energy of an IVCT band in strongly coupled systems is directly correlated with the electronic and solvent barrier (E_op_); therefore, this red-shift indicates that the vertical gap between the ground state and the excited state has decreased. This occurs when the charge states become more localized. This premise is further supported by the bleach at about 3244 cm^-1^ that denotes the ground-state population of a compressed, localized proton-donor H-bonded stretch. Based on the experimental spectrum, a very specific story begins to unfold. The attenuated yet still intense field at this distance causes immediate ground-state destabilization. The field applies a force that physically distorts the donor away from its field-free conformation, shifting the proton closer to its own oxygen core. This structural distortion (3244 cm^-1^ bleach) raises the internal energy ground state of the reactant, mechanically driving it closer to the transition state. As a result of the field-induced structural deformation, there is a concurrent reduction in electron delocalization across the homodimer interface. This is the source of the red-shift of the IVCT band in the 3.0 mm distance sample. In essence, these field-induced structural distortions effectively decrease the activation energy barrier, creating a low-barrier environment that supports a vibronically adiabatic reaction. In this context, we also observe a broad 300 – 1250 cm^-1^ absorbance continuum in the spectrum. Unlike the Zundel-type continuum detected in the 2.0 mm distance sample, there are sharp, resolved peaks in the low-frequency continuum at 312 cm^-1^, 442 cm^-1^ and 625 cm^-1^ in the 3.0 mm distance spectrum. The bands at 312 cm^-1^ and 442 cm^-1^ are assigned to low frequency acoustic phonon modes associated with the collective skeletal breathing and twisting motions of the protein surface backbone atoms. The 625 cm^-1^ mode corresponds to a hindered rotational libration of the immediate interfacial water network. Together, these spectral features demonstrate that the system utilizes a synchronized macromolecular-solvent acoustic channel to transiently lower the local potential barrier, permitting the proton wavefunction to undergo extensive spatial delocalization across the 300 – 1250 cm^-1^ continuum via an adiabatically gated pathway.

#### 2.4.4 The unperturbed non-adiabatic baseline (14.7 mm distance)

At the further distance of 14.7 mm, the protein-emitted electromagnetic field undergoes further attenuation, rendering it insufficient to promote either the instantaneous interfacial electronic coupling of the 2.0 mm distance sample or the prominent backbone distortion detected in the 3.0 mm distance sample. At the first time-constant (∼ 330 fs), the 14.7 mm distance spectrum strongly resembles the field-free single protein (control). The sub-picosecond dynamics are dominated by a deep, broad negative bleach centered near 2500 cm⁻¹, associated with the ground-state population of the tightly H-bonded donor O–H coordinate and a prominent vibrational combination absorbance centered at 4600 cm^-1^. Both spectral features provide strong spectroscopic evidence of a vibronic non-adiabatic reference baseline at the starting point of the reaction.

#### 2.4.5 The long-range polarized matrix (40.0 mm distance)

At the long-range spatial distance of 40.0 mm, the emitted field attenuates far below the threshold required to classically perturb or distort the structural dynamics of the protein. However, at a delayed first time constant of 0.674 ps, the 40.0 mm sample resolves a unique, multi-mode, field-induced structural polarization of the surface hydration layer that is entirely absent at all closer investigated distances. Spectral analysis reveals that the system utilizes a long-range amplification mechanism governed by a nonlinear dielectric response at the solvent interface. A resonant coupling between the emitted field and the collective librational rocking motions of weakly bound water molecules in the outermost hydration shell drives the initial response, arising as a distinct bleach at 884 cm⁻¹. In parallel, this collective dipole alignment propagates inward, altering the local dielectric permittivity and depleting the tightly bound, highly restricted surface water population. This depletion is resolved as a prominent bleach at 2294 cm^-1^, which corresponds to the first overtone (second harmonic) of the fundamental 1147 cm^-1^ Zundel stretch (1147 x 2 = 2294 cm^-1^)(*15*, *16*). As the water network reorganizes, it transitions into a highly coordinated, uniform interfacial phase that generates a prominent absorbance at 3244 cm^-1^, marking the formation of a strongly hydrogen-bonded, polarized water layer. The simultaneous emergence of a broad positive absorbance at 4400 cm^-1^ signifies the field-induced global restructuring of the solvent into an organized quantum environment. This 4400 cm^-1^ feature arises as an intensely coupled, anharmonic vibrational combination band originating from the transiently populated non-equilibrium state, comprised of the highly organized surface water stretch (ν_ΟΗ_ = 3244 cm^-1^) and the low-frequency Zundel-type shuttling motion (ν_Zudel_ = 1147 cm^-1^). Its presence maps a strong quantum mechanical mode-mixing between the translocating proton and the rearranging water network during the initial phase of the reaction. This field-driven structural polarization alters the local electric field gradient at the proton donor–solvent interface, lowering the activation barrier into a low-barrier regime and shifting the proton coordinate into a strictly anharmonically adiabatic pathway.

### 2.5 Proton Transfer Phase

#### 2.5.1 Distance-dependent mechanistic divergence of the proton transfer phase

The second time-constant shown in Figure 3a (and Supplementary Figure S11) follows the transfer of the proton from the protein to the H-bonding network of the interfacial solvent molecules. Interestingly, inspection of the data reveals a distinct shift toward longer timescales of the samples that we surmised (from Figure 2) to adopt an adiabatic mechanism compared with the samples that we identified as non-adiabatic. For instance, both the single protein (s-IFN-γ) and the 14.7 mm distance sample were identified as non-adiabatic PCET reactions and we observe in Figure 3 that they have similar time constants of 0.65 ps and 0.66 ps, respectively. In contrast, the field-induced adiabatic reactions have delayed time constants of 0.98 ps in the 2.0 mm distance sample, 0.99 ps in the 3.0 mm distance sample and 1.30 ps in the 40.0 mm distance sample. Therefore, it is likely that this temporal delay is directly correlated with the field-induced formation of a potential energy surface that supports an adiabatic process.

**Figure 3.**
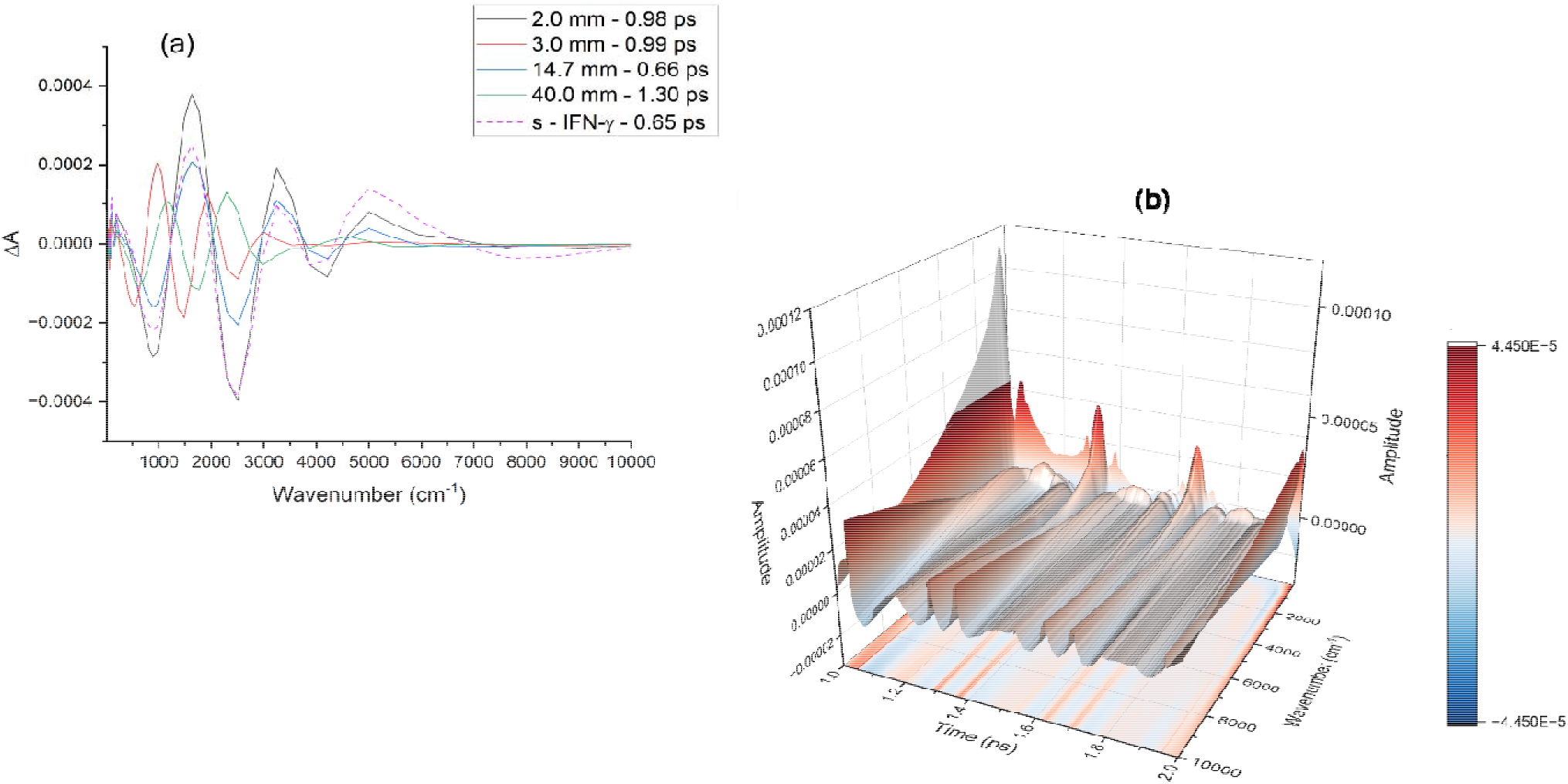
Intermediate transient state phase spectra and 3D temporal evolution of the proton-transfer coordinate **(a)** Experimentally resolved transient phase spectra corresponding to the second time constant extracted from EMD across the 10 to 10,000 cm⁻¹ spectral range. This secondary temporal component is associated with net proton translocation and structural relaxation within the solvent interfacial region. Traces show the 2.0 mm distance sample (black solid line, τ_2_ = 0.98 ps), the 3.0 mm distance sample (red solid line, τ_2_ = 0.99 ps), the 14.7 mm distance sample (blue solid line, τ_2_ = 0.66 ps), the 40.0 mm distance sample (green solid line, τ_2_ = 1.30 ps), and s-IFN-γ (magenta dashed line, τ_2_ = 0.65 ps). **(b)** 3-D spectral surface plot of the time-frequency contour map of the 2.0 mm distance sample across the subsequent 1.0 ps to 2.0 ps time delay window. The contour map has been smoothed using a 2-D 50-point Savitzky–Golay digital spectral filter to resolve the low-frequency oscillations running parallel to the high-frequency vibronic oscillations.

Despite the variations in time, the proton transfer dynamics in the single-protein (control), the 2.0 mm distance sample and the 14.7 mm distance sample are remarkably similar. In fact, the three distinct samples investigated share several common vibrational markers. The first is the 2501 cm^-1^ bleach. The GSB marks the depopulation of the strongly H-bonded aspartate carboxylate (O-H) as the proton is released from its oxygen core. The emerging transient states are observed as a coherently phase-coupled harmonic system defined by a fundamental absorbance at 1622 cm^-1^ and a corresponding first overtone (second harmonic) at 3244 cm^-1^ (1622 x 2 = 3244 cm^-1^). The 1622 cm^-1^ fundamental captures the degenerate bending mode of the newly populated hydronium core (H_3_O^+^) strongly mixed with the asymmetric carboxylate stretching vibration (ν_COO-_) of the resulting aspartate anion. The precise integer scaling of the 3244 cm^-1^ absorbance identifies it as the corresponding bending overtone (2ν_2_), which is nonlinearly enhanced via a Fermi resonance with the newly polarized interfacial O-H stretching network. Complementing this central transport coordinate are distinct GSBs resolved at 884 cm^-1^ and 4207 cm^-1^, which represent the secondary depopulation of local solvent rocking modes and high-frequency combination states, mapping the structural relaxation of the native protein hydration shell.

While these shared features confirm an identical product state, the data resolves clear spectral distinctions that expose how the external field alters the intermediate potential energy landscape. In the 2.0 mm configuration, the low-frequency region resolves a broad, prominent absorbance spanning the 200–350 cm^-1^ spectral window (Supplementary Figure S7). This low-frequency feature acts as a precise metric for a collective, field-induced structural alignment of the intermolecular (O···O) hydrogen-bond stretching coordinates within the interfacial solvent layer. This global ordering suppresses the activation barrier, driving the system into a symmetric, low-barrier hydrogen bond (LBHB) framework (*17*) where the proton wavefunction undergoes extensive quantum delocalization. The expansion of this physical reaction coordinate and its phase synchronization with the heavy-atom lattice slows the relaxation rate, extending the time constant to 0.98 ps. This phase-coherent state is confirmed by the 3D experimental coherence map (Figure 3b), where the 1–2 ps window of the 2.0 mm sample resolves highly discrete parallel ridges running straight along the temporal axis with zero diagonal tilt, demonstrating that the delocalized wavefunction is successfully shielded from stochastic liquid dephasing.

The other spectral distinction is the 5000 cm^-1^ combination band. The 5000 cm^-1^ absorbance band in Figure 3a denotes the off-diagonal anharmonic coupling between the solvent bending and stretching modes. In the single-protein control sample, this absorbance reaches maximum intensity, acting as a diabatic indicator of a steep, narrow potential well where the proton undergoes rapid, non-adiabatic tunneling due to the lack of a localized confining potential. On the other hand, the 2.0 mm distance sample has a peak with intermediate amplitude. The highly ordered interfacial solvent layer suppresses the activation barrier, reshaping the potential energy surface into an anharmonic single well. Consequently, the *v* = 0 → 2 overtone transition emerges as a spectroscopic signature of proton quantum delocalization, wherein the transition intensity is attenuated due to the spatial expansion of the nuclear wave packet across the diffuse interface. In the 14.7 mm distance sample, the peak intensity drops to a minimum intensity. This confirms that the weak, external field at 14.7 mm is sufficient to perturb the native, non-adiabatic tunneling pathway, yet it lacks the intensity required to induce the highly organized, low-barrier potential topology observed at shorter distances.

#### 2.5.2 Anharmonic vibrational states, soliton-driven pathways, and low-barrier network coupling

Both the 3.0 mm distance sample and the long-range configuration at 40.0 mm deviate significantly from the proton-transfer pathway characteristic of the field-free protein control. Rather than undergoing localized, non-adiabatic tunneling into a fluctuating hydration shell - which we observed as a 1622 cm^-1^ hydronium bending mode in the experimental spectrum in Figure 3a, these systems establish highly structured, low-barrier environments that alter the temporal and spectral landscape. On the experimental timeline, the 3.0 mm sample completes its proton transfer phase at 0.99 ps, while the 40.0 mm configuration resolves its transient vibrational signatures in the sequence at 1.30 ps.

The 3.0 mm distance sample utilizes macromolecular structural fluctuations to initiate interfacial proton transfer. Accordingly, this charge translocation is accompanied by the concurrent initialization of a propagating, non-dispersive, and self-consistent quantum wavepacket pathway at the protein-solvent interface. As illustrated in the experimental spectrum (Figure 3a), this propagating soliton manifests as a discrete vibrational (absorbance) progression exhibiting pronounced mechanical anharmonicity, resolving a fundamental transition at 964 cm^-1^ (*v* = 1) and its corresponding first overtone at 1929 cm^-1^ (*v* = 2). Interesting, while the unperturbed second overtone (*v* = 3) of this shared-proton coordinate is mathematically predicted to occur at 2892 cm^-1^, it is not resolved as an isolated peak. Instead, it undergoes an intense Fermi resonance with the fundamental 2975 cm^-1^ collective aliphatic (C-H) side-chain fluctuations localized on the protein surface immediately adjacent to the active proton donor site. This strong anharmonic mode-mixing induces a pronounced energy repulsion and allows the normally weak *v* = 3 overtone to borrow substantial oscillator strength from the allowed aliphatic transitions. The resulting precise spectral alignment at 2975 cm^-1^ provides direct evidence of a nonlinear coupling, demonstrating that the delocalized proton wavefunction of the moving wavepacket has established a coherent mechanical synchronization with the surrounding surface-exposed macromolecular structure.

Explicitly, the trajectory of the propagating wavepacket is dynamically synchronized with the collective movements of the protein’s surface-exposed aliphatic side chains. This structural coupling is driven by low-frequency acoustic backbone phonons that deform the local surface environment along the proton-transfer pathway. This structural reorganization is spectrally visualized through the GSBs of the localized side-chain modes, specifically capturing the deformation of the aspartate carboxylate skeletal rocking/bending mode at 525 cm^-1^ and the unperturbed asymmetric carboxylate stretching vibration (COO-) at 1487 cm^-1^. This tight structural synchronization establishes a highly stable, phase-coherent waveguide, effectively shielding the propagating wavefunction from rapid, room-temperature bulk thermal dephasing.

The 40.0 mm sample configuration exhibits a distinct kinetic retardation relative to closer near-field profiles, indicating that long-range charge translocation is rate-limited by the slow, collective dielectric relaxation and global orientation of distal solvent dipoles. Upon completion of this structural preorganization, deprotonation of the neutral aspartate carboxylic acid donor occurs. This event is characterized by coordinate GSBs at 1768 cm^-1^ (associated with a coupled motif of the protein ν_C=O_ stretch and the shifted flanking water bending modes of the emerging protonated cluster) and 625 cm^-1^, where the latter corresponds to a hindered rotational libration of the tightly bound surface water network. The simultaneous depletion of these states facilitates proton injection into a highly symmetric, interfacial Zundel-type solvent network. The migrating charge resolves as a prominent absorbance in the spectrum (Figure 3a) peaking near 1147 cm^-1^, which serves as the definitive spectroscopic signature of the symmetric Zundel proton-shuttling mode (H_5_O_2_^+^). Simultaneously, the interface adopts a highly ordered, nonlinear vibrational state configuration. This architecture is visualized as an anharmonic vibrational overtone progression, featuring the fundamental Zundel stretch at 1147 cm^-1^ (*v* = 1), its first overtone at 2294 cm^-1^ (*v* = 2), and its third overtone at 4588 cm^-1^ (*v* = 4). The precise integer scaling of this progression, combined with the absence of a sequential *v* = 3 transition, provides direct experimental evidence of a non-stationary wavepacket propagating along a symmetric LBHB topology, inducing extensive proton quantum delocalization along a stabilized solvent coordinate.

### 2.6 Wave Propagation Phase

The third time-constant captures the real-time spatial translation and structural evolution of the self-consistent quantum wavepacket across the macromolecular-solvent network. A comparative analysis of the TD – FRS spectra in Figure 4a (and Supplementary Figure S12) demonstrates that the boundary distance dictates the local electromagnetic field strength, which in turn, directly regulates the distinct activation timescales and the specific local solvation environment of the interfacial solvent network. The chronological sequence initiates at 0.97 ps, where the field-free protein (s-IFN-γ) and the 14.7 mm distance sample converge onto the same vibrational coordinates. Both spectra resolve an identical progression of discrete vibrational states characterized by a fundamental absorbance and sharp overtone bands at 964 cm^-1^, 1929 cm^-1^, and 2975 cm^-1^. At this step, this shared harmonic progression marks the initial initialization of wavepacket propagation across the protein-solvent interface. Although in this case, the mechanism of propagation relies on the collective dynamics of the solvent-exposed aliphatic side chains of the protein. Lacking the field-induced solvent organization required to form a delocalized, Zundel-type interfacial solvent network, the proton wavepacket synchronizes its large-amplitude oscillations with protein collective dynamics to initialize its trajectory down the timeline.

**Figure 4.**
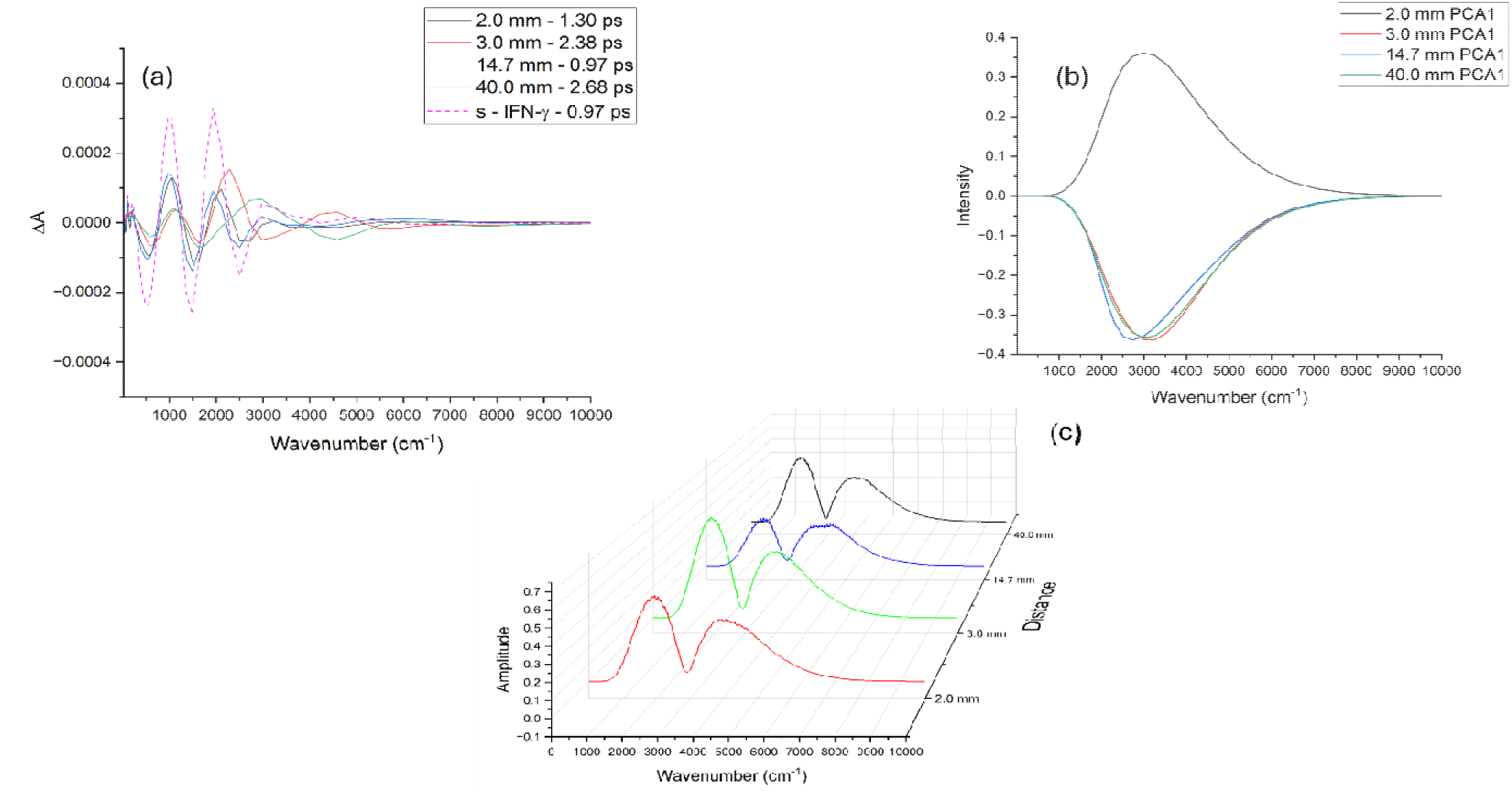
Transient phase spectra of the wave propagation coordinate and statistical principal component loading profiles **(a)** Experimentally resolved transient phase spectra corresponding to the third time constant extracted from EMD across the 10 to 10,000 cm⁻¹ spectral range. This temporal component is associated with the active wave propagation phase across the molecular ensemble. Traces display the 2.0 mm distance sample (black solid line, τ_3_ = 1.30 ps), the 3.0 mm distance sample (red solid line, τ_3_ = 2.38 ps), the 14.7 mm distance sample (blue solid line, τ_3_ = 0.97 ps), the 40.0 mm distance sample (green solid line, τ_3_ = 2.68 ps), and the single protein control (magenta dotted line, τ_3_ = 0.97 ps). **(b)** Fast Fourier Transform (FFT) intensity profiles of the primary principal component (PCA1) extracted from the experimental time-resolved phase matrices as a function of wavenumber (10 to 10,000 cm⁻¹). Traces map the baseline carrier wave loading for the 2.0 mm distance sample (black line), 3.0 mm distance sample (red line), 14.7 mm distance sample (blue line), and 40.0 mm distance sample (green line). **(c)** 3-D waterfall plot of the Fast Fourier Transform (FFT) spectral loading profiles for the secondary principal component (PCA2) mapped across the spatial distances. Traces isolate the nonlinear driver coordinates and spatial blue-chirping thresholds of the propagating wavepacket across the 2.0 mm sample (red line), 3.0 mm sample (green line), 14.7 mm sample (blue line), and 40.0 mm sample (black line).

It is important to note that the local solvation environment and phase coherence (Figures 2c and 2f) may distinguish the underlying quantum stability of these two samples. The 14.7 mm distance spectrum exhibits significantly lower spectral amplitudes when compared with the s-IFN-γ sample control. This difference in signal intensity is likely correlated with field-induced alterations of intermolecular interactions within the protein-solvent interface. For instance, from Supplementary Figure S6f we have deduced that both the librational (300 – 600 cm^-1^) and combination bending + librational modes (1900 – 2400 cm^-1^) are significantly enhanced in the map of 2D spectral correlations in the 14.7 mm configuration when contrasted with the s – IFN-γ control sample. The overall lower spectral amplitude observed in the 14.7 mm distance sample in Figure 4a is likely attributed to the increased librational motions in its dynamics compared with the control sample. Likewise, an analysis of the coherent oscillations in the distinct samples in Supplementary Figure S5b has revealed that the low-frequency modes in the 100 – 150 cm^-1^ spectral region are mostly absent in the 14.7 mm distance sample. These low frequency modes are associated with the collective intermolecular H-bond stretching and breathing motions of the solvent. They also represent a major avenue for quantum dephasing; hence, their absence suggests a suppression of low-frequency non-radiative decay channels in the 14.7 mm distance sample. Together, these observations indicate that the difference in spectral amplitude may be related to the efficiency of quantum wave propagation at the macromolecular interface. The strong coupling with high frequency interfacial solvent dynamics and the suppression of low-frequency dephasing modes would shield the propagating wave in the 14.7 mm configuration from thermal decoherence.

This behavior contrasts with the near-field solvation shift resolved at 1.30 ps in the 2.0 mm distance sample shown in Figure 4a. Driven by maximum near-field intensity, the proton wave function undergoes spatial delocalization inside a broad, single potential energy well. The delocalized wave function displays large-amplitude, highly anharmonic oscillations across the expanded interface, promoting a Zundel-solvent core progression within the H-bond network of the interfacial solvent layer. This non-harmonic progression is visualized (Figure 4a) as absorbance bands centered at 1051 cm^-1^ (*v* = 1), 2103 cm^-1^ (*v* = 2), and 3244 cm^-1^ (*v* = 3).

In this context, it is important to mention that the 1051 cm^-1^ and 1147 cm^-1^ Zundel coordinates share equivalent underlying soliton dynamics. The lower fundamental frequency at 1051 cm^-1^ reflects the highly structured arrangement of solvent H-bonds bordering the Zundel continuum in the 2.0 mm configuration. This structured water cage (200 – 350 cm^-1^) forms stronger hydrogen bonds with the outer hydrogens of the Zundel ion, which consequently elongates the central oxygen-oxygen distance of the core Zundel ion. The O···O distance changes modify the proton’s potential energy surface, causing the core proton-transfer band to red-shift. Analogous to what we surmised in the 14.7 mm configuration, the low-frequency water interactions likely suppress thermal fluctuations and provide the conditions required to sustain wavepacket propagation.

We also detect that the intense near-field environment of the 2.0 mm distance sample shifts the GSBs to 573 cm^-1^, 1622 cm^-1^ and 2728 cm^-1^. This contrasts with the 525 cm^-1^, 1487 cm^-1^, and 2501 cm^-1^ bleaches observed in the IFN-γ control and the 14.7 mm distance samples. The 573 cm^-1^ bleach is ascribed to the Trp indole ring skeletal vibration. The blue-shift from 525 cm^-1^ to 573 cm^-1^ supports the premise that the field-induced, solvent organization conformationally restricts the protein dynamics prior to wavepacket propagation. The blue-shift of the 2501 cm^-1^ feature to 2728 cm^-1^ signifies a major structural reorganization of the local solvent network; this 2728 cm^-1^ GSB is assigned directly to the localized O-H stretching vibration of an Eigen-type hydronium core (H_3_O^+^) (*18–20*). The occurrence of both the 573 cm^-1^ and 2728 cm^-1^ modes in the spectrum provides further confirmation that the external field pre-organizes these interfacial coordinates, establishing a highly interconnected, structured solvation shell immediately surrounding the active donor site. Furthermore, this structural reorganization is accompanied by a reorientation of the transition dipole moment vector of the aspartate donor group. The 1622 cm^-1^ bleach in the 2.0 mm configuration is associated with the asymmetric stretching mode of the aspartate carboxylate (ν_as_ COO-), while the 1487 cm⁻¹ bleach is ascribed to the symmetric carboxylate stretch (ν_s_ COO-). This shift in active vibrational character is significant for the transport mechanism. The transition dipole moment of the 1487 cm⁻¹ symmetric stretch points inward along the side-chain axis and directly into the hydrophobic protein core. Conversely, the dipole vector of the 1622 cm^-1^ asymmetric stretch aligns parallel with the adjacent interfacial solvent layer. Consequently, this spectral variation provides experimental indication that the intense near-field coupling in the 2.0 mm sample physically rotates the primary transition dipole moment of the starting donor group away from the protein core and projects it directly into the boundary solvent track, optimizing the interface for subsequent wavepacket propagation.

Presenting a distinct mechanistic departure from the 2.0 mm distance sample, the 3.0 mm and 40.0 mm configurations reach their wave propagation phase later in the chronological sequence. Both samples follow a mechanism where the proton wavefunction undergoes population transfer from the collective, surface-exposed aliphatic side-chains into a quantum-delocalized, low-barrier H-bonded network. Explicitly, the spectra in Figure 4a reveal that both samples have GSBs at 625 cm^-1^ and 1622 cm^-1^. Analogous to what we observed in the 2.0 mm distance sample, the1622 cm^-1^ bleach represents a field-induced transition dipole reorientation, shifting the starting dipole of the aspartate donor group into a geometry parallel to the interfacial water layer. However, contrasting with the 2.0 mm distance sample, the 3.0 mm and 40.0 mm samples do not possess interfacial structural ordering and the 573 cm⁻¹ bleach is no longer visible in the structurally relaxed local environment. Instead, the 1622 cm^-1^ mode couples its initial coordinates directly into a bulk-like H-bonded network of solvent molecules that remain electrodynamically pre-polarized by the protein-emitted field. This transition is visualized as a 625 cm⁻¹ GSB in the experimental spectrum in Figure 4a. The 625 cm^-1^ bleach is ascribed to collective solvent intermolecular torsional modes that are “pre-aligned” by the electric field to promote the subsequent population transfer of the excess proton into the delocalized Zundel-type continuum.

While both the 3.0 mm and 40.0 mm configurations utilize a common interfacial initiation point, there are clear spectral indicators that identify how the formation of the active propagating wave differs in each sample. The 3.0 mm distance sample reaches the wave propagation phase at 2.38 ps. The spectrum in Figure 4a illustrates a highly anharmonic, LBHB continuum. This continuum is visualized as a vibrational overtone progression with absorbances at the fundamental 1147 cm^-1^ (*v* = 1), 2294 cm^-1^ (*v* = 2), and 4588 cm^-1^ (*v* = 4) and a deep GSB at 2975 cm^-1^. The absence of a sequential *v* = 3 overtone, paired with the prominent emergence of the *v* = 2 and *v* = 4 transitions, characterizes the highly nonlinear response of the system. While a static, centrosymmetric potential would typically restrict transitions to even vibrational states due to rigid dipole parity selection rules (g → g), the propagation of the non-dispersive soliton manifests as a localized, non-stationary wavepacket. The periodic, non-harmonic charge oscillations of this moving wavepacket dynamically breaks the inversion symmetry of the potential on a femtosecond timescale. This dynamic relaxation of selection rules yields a Fourier-resolved series of integer harmonics, making the even overtones active and serving as a direct spectroscopic signature of a highly correlated, symmetric LBHB continuum. The spectral profile of the 3.0 mm configuration shows that active spatial translation of the wavepacket relies on the depopulation of the surface-exposed aliphatic side chains to energetically drive the transition that populates the delocalized Zundel-type solvent network.

Interestingly, this mechanical regime undergoes a reorganization in the 40.0 mm distance sample. In the 40.0 mm distance sample, the wave propagation phase occurs at 2.68 ps. As illustrated in the UB-TDS spectrum in Figure 4a, the sample comprises GSBs at 625 cm^-1^, 1622 cm^-1^, and 4588 cm^-1^, in coordination with absorbance bands at 1147 cm^-1^ and 2975 cm^-1^ along with a visible shoulder at 5950 cm^-1^. The emergence of the 1622 cm^-1^ water bending mode and the 4588 cm^-1^ combination state as GSBs indicates that long-range, field-induced polarization drives a macroscopic quantum pre-organization of the interface water dipoles. This specific pre-organization of the baseline solvent modes shifts the initial equilibrium reaction coordinate directly into the delocalized network prior to the electrodynamic activation that initiates wave propagation. Concurrently, the strong absorbance at 1147 cm^-1^ is associated with the active population of the fundamental Zundel transition, while the 2975 cm^-1^ absorbance and its corresponding whole-integer second harmonic overtone (*v* = 2) at 5950 cm^-1^ indicate that the propagating wavepacket continuously deposits a fraction of its energy back into the collective modes of the aliphatic side chains as it travels down this pre-polarized pathway. This coordinated spectral evolution provides experimental confirmation of environment-assisted quantum transport (ENAQT) (*21*), demonstrating that macroscopic spatial boundary conditions dictate spatial phase interference to eliminate the local Franck-Condon reorganization barrier, insulating the room-temperature traveling wavepacket from random liquid dephasing noise.

#### 2.6.1 Principal component analysis of carrier dynamics and adaptive chirping

To mathematically isolate the primary electrodynamic carrier from the underlying coherent nonlinear driver, a Principal Component Analysis (PCA) was performed on the UB-TDS spectra. The first principal component (PCA1, Figure 4b) captures the global variance of the macromolecular-solvent network, exposing how the bulk energy of the wavepacket “re-tunes” itself across the spatial matrix to avoid thermal dissipation. Spectroscopically, PCA1 reveals a distinct structural and directional transition across the spatial boundary coordinates, which is visualized as a sign inversion and a dynamic shift in peak positions. The 2.0 mm near-field sample presents as a positive loading spectral profile peaking at 2975 cm^-1^. This absorbance profile confirms that within the highly confined near-field environment, the carrier wavepacket initializes by actively populating the protein’s surface-exposed aliphatic side-chains, loading its initial energy into the macromolecular structure. In stark contrast, the longer distance configurations invert this relationship, becoming negative loading profiles that describe a ground-state depopulation mechanism driven by field-induced ground-state polarization. The 3.0 mm configuration blue-shifts its vibrational coordinate to a distinct bleach at 3244 cm^-1^, denoting the depopulation of a network of strongly hydrogen-bonded water molecules as the proton charge density transfers into the surrounding solvent.

Upon reaching the intermediate 14.7 mm distance sample, the carrier loading peak appears as a bleach at 2728 cm^-1^. The 2728 cm^-1^ mode is a signature of an Eigen-type protonated water cluster (H_9_O_4_^+^). The recurrence of the 2728 cm^-1^ bleach from the spectral profile of the 2.0 mm data in Figure 4a confirms that the wavepacket at 14.7 mm propagates within a highly organized cage of structural water molecules. This structural organization is visualized by the multi-dimensional mode coupling resolved in the 2D correlation maps shown in Supplementary Figure S6. The 2D correlations in Supplementary Figure S6f expose a synchronized relationship between the low-frequency 300–600 cm^-1^ solvent librational modes and the high-frequency 1900–2800 cm^-1^ hydronium bending + libration combination bands. The strong correlation linking these two groups highlights that the hydronium combination bands are vibrationally coupled to the low-frequency rocking modes of the surrounding water molecules. This, in addition to the absence of the 100–150 cm^-1^ coherent oscillations in the 14.7 mm distance sample (Supplementary Figure S5b) allows us to form a more defined picture of the interfacial solvent region and provides greater insight into the long-time phase coherence maintained in the sample dynamics. The vibrational coupling and the reduction of solvent low-frequency dephasing pathways permit the local solvent network to organize into a rigid, highly symmetric geometric solvation shell around the active interfacial region. This structural caging mechanically restricts the classical transition dipole moment from oscillating freely, suppressing the overall field-induced ground-state polarization response across the symmetric hydration shell. Incidentally, this electrodynamic interaction provides a physiochemical rationale for the notably attenuated spectrum that we previously observed in the 14.7 mm sample in Figure 4a when contrasted with the s-IFN-γ control sample. In summary, the 2728 cm^-1^ bleach reveals that the initial proton charge density begins from a structured, (field-induced) pre-organized solvent environment; and furthermore, this solvent structural encapsulation provides a basis for the underlying physical mechanism that sustains its long time-scale phase coherence when compared with s-IFN-γ in Figure 2.

In the 40.0 mm distance sample, the carrier loading peak blue-chirps back to a prominent bleach at 2975 cm^-1^. This shifting sequence (2975 cm^-1^ → 3244 cm^-1^→ 2728 cm^-1^→ 2975 cm^-1^) uncovers an adaptive frequency chirping mechanism that establishes a highly optimized quantum transport pathway across the spatial distance matrix. The carrier wavepacket shifts its fundamental frequency to align with the specific structural state of the local solvation environment dictated by each distance boundary. This sequence functions to maximize the quantum coherence of the carrier relative to the localized solvent-protein interface, optimizing the interfacial match to maintain long-range quantum coherence all the way to its 40.0 mm distance destination.

#### 2.6.2 PCA2 Analysis of the coherent nonlinear amplification pathway

The second principal component (PCA2) in Figure 4c displays the coherent nonlinear driver of the translating wavepacket. It maps the spectral vibrational modes through which the system executes multi-photon up-conversion and energy harvesting to counteract solvent dielectric damping. Spectroscopically, PCA2 displays as a positive loading profile across all spatial distances, showing a universal two vibrational-state quantum amplification pathway defined by a fundamental positive loading peak and a nonlinearly shifted first overtone. The systematic distance-dependence of these peak coordinates provides a direct probe of the local curvature and topological changes of the underlying potential energy surface.

##### 2.6.2.1 The 2.0 mm and 40.0 mm distances

At the 2.0 mm near-field boundary and the 40.0 mm far-field antinode, the nonlinear driver loading peaks converge with identical spectral modes, resolving a fundamental peak at 1960 cm^-1^ and a prominent overtone at 4020 cm^-1^. Comparing the measured overtone against the expected harmonic frequency (1960 × 2 = 3920 cm^-1^) reveals a notable anharmonic shift of +100 cm^-1^. This specific shift serves as a spectroscopic signature of a Zundel low-barrier double-minimum potential. As the central barrier separating the two minima is suppressed, the proton wavefunction delocalizes across a broad, flat potential region bounded by steep repulsive walls. The fact that both the 2.0 mm and 40.0 mm distance samples have vibrational modes with the same frequency and with equivalent amplitudes confirms that far-field spatial phase interference retains the same low-dephasing quantum environment that is activated inside the confined near-field solvent cage. This far-field conservation of the wavepacket shape is due to a macroscopic self-focusing mechanism. Through this mechanism, the propagating wavepacket undergoes spectral condensation, progressively narrowing its spectral bandwidth (FWHM) from 1375.98 cm^-1^ at the 2.0 mm near-field, through 1263.70 cm^-1^ at the intermediate node (14.7 mm), down to a minimum of 1228.44 cm^-1^ at the 40.0 mm far-field distance. This reduction in the wavepacket spectral entropy over distance demonstrates how the system compresses its phase space to sustain long-range quantum coherence.

##### 2.6.2.2 The 3.0 mm relaxation

At a distance of 3.0 mm, the potential energy surface shifts dynamically due to the field-induced structural distortions across the interface. At this spatial coordinate, the driver loading peaks red-shift to 1912 cm^-1^ and 4000 cm^-1^. This spectral shift reflects geometric relaxation and a concurrent lowering of the activation energy barrier, driven by the field-induced outward distortion of the donor group toward the solvent environment. This structural alteration mechanically guides the reactant closer to its transition state, establishing a low-barrier potential well. Rather than undergoing spectral broadening, the 3.0 mm profile exhibits the highest peak amplitude across the entire distance matrix. This behavior confirms that the potential energy surface is bounded by steep repulsive walls; these walls tightly confine the system and concentrate the transition intensity within two sharply defined vibrational states to drive a highly focused, adiabatic quantum transport pathway.

##### 2.6.2.3 The 14.7 mm confinement and self-focusing gateway

Conversely, upon entering the 14.7 mm spatial phase interference node, the nonlinear driver profile blue-shifts to 2050 cm^-1^ and 4217 cm^-1^ while also undergoing significant spectral broadening and amplitude reduction. This increase in the effective force constant and subsequent broadening provide experimental verification of the structural confinement imposed along the potential energy surface by the surrounding solvent environment. As previously established, the low-frequency 100–150 cm^-1^ solvent fluctuations are suppressed at this spatial phase interference node, subsequently supporting the formation of a highly ordered, symmetric solvent cage that encloses the propagating wavepacket. As the wavepacket couples to the rigid network of H-bonds comprising this structural water cage, the underlying potential well steepens. This increased confinement forces the core driver to blue-shift up to 2050 cm^-1^ to match the increased vibrational force constant of the confined interface. Concurrently, the pronounced spectral broadening of both vibrational bands reflects a vibrational mode filtering (spectral cooling) process, characterizing the phase-matched vibrational energy exchange between the Zundel wavepacket and the localized phonon modes of the structural water cage. The 14.7 mm dataset identifies the critical threshold triggering nonlinear self-focusing. Spatial isolation of the wavepacket within this rigid matrix induces self-phase modulation, enabling the wavepacket to modify the local dielectric environment and emerge into the far-field as a stable, self-refining soliton.

### 2.7 Distance-dependent Anharmonic Coupling and Vibronic Synchronization Across the Macroscopic Gap

To characterize the long-lived vibrational coherence of the wavepacket and resolve its asymptotic structural configuration. The 2D contour maps in Figure 5 reveal active vibrational coherences of the molecular ensemble spanning 10 cm^-1^ to 10,000 cm^-1^ within the 3.0 to 5.0 ps time window. In bulk liquid water at ambient conditions, coherent vibrations typically undergo rapid thermal dephasing on a 50–100 fs timescale. The persistence of these vibrational coherences up to 5.0 ps provides strong evidence for a topologically protected quantum state that is effectively decoupled from conventional environmental decoherence channels. To isolate the specific structural configurations governing the wavepacket trajectory during the energy exchange phase, we analyzed the fourth time constant derived from the UB-TDS phase measurements. This timescale corresponds to the intermolecular energy transfer between the propagating wavepacket and the protein-solvent interface (Figure 5e). Remarkably, at this specific stage, the control sample and all four macroscopic separation distances synchronize temporally onto a common convergence point at ∼3.3 ps. In classical solution-phase biophysics, disparate geometric boundary conditions naturally dictate decoupled relaxation rates. The global temporal synchronization observed here suggests that the entire protein macromolecule and its surrounding solvation layer function as a unified macroscopic quantum waveguide across the protein–protein separation distance. This synchronized temporal junction characterizes the phase where the traveling wavepacket, passing through its ballistic propagation loop, undergoes coordinated vibronic energy redistribution. The system undergoes highly directional intermolecular vibrational energy transfer from the delocalized, translating Zundel-type charge carrier into the localized modes of both the surrounding protein side chains and the adjacent interfacial water network, thereby driving the non-adiabatic population transfer required to sustain the quantum transport cycle.

**Figure 5.**
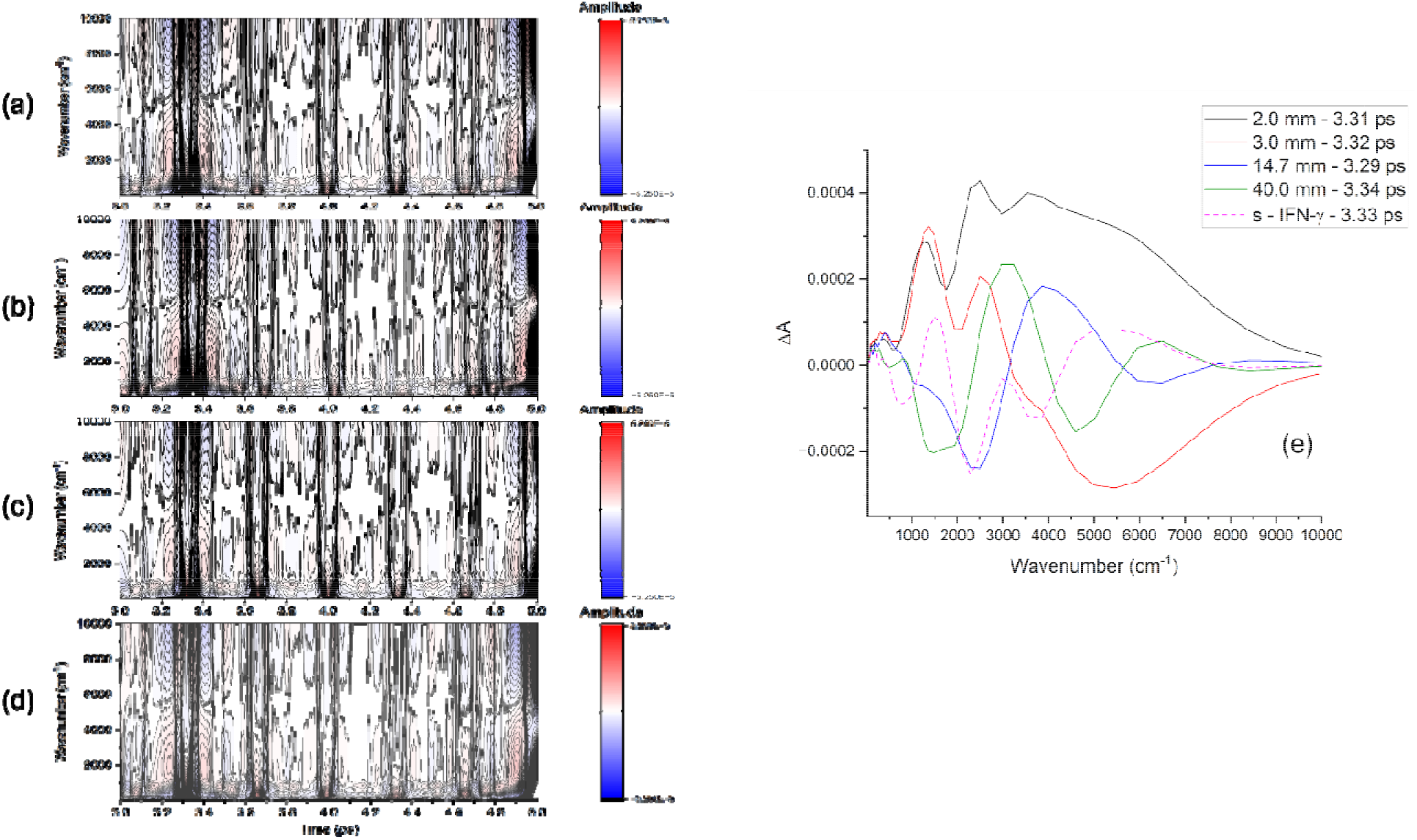
Late transient state phase maps and the fourth time constant spectra mapping intermolecular energy transfer **(a–d)** 2-D time-frequency spectral contour maps displaying the phase spectra of the UB-TDS data within the late transient time interval from 3.0 ps to 5.0 ps for **(a)** the 2.0 mm distance sample, **(b)** the 3.0 mm distance sample, **(c)** the 14.7 mm distance sample, and **(d)** the 40.0 mm distance sample. The matrices have been smoothed using a 2-D 50-point Savitzky–Golay digital spectral filter. The contour features trace the late-stage spatial organization and phase profiles of the wavepacket across the spatial distances. **(e)** Experimentally resolved transient phase spectra corresponding directly to the fourth time constant extracted from EMD across the 10 to 10,000 cm⁻¹ spectral range. This temporal component is associated with intermolecular energy transfer between the propagating wavepacket and the protein-solvent interface. Traces show the 2.0 mm distance sample (black solid line, τ_4_ = 3.31 ps), the 3.0 mm distance sample (red solid line, τ_4_ = 3.32 ps), the 14.7 mm distance sample (blue solid line, τ_4_ = 3.29 ps), the 40.0 mm distance sample (green solid line, τ_4_ = 3.34 ps), and s – IFN-γ (magenta dotted line, τ_4_ = 3.33 ps).

#### 2.7.1 The single protein field-free control (s - IFN-γ)

The IFN-γ control spectrum in Figure 5e resolves a broad, shallow bleach centered at 743 cm^-1^ alongside an absorbance centered at 1487 cm^-1^. The mid-infrared region resolves distinct GSBs at 2294 cm^-1^ and 3858 cm^-1^, which operate in tandem with a broad, high-energy absorbance centered at 5000 cm^-1^. The 2294 cm^-1^ bleach constitutes the first overtone (second harmonic) of the (unconstrained) surface-exposed Zundel stretch (1147 cm^-1^ x 2 = 2294 cm^-1^), while the 3858 cm^-1^ mode represents an intrinsic protein bending and solvent stretching combination state. The appearance of the 2294 cm^-1^ and 3858 cm^-1^ as GSBs yields a profound structural conclusion: they demonstrate that the control protein inherently possesses the pre-synchronized transport coordinates and coupled macromolecular framework required to initiate waveguide dynamics. However, in the absence of an external inter-protein electric field to stabilize these pathways, the system undergoes an immediate energy exchange with the *environment*. Through intense off-diagonal anharmonic coupling, the population captured by the 2294 cm^-1^ and 3858 cm^-1^ ground states is rapidly depleted. Instead of being channeled into a non-dispersive waveguide, this energy is stochastically down-converted and transferred into the bulk solvent, driving the emergence of the broad 5000 cm^-1^ bending-stretching combination absorbance. This spectral routing explicitly confirms the termination of the quantum transport propagation pathway via isotropic thermal dissipation into the bulk environment.

#### 2.7.2 The 2.0 mm near-field sample

The 2.0 mm distance spectrum resolves a broad continuum absorbance spanning from 10 to 10,000 cm^-1^, the definitive spectroscopic signature of a highly delocalized Zundel proton transport network. Superimposed on this continuum are three GSBs centered at 625 cm^-1^, 1768 cm^-1^, and 2975 cm^-1^. This unique profile reveals a mechanism wherein the propagating wavepacket actively drains population from specific localized coordinates to sustain the delocalized continuum state. Specifically, the system depopulates the collective intermolecular librational rocking modes of the interfacial water network (625 cm^-1^), the internal bending vibration of the protonated water core (1768 cm^-1^), and the collective modes of surface-exposed aliphatic side chains (2975 cm^-1^). This synchronized depopulation confirms that the global protein-solvent interface functions as a coherent, macroscopic quantum waveguide that harvests energy directly from its core protonated coordinates to drive the quantum transport cycle.

#### 2.7.3 The 3.0 mm intermediate distance sample

In stark contrast, the 3.0 mm distance sample maps a highly specialized, asymmetric energy-exchange profile that reflects a narrow, low-barrier potential well. In this case, the absorbance continuum is truncated, extending only from 10 to 3000 cm^-1^ and is overlaid with distinct GSBs. The first bleach resolves as a blue-shifted interfacial water mode centered at 682 cm^-1^, denoting that the outward distortion of the donor group compresses and stiffens the local H-bonding network. This structural constraint forces the adjacent water molecules into a tightly bounded network, steepening the local force constant, leading to a blue-shift of the fundamental librational mode frequency relative to the 2.0 mm near-field baseline. This is accompanied by a bleach of the core vibrational mode of the propagating wavepacket at 2103 cm^-1^ that reflects a direct population depletion across the hydronium combination band. This feature represents the dynamic blue-shift and depopulation of the fundamental nonlinear driver mode (resolved at 1912 cm^-1^ in PCA2) as the local force constant rapidly spikes during the active energy transfer phase. The higher frequency region of the spectrum is dominated by a broad GSB that spans from 3100 to 8500 cm^-1^. Within this broad bleach there is a barely perceptible minimum at 3538 cm^-1^ and a deep minimum at 5450 cm^-1^. Since this broad bleach feature directly overlaps with the IVCT band, its conversion into an intense bleach confirms that the low-barrier, vibronically adiabatic pathway operates by systematically depopulating this pre-polarized electronic transition to continuously pump energy back into the propagating wavepacket.

#### 2.7.4 The 14.7 mm intermediate node sample

The spectrum of the 14.7 mm distance sample features a broad GSB extending from 930 to 3000 cm^-1^ with a deep minimum centered at 2470 cm^-1^. In aqueous molecular spectroscopy, the 2470 cm^-1^ coordinate serves as the distinctive signature of an Eigen-type (H₉O₄⁺) hydronium stretching (*22*) environment. Within this deep Eigen-type bleach is a broad, non-localized weaker bleach centered at 1051 cm^-1^, which serves as the spectroscopic marker for a delocalized Zundel core environment. Concurrently, the low-frequency region of the spectrum reveals a broad absorbance extending from 200 – 800 cm^-1^ with a central, prominent peak at 405 cm^-1^ and this is accompanied by a broad absorbance band spanning from 3200 to 5000 cm^-1^ and centered at 3858 cm^-1^ in the mid-infrared region of the spectrum. This feature reflects the direct activation of a strongly coupled combination band (2236 + 1622 = 3858 cm^-1^), which unifies a strongly red-shifted, Zundel-type H-bonded O–H stretch with the core water bending vibration. And finally, the near-infrared region of the spectrum depicts a broad, shallow GSB spanning from 5500 to 7200 cm^-1^ and centered precisely at 6488 cm^-1^. The 6488 cm⁻¹ coordinate represents the fourth harmonic overtone (4 × 1622 cm⁻¹) of the collective water bending mode. These simultaneous features in the experimental spectrum capture a dynamic coexistence where the two limiting geometries of the hydrated proton integrate into a single, united structure to drive a phase-coherent, dual-frequency propagating wavepacket across the interface.

Perhaps, most curious is the 6488 cm^-1^ overtone mode. The 6488 cm^-1^ bleach marks the fourth harmonic (4 x 1622 cm^-1^ = 6488 cm^-1^) of the local water bending mode. One would normally expect anharmonicity to red-shift the energy of the 4^th^ harmonic of the overtone to lower energies. But in this case, the mode behaves as a linear harmonic, which consequently, provides some insight into the nature of the dynamics of the water molecules in the interfacial region of the protein. The precise frequency of the overtone mode discloses that the potential energy well of the H-O-H bend undergoes a field-induced potential well distortion due to the protein emitted field and the consequent Eigen cage environment of the propagating wavepacket. This structural confinement stiffens the localized bond coordinates, counterbalancing the bulk-like anharmonicity to match this precise integer multiple. Accordingly, the system utilizes this populated fourth harmonic state to actively generate and “fuel” the low-frequency lattice mode via quantum parametric resonance. In this mechanism, intermolecular vibrational energy transfer is driven by a high-order nonlinear coupling channel (*23*, *24*) at a near-perfect 16:1 integer frequency ratio, where the 6488 cm^-1^ harmonic transfers energy directly to the coherent acoustic phonon mode peaking at 405 cm^-1^ (405 x 16 = 6480 cm^-1^). The combination of active modes forms a transport mechanism: within the structured solvent environment, the newly formed 405 cm^-1^ acoustic phonon acts as a physical lattice that encapsulates and directly shields the 3858 cm^-1^ propagating wave core from random solvent thermal fluctuations. Since a non-dispersive soliton cannot exist in a purely linear system, this cross-mode nonlinear coupling drives a continuous process of anharmonic vibrational energy pooling between the low-frequency acoustic shield and the 3858 cm^-1^ wave core, effectively removing boundary drag to prevent vibrational energy relaxation into the bulk solvent. This integrated framework is actively required to both power and stabilize the solitonic wave to preserve its anomalous long-lived vibrational coherence.

#### 2.7.5 The 40.0 mm far-field destination sample

The spectral profile of the 40.0 mm distance sample resolves three distinct GSBs centered at 442 cm^-1^, 1422 cm^-1^, and 4588 cm^-1^ along with two prominent absorbance bands centered at 3100 cm^-1^ and 6488 cm^-1^. The GSBs reflect a single, hybrid quantum state arising from anharmonic mixing between an inner Zundel core and a librational network of peripheral water molecules forming a structural shell within the interfacial layer. Within this delocalized system, three distinct spectral features are anharmonically coupled as constituent components of the same ground state: the low-frequency peripheral water libration at 442 cm^-1^, the Zundel shared-proton bending deformation (*25*) at 1422 cm^-1^, and the mid-infrared fourth harmonic overtone of the Zundel core environment at 4588 cm^-1^ (4 × 1147 cm^-1^). The active dynamical role of this peripheral network is to induce anharmonic mode mixing, which mediates the intermolecular vibrational energy transfer directly to the traveling wavepacket. As the flanking solvent shell performs its low-frequency 442 cm^-1^ out-of-plane librational/rocking motion, its dynamic geometric displacement introduces strong cubic and quartic anharmonic terms into the potential energy surface of the Zundel core, causing the wavefunctions of the central proton and the surrounding cage to hybridize via the shared hydrogen bonds. This cage-driven distortion modifies the local potential well, lowering the potential energy barrier for proton transfer and breaking local inversion symmetry. The resulting relaxation of conventional parity selection rules elevates the nonlinear transition dipole moment gradient, allowing the energy from these three GSBs to couple constructively into the broad near-infrared vibrational state centered at 6488 cm^-1^, which represents the fourth harmonic overtone (4 × 1622 cm^-1^) of the collective water bending mode. The conversion of this near-infrared transition into a prominent absorbance signifies a complete structural inversion of the 14.7 mm intermediate node spectral profile, demonstrating an elegant macroscopic quantum reciprocity across the macroscopic gap. While the 14.7 mm node utilizes the 6488 cm^-1^ overtone as an energy source to generate the 405 cm^-1^ acoustic structural shield, the 40.0 mm destination utilizes this vibrational coordinate as an energy sink, where the arriving wavepacket deposits its remaining energy directly back into this higher energy vibrational state. Simultaneously, the mid-infrared region populates a broad absorbance continuum centered at 3100 cm⁻^1^, reflecting the active population of the highly polarized flanking water molecules of the Zundel core. Once populated via this multi-quantum coherent transition, the cross-mode anharmonic coupling of the 6488 cm^-1^ state establishes a nonlinear resonant cross-coupling that channels energy downward to sustain this broad 3100 cm^-1^ stretching continuum. The precise resolution of these discrete, higher-order fourth harmonics at this 3.3 ps time constant across a macroscopic distance provides direct spectroscopic confirmation that the interface has entered a macroscopically ordered quantum state. In a classical, disordered bulk solvent, these ultra-weak multi-quantum transitions would be severely degraded and rendered incoherent by rapid thermal dephasing. Their prominent, sharp visualization in the experimental spectrum confirms that the protein emitted-field organizes the interfacial layer into a cooperative network. This establishes a continuous coherent self-stabilization mechanism that allows the arriving wave core and its peripheral solvation cage to maintain absolute phase coherence and suppress classical thermal noise. This nonlinear multi-quantum resonance suppresses randomized energy dissipation, enabling the system to guide the propagating solitonic wavepacket along the macroscopic proton transport pathway without thermal dispersion.

Moreover, we have identified that the fundamental 1622 cm^-1^ water bending vibration functions as the central vibronic coordinating mode for the entire interfacial region. This bending vibration acts as a macroscopically coherent collective mode, where its period establishes the fundamental timing frequency that dynamically synchronizes every active vibrational coordinate across a macroscopic distance. Through field-driven anharmonic coupling, the low-frequency lattice waves, the core wave transitions, and the higher-energy near-infrared overtones exhibit harmonic commensurability with this 1622 cm^-1^ base frequency. Consequently, the entire proton transport pathway operates as a matrix of harmonically synchronized vibrational states. This precise frequency relationship ensures that the overlapping, multi-frequency time-dependent wavefunctions share a common fundamental period, maintaining continuous constructive phase interference across the interface. Constrained by this harmonic integer alignment, the synchronized matrix establishes direct multi-quantum resonances that channel energy across these distinct stretching, bending, and lattice states with minimal phase velocity dispersion or inelastic scattering into the surrounding medium. This global phase synchronization of the multi-frequency eigenstates effectively suppresses classical thermal dephasing noise, stabilizing and shielding the traveling solitonic wavepacket across the macroscopic gap.

### 2.8 Active Bimodal Soliton Release, Vibronic Coupling, and Interfacial Lattice Organization

The fifth and final time constant resolves the active release of the soliton across the coupled protein-solvent interface. Within the active, field-modulated configurations, the excess energy of the wave core is redistributed across the interfacial network via intense off-diagonal anharmonic coupling. Specifically, the interfacial solvent undergoes a structural phase transition that modifies the topography of the local potential energy landscape. Consequently, the precise molecular topology of this field-driven solvent transition dictates the transport dynamics and subsequent efficiency of the non-dispersive soliton release.

#### 2.8.1 Control sample

In distinct contrast to the active, field-modulated samples, the standalone s – IFN-γ control sample resolves a relaxation pathway dominated by vibrational dephasing via solvent friction. The experimental spectrum of s – IFN-γ (Figure 6a) features a broad bleach extending from approximately 185 cm^-1^ to 970 cm^-1^. This extended envelope encompasses both the intermolecular solvent hydrogen-bond stretch and the bulk-like water libration, resolving a local minimum at 573 cm^-1^. Additionally, the spectrum exhibits a distinct GSB centered at 2103 cm^-1^, which we attribute to the anharmonic first overtone (second harmonic) of the 1051 cm^-1^ Zundel stretching mode, alongside a weak bleach centered at 3858 cm^-1^ (1622 cm^-1^ bend + 2236 cm^-1^ stretch combination mode). The coexistence of the intrinsic Zundel signature and the 1622 cm^-1^ IFN-γ bend frequency yields a remarkable biophysical conclusion: it demonstrates that the control protein inherently possesses the necessary structural framework to support bimodal soliton transport dynamics. Consequently, a distance-dependent, inter-protein radiative interaction is the singular macroscopic trigger required to activate these non-dispersive waveguide dynamics. The control spectrum also resolves a narrow, low-frequency absorbance centered at 110 cm^-1^, which is associated with localized solvent hydrogen-bonding interactions with the polar side chains (*26*) of the protein. Furthermore, two prominent absorbance bands emerge at 1364 cm^-1^ and 2728 cm^-1^. The 1364 cm^-1^ mode is assigned to an intrinsic tryptophan indole ring vibration, reflecting the structural relaxation of the residue back to its inactive ground state. Meanwhile, the 2728 cm^-1^ absorbance serves as a distinct spectroscopic signature of an isolated, Eigen-type protonated water cluster (H_9_O_4_^+^), reflecting the localized asymmetric stretching of the hydronium core. Taken together, these absorbances describe a localized regime dominated by stochastic structural fluctuations. In the absence of external field coordination, these uncoupled modes fail to support long-range phase coherence, effectively prohibiting subsequent soliton release into the bulk medium.

**Figure 6.**
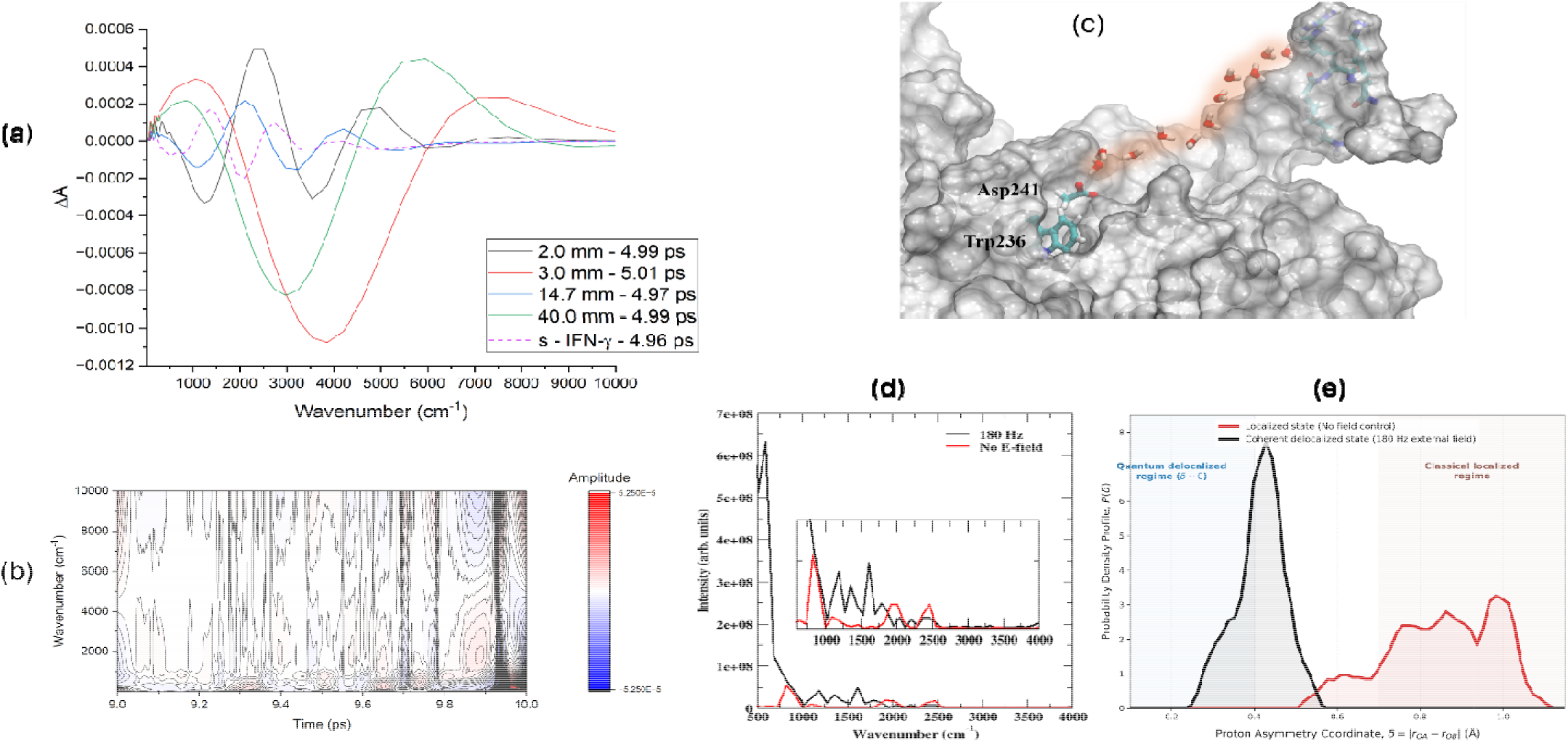
Long-lived far-field phase-coherence, ballistic soliton release, and atomistic QM/MM quantum-delocalized proton transfer mechanisms **(a**) Experimentally resolved transient phase spectra corresponding to the fifth and final time constant extracted from EMD. This terminal temporal component maps the phase-coherent ballistic release of the wavepacket into the extended bulk environment. The 2.0 mm distance sample is shown with a black solid line (τ_5_ = 4.99 ps), the 3.0 mm distance with a red solid line (τ_5_ = 5.01 ps), the 14.7 mm distance with a blue solid line (τ_5_ = 4.97 ps), the 40.0 mm distance sample with a green solid line (τ_5_ = 4.99 ps), and s – IFN-γ with a magenta dotted line (τ_5_ = 4.96 ps). **(b)** The 2-D time-frequency spectral contour map displaying the phase spectrum of the 40.0 mm distance sample within the terminal 9.0 ps to 10.0 ps time window, smoothed using a 2-D 50-point Savitzky–Golay digital spectral filter. **(c)** Molecular surface representation of the IFN-γ homodimer along with the functional catalytic residues Trp236, Asp241, and an intervening 11-molecule water wire illustrated in a licorice model to denote the chemically active subsystem treated quantum mechanically, embedded within the classically modeled protein structure. **(d)** Infrared absorption spectrum computed from the Fourier transform of the quantum-mechanical total dipole autocorrelation function (DACF) tracked over the 350 fs QM/MM trajectories. The control protein (no electric field) is shown with a red trace, while the field-driven trajectory (180 Hz) is shown with a black trace. **(e)** Statistical probability density distribution of the proton asymmetry parameter computed across the 11-molecule water wire network under resting (red line) and field-driven (black line) regimes.

#### 2.8.2 The 2.0 mm field environment

At a separation distance of 2.0 mm, we identify a localized structural phase transition within the interfacial solvent network that fundamentally reshapes the topography of the potential energy landscape governing the propagating wavepacket. The time-resolved spectrum resolves two broad GSBs centered at 1250 cm^-1^ and 3858 cm^-1^. We ascribe the 1250 cm^-1^ mode to a delocalized Zundel-type continuum, while the 3858 cm^-1^ feature is assigned to a combination mode comprising the 1622 cm^-1^ protein bending vibration and a 2236 cm^-1^ heavily polarized Zundel-type interfacial solvent stretch. The concurrent bleaching of the 1250 cm^-1^ and 3858 cm^-1^ states demonstrates that the flat, symmetric PES characteristic of the initial Zundel framework undergoes a rapid topographical transition. It shifts dynamically into a deep, asymmetric potential energy well localized at the Trp-Asp donor-residue complex within the first coordinating solvation shell. Concurrently, the experimental spectrum resolves two prominent emerging absorbances centered at 2400 cm^-1^ and 4800 cm^-1^. We associate the 2400 cm^-1^ absorbance to an asymmetric proton-stretching transition localized within a short, strong hydrogen bond (SSHB) directly adjacent to the proton donor group. This feature signifies that the surrounding solvent molecules cluster around the excess charge to yield a highly localized, distorted Eigen-type asymmetric proton complex. The interplay between these coordinate GSBs and the emerging absorbances marks a radical alteration of the interfacial PES. Confining the translation-invariant wave core within this deep potential well triggers an instantaneous, high-intensity surge in localized charge separation along the SSHBs. Taken together, these spectral signatures support the premise that soliton release is driven by short-distance propagation coupled intensely with the localized dynamics of the protein-interfacial boundary. This electronic and vibrational surge is amplified by the collective dynamics of the protein, clarifying the physical nature of the 2975 cm^-1^ PCA1 FFT mode (Figure 4b). In multivariate statistical terms, PCA1 captures the dominant collective motion of the entire system, mapping directly onto the collective, hydrophobic aliphatic C–H side-chain fluctuations of the protein core. The 2975 cm^-1^ PCA1 loading resolves as a net absorbance at this distance, whereas it displays as a clear GSB across all other separation intervals. This sign inversion indicates a 180-degree phase reciprocity, wherein the radiative near-field boundary conditions induce a localized structural and mechanical compression of the hydrophobic core. This core compression reverberates outward, mechanically condensing the local boundary water layers into a highly constrained geometry that directly drives the emerging 2400 cm^-1^ absorbance state. The intense vibrational energy transferred via this in-phase mechanical coupling excites a coherent response within the localized interface, resolving as two narrow low-frequency absorbance bands centered at 185 cm^-1^ and 371 cm^-1^ (Supplementary Figure S8). We attribute the 185 cm^-1^ mode to the collective intermolecular H-bond stretching vibration of the immediate solvation layer, while the narrow 371 cm^-1^ mode represents the discrete intermolecular stretching vibration between the localized hydronium ion and the Asp carboxylate oxygen. These low-frequency modes serve as precise structural metrics, demonstrating that the first hydration shell, directly interacting with the Trp-Asp complex organizes into a highly ordered, structurally uniform lattice bound by a rigid hydrogen-bonding network. This ordered surface hydration layer creates a sharp physical boundary that structurally constrains the phase-coherent dynamics driven by the 3858 cm^-1^ (1622 cm^-1^ + 2236 cm^-1^) IFN-γ bending mode within the immediate first hydration shell. This constraint keeps the wavepacket energy directed along the primary reaction coordinate, thereby mechanically insulating the transport pathway from the random thermal fluctuations of the surrounding solvent bath. Further, the presence of the 3858 cm^-1^ GSB implies that a specific, pre-synchronized population of water molecules was actively present prior to the soliton release event. Since this specific population is efficiently transferred out of the ground state and into higher-energy absorbance states, the IFN-γ bending mode dictates the trajectory of the wavepacket at the interface. This ensures that the high-power charge surge routes exclusively via intense off-diagonal anharmonic coupling directly into the 4800 cm^-1^ bending-stretching combination band. This high-energy vibrational mode holds the charge-separation state at the Trp-Asp interface, providing the localized work required to drive the protein’s structural rearrangements. Ultimately, this high-energy pathway promotes a highly localized vibrational excitation of the immediate interfacial water layer, completing its trajectory within the phase-coherent first hydration shell bound by the 185 cm^-1^ / 371 cm^-1^ intermolecular lattice boundary.

#### 2.8.3 The 3.0 mm field environment

As the inter-protein separation distance shifts to the 3.0 mm spatial coordinate, the time-resolved spectrum captures the active macroscopic release of a high-energy, non-dispersive soliton across the coupled protein-solvent interface. This nonlinear transport state is defined by a broad GSB centered at 3858 cm^-1^ in coordination with a broad near-infrared absorbance at 7075 cm^-1^, a continuous absorbance envelope spanning the far-to-mid-infrared region from 20 to 1750 cm^-1^ (centered at 1050 cm^-1^), and narrow low-frequency lattice resonances at 101 cm^-1^ and 202 cm^-1^. The restriction of the ground-state depletion to a singular 3858 cm^-1^ GSB establishes a strict, conservation-compliant energy pathway governing this spatial coordinate. Since no other distinct bleaches are resolved in the spectrum, the 3858 cm^-1^ pre-synchronized coupled protein-solvent state serves as the exclusive quantum-mechanical reservoir and upstream driver fueling the entire nonlinear transport event. As the non-dispersive soliton completes its trajectory, this pre-synchronized population undergoes a localized structural transition representing a non-thermal structural melting of the boundary hydration layer. This phase transformation is inferred by the concurrent emergence of the 7075 cm^-1^ and 1050 cm^-1^ (continuum) absorbances, which reflect a rapid flattening of the local potential energy surface and the subsequent collapse of the steep angular and torsional potential barriers - the directional coordinates that sustain a highly correlated, geometric alignment across the ground-state 3858 cm^-1^ population. Overcoming these directional barriers flattens the anisotropic potential wells of the initial H-bonded framework, splitting the population into two co-emergent structural environments. A sub-population of these water molecules evolves into a vibrationally decoupled, monomeric configuration where the uncoupled O-H oscillators of these isolated monomers drive the 7075 cm^-1^ near-infrared absorbance. The 7075 cm^-1^ mode is the first vibrational overtone combination (ν_1_ + ν_3_) that denotes the structural wake left behind by the passing wave core. Simultaneously, the remaining portion of the energy from the 3858 cm^-1^ baseline population is partitioned into the absorbance continuum centered at 1050 cm^-1^. The distinctive shape and breadth of this envelope provide a clear spectroscopic marker for an extended Zundel-type waveguide. This delocalized continuum functions as a non-thermal electronic-vibrational dissipation buffer, rapidly redistributing the high-frequency charge across a translation-invariant continuum of highly mobile, shared-proton topologies, thereby preventing thermal dephasing at the interface. Rather than allowing this energy to diffuse stochastically into the bulk medium, the lower region of this continuous spectral envelope is explicitly dominated by a highly ordered, low-frequency acoustic hydration lattice resolving at 101 cm^-1^ (the fundamental lattice mode) and 202 cm^-1^ (its discrete harmonic overtone). This coherently phase-coupled harmonic system exhibits a 1:2 frequency ratio (Supplementary Figure S8), demonstrating a quantum-mechanical resonance that actively enforces macroscopic phase coherence across the interfacial region. The synchronized behavior of this acoustic lattice establishes a non-thermal waveguide that dynamically shapes the local potential energy surface. This framework provides the necessary topological confinement for directing the generated phonon mode (101/202 cm^-1^), focusing the high-power charge surge exclusively along a one-dimensional reaction coordinate and preventing the high-energy soliton from scattering laterally into the bulk solvent.

#### 2.8.4 The 14.7 mm field environment

At the 14.7 mm distance coordinate, the spectrum illustrates the active release of a highly mobile, ballistic soliton wavepacket where the excess charge carrier is launched as a structurally intact, delocalized Zundel core (H_5_O_2_^+^) along the interface. This nonlinear transport state is characterized by a dual GSB profile featuring a broad continuum centered at 1051 cm^-1^ and a structured solvent stretch centered at 3244 cm^-1^. In strict coordination with these bleaches are two broad higher-energy absorbances centered at 2102 cm^-1^ and 4207 cm^-1^, alongside narrow, low-frequency lattice resonances at 110 cm^-1^ and 221 cm^-1^ (Supplementary Figure S8). The dual-GSB architecture establishes that the baseline population prior to wavepacket release is natively composed of a highly correlated Zundel-type continuum (1051 cm^-1^), mapping the core asymmetric (O···H^+^···O) proton-transfer coordinate (ν_as_) embedded within a structured outer-hydration shell network centered at 3244 cm^-1^. Importantly, the 3858 cm^-1^ coupled protein-solvent mode, which resolved as a prominent absorbance in the immediately preceding time constant, is completely absent in the current spectrum (Figure 6a). This spectral evolution directly follows the complete population decay of this transient quantum-mechanical reservoir, where the energy harvested from the relaxation of this 3858 cm^-1^ state serves as the exclusive upstream driver fueling the ballistic transport event. As the high-energy wavepacket is released, this decaying population undergoes a resonant, non-adiabatic energy transfer into the interface, appearing as the broad 2102 cm^-1^ and 4207 cm^-1^ absorbances. These active features represent the first overtone (second harmonic, *v* = 2) and third overtone (fourth harmonic, *v* = 4) of the 1051 cm^-1^ fundamental Zundel transition, respectively (1051 × 2 = 2102 cm^-1^ and 1051 × 4 = 4204 ≈ 4207 cm^-1^). This precise integer alignment confirms that the decaying population has relocated vertically to drive the ballistic transport of the delocalized charge core. Governing this entire multi-quantum process is an intense off-diagonal anharmonic coupling, which allows the high-frequency electronic and vibrational energy to accumulate as a stable, long-lived population within these collective vibrational states. This coupling effectively suppresses classical bulk-water thermal dephasing, preventing the traveling wave core from losing energy as random local heat. This highly energized, propagating Zundel waveguide is dynamically stabilized by the low-frequency acoustic hydration lattice resolving at 110 cm^-1^ (the fundamental lattice mode) and 221 cm^-1^ (its discrete harmonic overtone). This coherently phase-coupled harmonic system exhibits a precise 1:2 frequency ratio, denoting that a quantum-mechanical resonance actively maintains macroscopic phase coherence across the interface. The blue-shift of this acoustic lattice relative to the 3.0 mm coordinate (101/202 cm^-1^) reflects a localized geometric compression and mechanical stiffening of the boundary water layers. This compressed harmonic framework provides the necessary topological confinement to direct the generated phonon mode at 110/221 cm^-1^, focusing the energized proton transport along a stable, non-thermal one-dimensional waveguide, ensuring that the soliton’s forward momentum is preserved without undergoing spatial expansion or radiative decay into uncoupled bulk degrees of freedom.

#### 2.8.5 The 40.0 mm field environment

Evaluating the long-range far-field profiles reveals that these low-frequency terahertz (THz) modes are fundamentally important for sustaining the long-range transport of the soliton, acting as an emergent structural property of the coupled protein-solvent interface. For the 40.0 mm distance sample, the 5.0 ps time constant does not mark a terminal relaxation or structural end; instead, it captures an ongoing, active transport phase that continues its non-dispersive propagation out to a 10.0 ps temporal limit. This long-range, non-equilibrium transport phase (Figure 6a) is defined first by a restricted ground-state profile, resolving as a broad GSB centered at 2975 cm^-1^. Additionally, there is a broad absorbance band centered at 5950 cm^-1^. In the lower frequency region of the spectrum, there is also broad Zundel continuum absorbance extending from the lowest recorded frequency up to about 1370 cm^-1^, within which are embedded two narrow absorbance bands centered at 110 cm^-1^ and 221 cm^-1^ (Supplementary Figure S8). The restriction of the ground-state depletion to the 2975 cm^-1^ bleach highlights the core structural axis of the long-range transport phase, tracing the direct, systematic depletion of the collective, hydrophobic aliphatic C–H side-chain fluctuations on the protein surface. We note that the 2975 cm^-1^ mode serves as a primary feature in the principal component loading profile (PCA1) for the 40.0 mm distance sample (Figure 4b), verifying that the fast proton wave core inside the Zundel environment and these collective hydrophobic side-chain fluctuations are oscillating in phase synchronization as a bimodal soliton. The evolution of this initial 2975 cm^-1^ baseline population corresponds directly to the transiently generated 5950 cm^-1^ state. Through off-diagonal anharmonic coupling, the population undergoes a vertical multi-quantum state transfer, driving the prominent emergence of the 5950 cm^-1^ absorbance, which arises as the first overtone (second harmonic) of the 2975 cm⁻¹ fundamental transition (2975 x 2 = 5950 cm^-1^). This up-converted energy propagates as a coherent, non-dispersive wave packet, matching the velocity of the underlying Zundel continuum to transit to the interface without dephasing into the bulk solvent. In concert, the un-damped lattice phonons at 110 cm^-1^ and 221 cm^-1^ operate directly to stabilize the phase coherence of this high-energy soliton. This coherently phase-coupled harmonic system exhibits a strict 1:2 frequency ratio (110 x 2 = 220 ≈ 221 cm^-1^), signifying a quantum-mechanical resonance that actively maintains macroscopic phase coherence across the interface. This synchronized acoustic framework creates topological confinement to direct the wavepacket along a one-dimensional reaction coordinate, effectively preventing the traveling soliton from undergoing lateral dissipative scattering into the bulk solvent. The greater physical meaning of these coupled modes is revealed by the fact that the active transport dynamics survive and persist entirely out to 10 ps before concluding (Figure 6b), matching a non-equilibrium lifespan that significantly exceeds standard condensed-matter relaxation windows. In unperturbed aqueous environments, intermolecular phase coherence naturally dephases within a sub-picosecond regime due to rapid, stochastic thermal fluctuations in the solvent network. While the phase-coherent dynamics of the shorter field distance configurations conclude near the 5.0 ps time limit, the 2D experimental contour map captured for the 40.0 mm distance sample provides spectroscopic verification of phase coherence that is sustained into the 9–10 ps experimental window. The vertical coherence lines resolved in the spectral map (Figure 6b) demonstrate that the long-distance field modulation has structured a stable, periodic solvent lattice. This driven framework successfully shields the traveling bimodal soliton from standard bulk thermal dephasing, allowing the wave core to preserve its structural shape and amplitude over macroscopic timelines before the underlying Zundel depletions finally conclude and transduce back into localized resting modes at 10 ps to complete the 40.0 mm distance configuration timeline.

### 2.9 QM/MM computational characterization of field-induced solvent delocalization

To investigate the atomistic solvent mechanisms corresponding to the initial dielectric relaxation and sub-picosecond molecular rearrangements observed at the experimental first time constant (Figure 2e), hybrid quantum mechanics/molecular mechanics (QM/MM) molecular dynamics simulations were conducted, from each of which a 350 fs trajectory window was evaluated. In the QM/MM simulations, the active interfacial region comprising the dual proton-donor gating complex of residues Asp41 and Trp36, which correspond to Asp241 and Trp236 on the second monomer of the homodimer (Figure 6c) and an intersecting 11-molecule water wire connecting the proton donor group with the conserved KRKR motif in the C-terminal tail were treated explicitly within the quantum region. This sub-picosecond computational window is utilized to interpret the non-equilibrium structural, electronic, and optical state of the system prior to net proton transfer, mapping the landscape required to reach the initial Franck-Condon transition state. Trp236 (Trp36) was specifically included within the QM region to capture its electronic and structural dynamics, which act in tight cooperative alignment with the aspartate carboxylate coordinate to modulate the local electrostatic field gradient at the protein–solvent interface. The infrared absorption spectrum (Figure 6d), derived directly from the Fourier transform of the quantum-mechanical total dipole autocorrelation function (DACF), highlights a field-induced structural divergence that provides computational corroboration and additional insight into the experimental diabatic and adiabatic regimes initially uncovered in the ≤ 300 fs 2D spectral maps shown in Figures 2a–d and 2f. In the no-field control simulation (analogous to the experimental s–IFN-γ control sample), the optical response of the hydration layer is defined by a distinct librational mode peaking at 833 cm^-1^, alongside localized bending and stretching overtones resolved at 1960 cm^-1^ and 2450 cm^-1^ (Figure 6d). The presence of these discrete, high-frequency overtones and combination states provides a clear comparison to the experimental single-protein control in Figure 2e and indicates that in the computed baseline configuration, the non-equilibrium wavefunctions are localized within decoupled diabatic potential wells that support a non-adiabatic transport pathway.

We have chosen to apply an external field with a frequency of 180 Hz in the simulations. We conjecture that the 180 Hz temporal field oscillations represent a slow, macroscopic collective breathing mode of the IFN-γ homodimer that acts as an electro-mechanical envelope modulator. We experimentally observed that these structural fluctuations periodically deform the local dielectric environment, encoding a subtle but detectable 180 Hz undulation onto the high-frequency carrier field that becomes explicitly visible in the time-resolved phase measurements in the cavity configurations outlined in Section 2.1. Upon application of the external 180 Hz electric field, the unperturbed profile transitions into a highly cooperative quantum environment, establishing a broad spectral continuum (Figure 6d) that mirrors the signatures resolved in the experimental 40.0 mm distance sample in Figure 2e. Under field exposure, the low-frequency THz spectrum is dominated by a collective intermolecular mode centered at 85 cm^-1^ (Supplementary Figure S9), while the localized 833 cm^-1^, 1960 cm^-1^, and 2450 cm^-1^ states in the no-field spectrum are replaced by a red-shifted librational mode centered at 600 cm^-1^ and the activation of the broad Zundel proton transfer continuum spanning the ∼1000–1850 cm^-1^ mid-infrared region (Figure 6d). The field-driven shift of the 833 cm^-1^ population down to 600 cm^-1^ provides a clear computational analogue to the broad, low-frequency 884 cm^-1^ transient bleach resolved in the experimental UB-TDS phase measurements shown in Figure 2e, indicating that the field shifts the network away from its native resting frequencies by softening its torsional force constants. This low-frequency relaxation occurs in parallel with, but remains energetically distinct from, the activation of the broad, structureless Zundel continuum resolved at significantly higher frequencies in the spectrum. A comparative analysis of these profiles suggests that the emergence of this continuum absorption is a signature feature in the formation of the adiabatic pathway. The presence of the continuum indicates that the 180 Hz external field flattens the local electric field gradients, merging independent diabatic wells into a broad, translation-invariant wave channel.

To map the real-space geometric coordinates associated with this transition, the trajectory was analyzed using the statistical probability density of the proton asymmetry parameter (δ) across the 11-water wire (Figure 6e). In the no-field control simulation, the distribution of δ resolves as a broad, multi-peaked envelope spanning from 0.7 to 1.1 Å, representing a disordered, classical localized regime where the charge carrier remains confined within decoupled diabatic potential wells. Conversely, under the 180 Hz external driving field, this localized profile transitions into an intense, highly uniform quantum delocalized state centered at δ = 0.43 Å (Figure 6e). This highly symmetric distribution illustrates the generation of a coherent quantum mechanical superposition across the shared-proton coordinates, suppressing independent classical fluctuations. Driven by the coupled dynamic gating of the tryptophan–aspartate complex, the external field guides the entire 11-water network to operate as a single, translation-invariant quantum state. As the coherent wavepacket evolves over this flattened potential energy surface, the underlying quantum dynamics sample an infinite continuum of simultaneous transition pathways, thereby generating the broad Zundel continuum resolved in the spectrum. Consequently, the simulation suggests that the 180 Hz field flattens the local electric field gradient to shift the proton coordinate into a strictly anharmonically adiabatic pathway.

## 3. Discussion & Conclusions

The experimental evidence presented in this work establishes a profound new paradigm in quantum biology and condensed matter biophysics: the first direct observation of macroscopically extended, long-range phase coherence sustained across a room-temperature biological solution ensemble. By capturing un-attenuated, phase-resolved electromagnetic fields crossing macroscopic boundaries up to 40 mm, these results systematically challenge the conventional assumption that room-temperature thermal fluctuations inherently confine quantum-coherent transport to sub-nanometer, sub-picosecond boundaries. The core of this breakthrough resides in the mathematical purity of the distance-dependent metrics extracted from the statistical variance of the experimental PCA2 Fast Fourier Transform (FFT) data. When migrating from the near-field (2.0 mm) boundary out to the far-field (40.0 mm) destination, the system exhibits an anomalous spatial optimization: the primary frequency mode undergoes a highly adaptive blue-shift to 2050 cm^-1^ while its bandwidth (FWHM) contracts from 1375.98 cm^-1^ down to 1228.44 cm^-1^. In classical fluid mechanics, a wavepacket traveling through a high-permittivity dielectric solvent medium must undergo severe diffractive dispersion, broadening its linewidth and driving a sharp drop in resonance quality. The experimental observation of an optimized far-field Quality Factor (*Q*_40*mm*_ ≈ 1.5955) over the near-field distance (*Q*_2*mm*_ ≈ 1.4244) is a definitive indicator of spectral condensation - the signature of a self-stabilizing, non-dispersive propagating wavepacket that actively suppresses thermal entropy across macroscopic space.

This structural stability requires a multi-scale energy funneling mechanism that spans four orders of magnitude across the 10 to 10,000 cm^-1^ spectral region. UB-TDS reveals that the system accomplishes this by coupling light-mass, ultra-fast protonic translations to the heavy, slow collective low-frequency translations of the surrounding solvent network of interactions. The broad 884 cm^-1^ GSB, resolved in the transient spectrum of the 0.674 ps first time constant component for the 40.0 mm distance sample, provides the direct spectroscopic signature of a standalone solvent framework relaxation. This feature confirms that the local hydration network softens its collective torsional force constants to organize a highly compliant, low-friction wave channel. As the active Zundel core blue-shifts to adapt to local polaronic solvent cages, the fast phase components remain continuously coupled directly to the persistent, low-frequency collective oscillations resolved in the experimental phase spectra.

These quantum observables provide deep insight into the functional biology of the cytokine interferon-gamma (IFN-γ). As a critical signaling molecule orchestrating broad immune and inflammatory responses, IFN-γ must operate reliably across complex cellular environments where standard biochemical diffusion is heavily restricted. The emergence of a non-dispersive wavepacket suggests that the protein has evolutionarily optimized its nonlinear structural coordinates to act as a long-range communication switch. Rather than relying on passive, localized thermal collisions, the phase-synchronized alignment of the protein-solvent network effectively transforms the IFN-γ homodimer into a coherent electrodynamic antenna array. By utilizing a common collective frequency - the fundamental 1622 cm^-1^ water bending vibration - as a universal temporal clock to synchronize every stretching, bending, and lattice mode across the interface, the biological ensemble achieves the exact harmonic integer alignment required to eliminate the local Franck-Condon reorganization barrier. Consequently, these findings indicate that IFN-γ’s signaling mechanism is fundamentally driven by physical wave-mechanics. The macroscopic quantum pre-organization of the surface water dipoles allows the cytokine network to bypass thermal dissipation, projecting a robust, phase-coherent signaling field across extended boundaries. This establishes a new physical baseline for field-mediated cellular communication, demonstrating that quantum-scale phase synchronization is an active, functional component required to drive macroscopic biological signaling.

## 4. Materials and Methods

### 4.1 Experimental Measurements

#### 4.1.1 Time-resolved spectroscopy

##### 4.1.1.1 Ultrafast source, CPA and pulse compression

The experimental set-up is depicted in Supplementary Figure S10a. The pump source was a 1550 nm Erbium-based fiber laser oscillator (C-Fiber, Menlo Systems) delivering initial 70 fs pulses with an average power of 250 mW at a repetition rate of 100 MHz. The pump source output beam was divided by a non-polarizing beam splitter with a 10:90 reflection-to-transmission (R:T) split ratio. To provide a stable, high-efficiency driving field without exceeding material damage constraints, the transmitted pump pulses were routed into a custom Chirped-Pulse Amplifier (CPA) network. This sequence utilized a high-throughput fused-silica transmission-grating-based stretcher and compressor (Gitterwerk GmbH, 1000.0 lines/mm) to optimize peak power distribution during the gain phase. Amplification was driven by a single-stage fiber amplifier consisting of two 10 W, 915 nm semiconductor diode lasers (AeroDIODE) coupled to Er/Yb co-doped fibers (Coherent).

The amplified output of the CPA system was passed through an optical chopper operating at a modulation frequency of 6 kHz, immediately followed by a high-extinction polarizer. To achieve final dispersion management and to reduce the amplified optical temporal profile down to the sub-two-cycle regime, the pulse train was subsequently compressed using a second layer consisting of a multi-bounce configuration of custom high-order dispersive mirrors (UltraFast

Innovations GmbH). These specialized chirped mirrors are specifically engineered to provide simultaneously optimized Group Delay Dispersion (GDD) and Fourth-Order Dispersion (FOD) compensation, systematically counterbalancing the high-order phase errors and residual cubic/quartic chirp introduced by the transmission gratings and the nonlinear medium. We currently do not have access to a phase-resolved ultrafast diagnostic system, therefore an evaluation of this final free-space compression profile was performed by fitting the time-zero non-resonant cross-correlation instrument response function (IRF) amplitude profile (Supplementary Figure S10b), demonstrating an exceptionally tight fit convergence (R^2^ = 0.99665) and yielding an experimental Full Width at Half Maximum (FWHM) estimate pulse duration proxy of 2.98 fs.

##### 4.1.1.2 Multi-octave infrared generation inside the DAST lattice

This ultra-compressed, polarized 2.98 fs near-transform-limited optical pulse was focused onto an organic nonlinear 4-N,N-dimethylamino-4’-N’-methyl-stilbazolium tosylate (DAST) crystal (Swiss Terahertz, LLC). The sub-three-cycle temporal profile concentrates the electric field envelope into a compressed time window. This structural feature drives highly efficient intra-pulse difference frequency generation (optical rectification) within the organic DAST crystal, broadening the phase-coherent continuum (*27*) down to 10 cm^-1^. Concurrently, the short pulse duration drives a cascaded third-order nonlinear effect (χ(^3^) self-phase modulation) along the organic molecular chains, extending the coherent continuum continuously up into the mid- and near-infrared regimes out to 10,000 cm^-1^. Since the 1550 nm fundamental pump wavelength matches the group velocity dispersion minimum of the DAST crystal substrate, the overlapping χ(^2^) and χ(^3^) processes remain velocity-matched, eliminating spectral gaps or destructive phase interference across the entire multi-octave window.

##### 4.1.1.3 Time-resolved field detection (TD-EOS)

Three high-precision 90° off-axis parabolic mirrors (OAPMs) were utilized to collect, collimate, and focus the generated broadband infrared radiation. The secondary 10% reflection probe arm was guided through a zero-order half-waveplate (λ/2) and subsequently introduced coaxially into the system’s mechanical delay line through a center-drilled aperture hole in the second parabolic mirror. The co-propagating infrared field and optical probe beams were co-focused onto the primary protein sample cell and subsequently into a 100 μm thick <110>-oriented gallium phosphide (GaP) crystal to implement phase-resolved electro-optic sampling (TD-EOS). The resulting linear Pockels-effect polarization modulations were captured via a balanced photodiode arrangement and continuously digitized by a high-frequency digital lock-in amplifier (MFLI, Zurich Instruments) referenced directly to the 6 kHz chopper TTL sync clock.

##### 4.1.1.4 Sample configuration

The primary protein sample cell consisted of two high-resistivity, float-zone silicon windows (HRFZ-Si, TYDEX) featuring an optical diameter of 25.4 mm and a substrate thickness of 1.0 mm held within a sealed transmission cell. To systematically examine the non-local long-range fields, the second protein volume (functioning explicitly as the independent excitation source for the experiment) was housed in an analogous dual-silicon sealed sample cell. The excitation cell was fixed to a manual micrometer attached directly to a high-precision linear translation stage to regulate the relative spatial separation distance and was positioned at a fixed 90-degree orthogonal orientation entirely outside of the active optical beam path of the main infrared experimental spectroscopy assembly.

##### 4.1.1.5 Materials

Recombinant human interferon gamma (1 mg/mL, Tris HCl pH 8.0) was purchased from AB Biotechnology Limited and was used directly without further modification.

#### 4.1.2 Heterodyne lock-in detection

##### 4.1.2.1 Electrodynamic cavity design

The multi-scale open-air cavity resonator as illustrated in Supplementary Figures S13a - b was constructed using two independent, parallel sample holders mounted on high-precision manual translation stages to allow linear micro-positioning from a sub-wavelength near-field coordinate of 2.0 mm out to a macroscopic far-field coordinate of 40.0 mm. To ensure maximum field transparency and match the substrate constraints of the independent sub-picosecond time-resolved optical experiments, the sample cells were assembled from high-resistivity float-zone silicon (HRFZ-Si) windows. To rule out material-specific anomalies, secondary verification control sets were replicated with research-grade polystyrene windows, yielding statistically identical phase velocity profiles.

##### 4.1.2.2 Signal acquisition and heterodyne lock-in demodulation

A mechanical chopper wheel was integrated transversely into the cavity path at a 90-degree angle relative to the inter-sample field vectors of the propagating wavefront to create a continuous carrier modulation envelope. Active macromolecular field vectors escaping the cavity boundaries were captured by a wideband directional log-periodic copper microstrip antenna array and routed into a high-speed zero-bias semiconductor diode detector. To achieve localized, non-invasive near-field profiling without introducing metallic reflections or destabilizing the internal cavity boundary potentials, a stripped single-mode fiber (SMF-28) was employed as a dielectric scattering probe transversely within the sub-millimeter gap. The high-frequency electromagnetic carrier waves captured by the probe array were resolved using a Mini-Circuits ZV47-K44RMS+ Coaxial RF Power Detector, operating over a native calibrated frequency range of 10 MHz to 40.0 GHz. The resulting audio-frequency voltage envelope was fed directly into the digital lock-in amplifier (MFLI, Zurich Instruments). The input coupling was set to AC mode with an input impedance at 50 Ω to prevent impedance mismatches. The MFLI internally cross-correlated the input voltage against the mechanical chopper reference clock (1 kHz), streaming real-time complex demodulated polar coordinates of instantaneous amplitude (R) and phase (Φ) at a stabilized digital sampling rate. The entire measurement apparatus was enclosed within a RF (Faraday) enclosure (Shieldex GmbH).

### 4.2 Computational Measurements

#### 4.2.1 MD simulations

##### 4.2.1.1 Control Protein

The initial protein was taken from the pdb structure IEKU. The pdb structure contains a single monomer rather than the functional IFN-γ homodimer but it retains the conserved C-terminal KRKR structure that is absent in the other crystal structures of IFN-γ. For our MD simulations we converted the single monomer to a functional homodimer in VMD (*28*) by duplicating the coordinate set and applying spatial transformations via the Tk Console, and assigning distinct chain IDs. Subsequently the mutation at Asp11 in the crystal structure was back-mutated to His11. MD simulations were run with GROMACS 2026 (*29*) employing the all-atom CHARMM36 forcefield (*30*). The protein was placed in the center of a cubic periodic box (6.5 nm)^3^ and filled with 7576 SPC water molecules. Na^+^ and Cl^−^ atoms were added to neutralize the system charge. Energy minimization was performed using the steepest descent algorithm with 2000 steps and this was followed by a NPT equilibration simulation of 5 ns. The production simulations were performed under NVT conditions for 100 ns with temperature control using velocity rescale (*31*) with a coupling time constant of 0.1 ps. The cutoff distance for van der Waals forces and short-range electrostatic interactions was 1 nm, and the long-range electrostatic interaction was treated using the smooth Particle Mesh Ewald (PME) method (*32*, *33*). All bonds were constrained using the LINCS method (*34*, *35*).

##### 4.2.1.2 External electric field

The equilibrated system was exposed to an oscillating electric field of the form *E*(*t*) = *E*_0_cos(*ωt*) with a frequency of either 7.5 GHz or 150 GHz. The field was applied in the z-direction with an amplitude of 0.1 V/nm and run for an additional 50 ns.

##### 4.2.1.3 Full correlation Analysis (FCA)

The covariance matrix of the MD simulation trajectories was extracted with the gmx covar tool in GROMACS and the mutual information calculated with the FCA toolset (14) developed by the Grubmüller group.

#### 4.2.2 QM/MM Simulations

Hybrid quantum mechanics/molecular mechanics (QM/MM) simulations were performed using the GROMACS-CP2K interface (*36*). The QM region consisted of the side-chains of the proton donor groups (Trp36 and Asp41) as well as a 11-water wire as shown in Figure 6c. The QM region was treated with unrestricted Kohn-Sham density functional theory (UKS-DFT) using the generalized gradient approximation (GGA) formulated by Perdew, Burke, and Ernzerhof (PBE)(*37*). Long-range dispersion interactions were accounted for using Grimme’s empirical DFT-D3 corrections (*38*). The electronic structure was simulated under a net charge of +1 and a doublet spin multiplicity state. All-electron properties were treated utilizing the Gaussian and augmented plane-wave (GAPW) formalism (*39*). Molecularly optimized double-zeta valence polarized basis sets (DZVP-MOLOPT-GTH)(*40*) were paired with Goedecker-Teter-Hutter (GTH-PBE) dual-space pseudopotentials (*41*). The auxiliary plane-wave basis density grid was specified with a real-space grid cutoff of 400 Ry and a relative cutoff of 60 Ry. Self-consistent field (SCF) convergence was accelerated through the Orbital Transformation (OT) scheme (*42*) utilizing a DIIS minimizer, a step size of 0.15, and FULL_ALL preconditioning to an energy tolerance of 1.0 × 10^-6^ Ry.

The total system was enclosed in a cubic simulation cell of (6.5 nm)^3^ subject to full 3D periodic boundary conditions. Electrostatic interactions between the QM and MM domains were explicitly accounted for via Gaussian expansion of the electrostatic potential (ECOUPL GAUSS) using 12 Gaussians via the GEEP library (*43*). Long-range electrostatics were managed with a periodic multipole scheme configured with an electrostatic cutoff radius of 10.0 Å and an Ewald precision of 1.0 × 10^-6^. Covalent bonds intersecting the QM/MM boundary were handled using link atoms across 7 distinct link positions. Time-dependent perturbation was introduced by applying an external homogeneous electric field polarized along the Z-axis vector (0.0, 0.0, 1.0), specified with a field intensity amplitude of 9.723 × 10^-5^ a.u. and an initial spatial wavelength parameter of 1,665,513.65 a.u. The total dipole moments were calculated relative to the system’s center of mass.

## References

1. H.-G. Duan, V. I. Prokhorenko, R. J. Cogdell, K. Ashraf, A. L. Stevens, M. Thorwart, R. J. D. Miller, Nature does not rely on long-lived electronic quantum coherence for photosynthetic energy transfer. Proc. Natl. Acad. Sci. 114, 8493–8498 (2017).

2. M. I. Mallus, M. Aghtar, S. Chandrasekaran, G. Lüdemann, M. Elstner, U. Kleinekathöfer, Relation between Dephasing Time and Energy Gap Fluctuations in Biomolecular Systems. J. Phys. Chem. Lett. 7, 1102–1108 (2016).

3. D. M. Leitner, H. D. Pandey, K. M. Reid, Energy Transport across Interfaces in Biomolecular Systems. J. Phys. Chem. B 123, 9507–9524 (2019).

4. M. Lechelon, Y. Meriguet, M. Gori, S. Ruffenach, I. Nardecchia, E. Floriani, D. Coquillat, F. Teppe, S. Mailfert, D. Marguet, P. Ferrier, L. Varani, J. Sturgis, J. Torres, M. Pettini, Experimental evidence for long-distance electrodynamic intermolecular forces. Sci. Adv. 8, eabl5855.

5. N. Agmon, The Grotthuss mechanism. Chem. Phys. Lett. 244, 456–462 (1995).

6. A. Scott, Davydov’s soliton. Phys. Rep. 217, 1–67 (1992).

7. L. Cruzeiro, The Davydov/Scott Model for Energy Storage and Transport in Proteins. J. Biol. Phys. 35, 43–55 (2009).

8. W. W. Cleland, M. M. Kreevoy, Low-Barrier Hydrogen Bonds and Enzymic Catalysis. Science 264, 1887–1890 (1994).

9. N. Penkov, N. Penkova, Measurement of the Emission Spectra of Protein Solutions in the Infrared Range. Description of the Method and Testing Using Solution of Human Interferon Gamma as an Example. Front. Phys. 8 (2020).

10. M. Huber, M. Trubetskov, W. Schweinberger, P. Jacob, M. Zigman, F. Krausz, I. Pupeza, Standardized Electric-Field-Resolved Molecular Fingerprinting. Anal. Chem. 96, 13110–13119 (2024).

11. R. Kubo, The fluctuation-dissipation theorem. Rep. Prog. Phys. 29, 255 (1966).

12. J. G. Restrepo, E. Ott, B. R. Hunt, Onset of synchronization in large networks of coupled oscillators. Phys. Rev. E 71, 036151 (2005).

13. J. A. Acebrón, L. L. Bonilla, C. J. Pérez Vicente, F. Ritort, R. Spigler, The Kuramoto model: A simple paradigm for synchronization phenomena. Rev. Mod. Phys. 77, 137–185 (2005).

14. O. F. Lange, H. Grubmüller, Full correlation analysis of conformational protein dynamics. Proteins 70, 1294–1312 (2008).

15. F. Dahms, B. P. Fingerhut, E. T. J. Nibbering, E. Pines, T. Elsaesser, Large-amplitude transfer motion of hydrated excess protons mapped by ultrafast 2D IR spectroscopy. Science 357, 491–495 (2017).

16. M. Thämer, L. De Marco, K. Ramasesha, A. Mandal, A. Tokmakoff, Ultrafast 2D IR spectroscopy of the excess proton in liquid water. Science 350, 78–82 (2015).

17. A. Sirohiwal, J. John, Y. Kutin, R. Kumar, F. Baserga, V. Srinivas, H. Lebrette, M. C. Pöverlein, A. P. Gamiz-Hernandez, J. Heberle, M. Kasanmascheff, M. Högbom, V. R. I. Kaila, Low-barrier hydrogen bond powers long-range radical transfer in the metal-free ribonucleotide reductase. Proc. Natl. Acad. Sci. 123, e2529856123 (2026).

18. J.-W. Shin, N. I. Hammer, E. G. Diken, M. A. Johnson, R. S. Walters, T. D. Jaeger, M. A. Duncan, R. A. Christie, K. D. Jordan, Infrared Signature of Structures Associated with the H+(H2O)n (n = 6 to 27) Clusters. Science 304, 1137–1140 (2004).

19. J. M. Headrick, E. G. Diken, R. S. Walters, N. I. Hammer, R. A. Christie, J. Cui, E. M. Myshakin, M. A. Duncan, M. A. Johnson, K. D. Jordan, Spectral signatures of hydrated proton vibrations in water clusters. Science 308, 1765–1769 (2005).

20. M. Miyazaki, A. Fujii, T. Ebata, N. Mikami, Infrared spectroscopic evidence for protonated water clusters forming nanoscale cages. Science 304, 1134–1137 (2004).

21. P. Rebentrost, M. Mohseni, I. Kassal, S. Lloyd, A. Aspuru-Guzik, Environment-assisted quantum transport. New J. Phys. 11, 033003 (2009).

22. R. Biswas, W. Carpenter, J. A. Fournier, G. A. Voth, A. Tokmakoff, IR spectral assignments for the hydrated excess proton in liquid water. J. Chem. Phys. 146, 154507 (2017).

23. Y. Jia, S. Du, A. A. Seshia, Twenty-Eight Orders of Parametric Resonance in a Microelectromechanical Device for Multi-band Vibration Energy Harvesting. Sci. Rep. 6, 30167 (2016).

24. R. Pandya, R. Y. S. Chen, A. Cheminal, T. Thomas, A. Thampi, A. Tanoh, J. Richter, R. Shivanna, F. Deschler, C. Schnedermann, A. Rao, Observation of Vibronic-Coupling-Mediated Energy Transfer in Light-Harvesting Nanotubes Stabilized in a Solid-State Matrix. J. Phys. Chem. Lett. 9, 5604–5611 (2018).

25. Q. Yu, W. B. Carpenter, N. H. C. Lewis, A. Tokmakoff, J. M. Bowman, High-Level VSCF/VCI Calculations Decode the Vibrational Spectrum of the Aqueous Proton. J. Phys. Chem. B 123, 7214–7224 (2019).

26. K. N. Woods, Solvent-induced backbone fluctuations and the collective librational dynamics of lysozyme studied by terahertz spectroscopy. Phys. Rev. E 81, 031915 (2010).

27. A. Trisorio, M. Divall, B. Monoszlai, C. Vicario, C. P. Hauri, Intense sub-two-cycle infrared pulse generation via phase-mismatched cascaded nonlinear interaction in DAST crystal. Opt. Lett. 39, 2660–2663 (2014).

28. W. Humphrey, A. Dalke, K. Schulten, VMD: visual molecular dynamics. J. Mol. Graph. 14, 33–38, 27–28 (1996).

29. M. J. Abraham, T. Murtola, R. Schulz, S. Páll, J. C. Smith, B. Hess, E. Lindahl, GROMACS: High performance molecular simulations through multi-level parallelism from laptops to supercomputers. SoftwareX 1–2, 19–25 (2015).

30. B. R. Brooks, C. L. Brooks, A. D. Mackerell, L. Nilsson, R. J. Petrella, B. Roux, Y. Won, G. Archontis, C. Bartels, S. Boresch, A. Caflisch, L. Caves, Q. Cui, A. R. Dinner, M. Feig, S. Fischer, J. Gao, M. Hodoscek, W. Im, K. Kuczera, T. Lazaridis, J. Ma, V. Ovchinnikov, E. Paci, R. W. Pastor, C. B. Post, J. Z. Pu, M. Schaefer, B. Tidor, R. M. Venable, H. L. Woodcock, X. Wu, W. Yang, D. M. York, M. Karplus, CHARMM: the biomolecular simulation program. J. Comput. Chem. 30, 1545–1614 (2009).

31. G. Bussi, D. Donadio, M. Parrinello, Canonical sampling through velocity rescaling. J. Chem. Phys. 126, 014101 (2007).

32. U. Essmann, L. Perera, M. L. Berkowitz, T. Darden, H. Lee, L. G. Pedersen, A smooth particle mesh Ewald method. J. Chem. Phys. 103, 8577–8593 (1995).

33. T. Darden, D. York, L. Pedersen, Particle mesh Ewald: An N⋅log(N) method for Ewald sums in large systems. J. Chem. Phys. 98, 10089–10092 (1993).

34. B. Hess, P-LINCS: A Parallel Linear Constraint Solver for Molecular Simulation. J. Chem. Theory Comput. 4, 116–122 (2008).

35. B. Hess, H. Bekker, H. J. C. Berendsen, J. G. E. M. Fraaije, LINCS: A linear constraint solver for molecular simulations. J. Comput. Chem. 18, 1463–1472 (1997).

36. T. D. Kühne, M. Iannuzzi, M. Del Ben, V. V. Rybkin, P. Seewald, F. Stein, T. Laino, R. Z. Khaliullin, O. Schütt, F. Schiffmann, D. Golze, J. Wilhelm, S. Chulkov, M. H. Bani-Hashemian, V. Weber, U. Borštnik, M. Taillefumier, A. S. Jakobovits, A. Lazzaro, H. Pabst, T. Müller, R. Schade, M. Guidon, S. Andermatt, N. Holmberg, G. K. Schenter, A. Hehn, A. Bussy, F. Belleflamme, G. Tabacchi, A. Glöß, M. Lass, I. Bethune, C. J. Mundy, C. Plessl, M. Watkins, J. VandeVondele, M. Krack, J. Hutter, CP2K: An electronic structure and molecular dynamics software package - Quickstep: Efficient and accurate electronic structure calculations. J. Chem. Phys. 152, 194103 (2020).

37. J. P. Perdew, K. Burke, M. Ernzerhof, Generalized Gradient Approximation Made Simple. Phys. Rev. Lett. 77, 3865–3868 (1996).

38. S. Grimme, J. Antony, S. Ehrlich, H. Krieg, A consistent and accurate ab initio parametrization of density functional dispersion correction (DFT-D) for the 94 elements H-Pu. J. Chem. Phys. 132, 154104 (2010).

39. G. Lippert, J. Hutter, M. Parrinello, The Gaussian and augmented-plane-wave density functional method for ab initio molecular dynamics simulations. Theor. Chem. Acc. 103, 124–140 (1999).

40. J. VandeVondele, J. Hutter, Gaussian basis sets for accurate calculations on molecular systems in gas and condensed phases. J. Chem. Phys. 127, 114105 (2007).

41. S. Goedecker, M. Teter, J. Hutter, Separable dual-space Gaussian pseudopotentials. Phys. Rev. B 54, 1703–1710 (1996).

42. J. VandeVondele, J. Hutter, An efficient orbital transformation method for electronic structure calculations. J. Chem. Phys. 118, 4365–4369 (2003).

43. T. Laino, F. Mohamed, A. Laio, M. Parrinello, An Efficient Real Space Multigrid QM/MM Electrostatic Coupling. J. Chem. Theory Comput. 1, 1176–1184 (2005).

