## Supplementary Information for "Macroscopic Vibronic Coherence and Non-Dispersive Soliton Propagation Across a Protein-Solvent Interface"

Kristina N. Woods*
*Technical University of Munich*
*TUM School of Natural Sciences*
*Department of Chemistry*
*Lichtenbergstraße 4*
*85748 Garching*
*Germany*

***Suppementary Material Text***

### I. Experimental heterodyne detection of liquid water

We have conducted heterodyne detection experiments on samples of liquid water that serve as control measurements to our IFN-γ measurements detailed within the manuscript in Section 2.1 and in Figure 1. Intriguingly, these control experiments with water reveal a highly localized, short-range cooperative phenomenon that serves as a critical baseline for the macromolecular studies. While the single water control in Supplementary Figure S1c and configurations at distances > 2.0 mm (similar to the 15.0 mm distance sample shown in Figure S1b) exhibit only a single, standard step-jump into a fluctuating regime, the 2.0 mm configuration in Figure S1a uniquely undergoes a distinct secondary step-jump at $t = -1.0\ s$. This short-range staircase profile indicates that bulk water possesses an unexpected, distance-dependent structural susceptibility. When confined beneath a critical spatial boundary of 2.5 mm, the localized electrodynamic interactions within the solvent network are sufficient to drive a multi-stage, collective phase-locking event independently of any protein macro-dipoles.

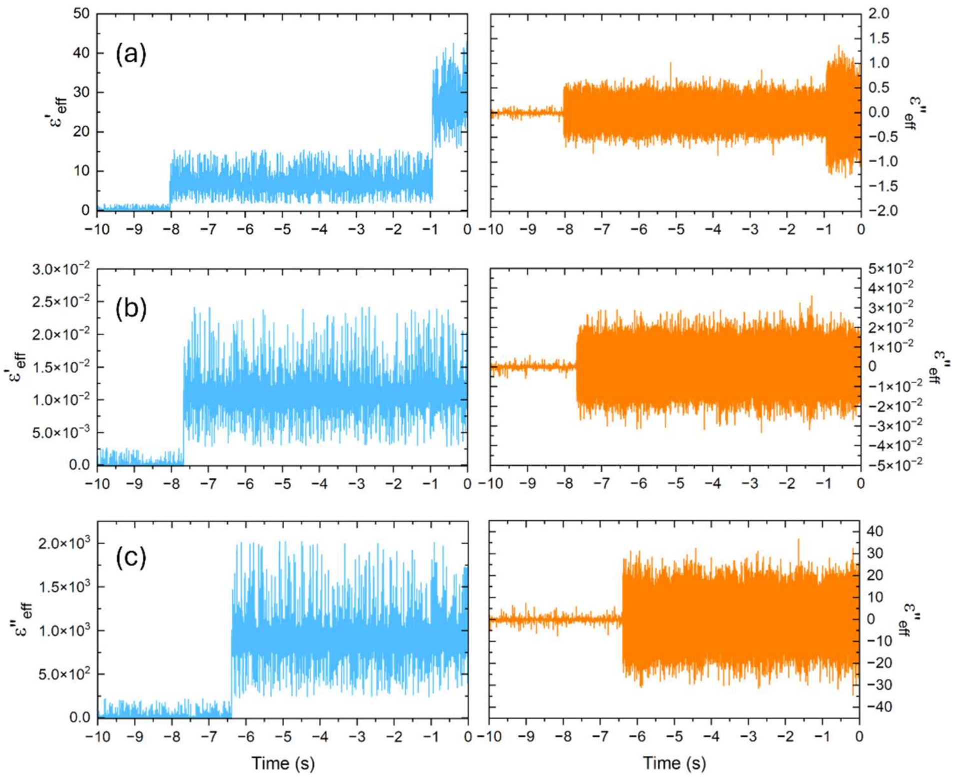


***Supplementary Figure S1*:** Heterodyne lock-in detection of the emitted-field of two liquid water samples at **(a)** 2.0 mm distance separation and **(b)** at 15.0 mm distance separation compared with the single water sample in **(c)**.

### II. Molecular Mechanisms of Field-Driven Dynamical Decoupling via Full Correlation Analysis

To elucidate the atomistic mechanisms driving the frequency-specific modulations observed experimentally at localized spatial wavelengths, we applied Full Correlation Analysis (FCA) to the steady-state regimes (100 ns) of the all-atom MD trajectories. By computing the non-linear mutual information ($I$) across the principal functional modes, we mapped the reorganization of the protein dimer's conformational landscape under both control conditions and high-frequency oscillating external electric fields (150 GHz and 7.5 GHz).

In the protein control, the protein dimer relies on a highly integrated, cooperative dynamical network to coordinate its biological functions. The native protein landscape is dominated by a prominent non-linear coupling between global collective fluctuations and localized catalytic sites. Specifically, the global motion of the helical core (**Mode 1**) is strongly coupled to the structural displacements of the essential proton donor groups located at the terminal domains (**Mode 6**, $I = 0.00601$; and **Mode 8**, $I = 0.00155$). This intense native correlation (Supplementary Table I) demonstrates that under standard isotropic thermal equilibrium, large-scale breathing and bending transitions of the structural core (Supplementary Figure S2) promote the spatial orientation and solvent accessibility of the catalytic proton donor groups.

The introduction of high-frequency external electric fields disrupts this native equilibrium network, altering the structural pathways into frequency-dependent regimes. Under the **150 GHz field** (corresponding to the $\lambda = 2.0$ mm experimental node), the native core-to-proton-donor coupling is effectively replaced with the $I$(Mode 1, Mode 6) value collapsing precipitously to 0.00013 (Supplementary Table II). In its place, the external field splits the macromolecular response into two distinct, newly emerged mechanical channels (Supplementary Figure S3):

- **Channel A (Gly26/C-Terminal Hinge compression):** A clear off-diagonal correlation peak emerges between **Mode 4** and **Mode 5** ($I = 0.00030$), isolating the high-amplitude displacements of the highly flexible Gly26 hinge loop and coupling them directly to the rigid frame of the C-terminal tail.
- **Channel B (Solvent-Exposed Helical Resonance):** Concurrently, **Mode 1** is no longer associated with the core-breathing dynamics but now establishes a localized non-linear correlation peak ($I = 0.00024$) with **Mode 3**, which corresponds to the coordinates of the solvent-exposed alpha-helices.

Mechanistically, because these outer, solvent-exposed helices possess large net dipole moments along their longitudinal axes, they act as macroscopic mechanical antennas that torque and oscillate in phase with the 150 GHz field. This intense structural resonance forces the outer helices to decouple from the core, while simultaneously whipping the adjacent flexible Gly26 loops into an isolated, independent non-linear path.

When the external driving frequency is shifted down to the **7.5 GHz field** (corresponding to the $\lambda$= 40.0 mm experimental node), the protein's overall energy landscape is transformed - collapsing the multi-component resonance into a single field-driven pathway. The original biological core-to-proton-donor correlation pathways (Supplementary Table III) are suppressed ($I$ = -0.00005). Instead of exciting multiple outer elements, the 7.5 GHz field drives an intense **dynamical caging effect**, wherein all collective transitions are focused exclusively within the localized C-terminal and Gly26 cluster (Supplementary Figure S4). This caging is marked by a single, tightly coordinated cooperative pathway where **Mode 5** displays dominant mutual information peaks with both **Mode 7** ($I$ = 0.00031) and **Mode 8** ($I$ = 0.00027). By confining the protein's conformational freedom to this singular, field-synchronized hinge interaction, the 7.5 GHz field effectively freezes out the classical, background thermal fluctuations that normally distribute energy across the broader helical core, isolating the local macromolecular frame from random thermal noise.

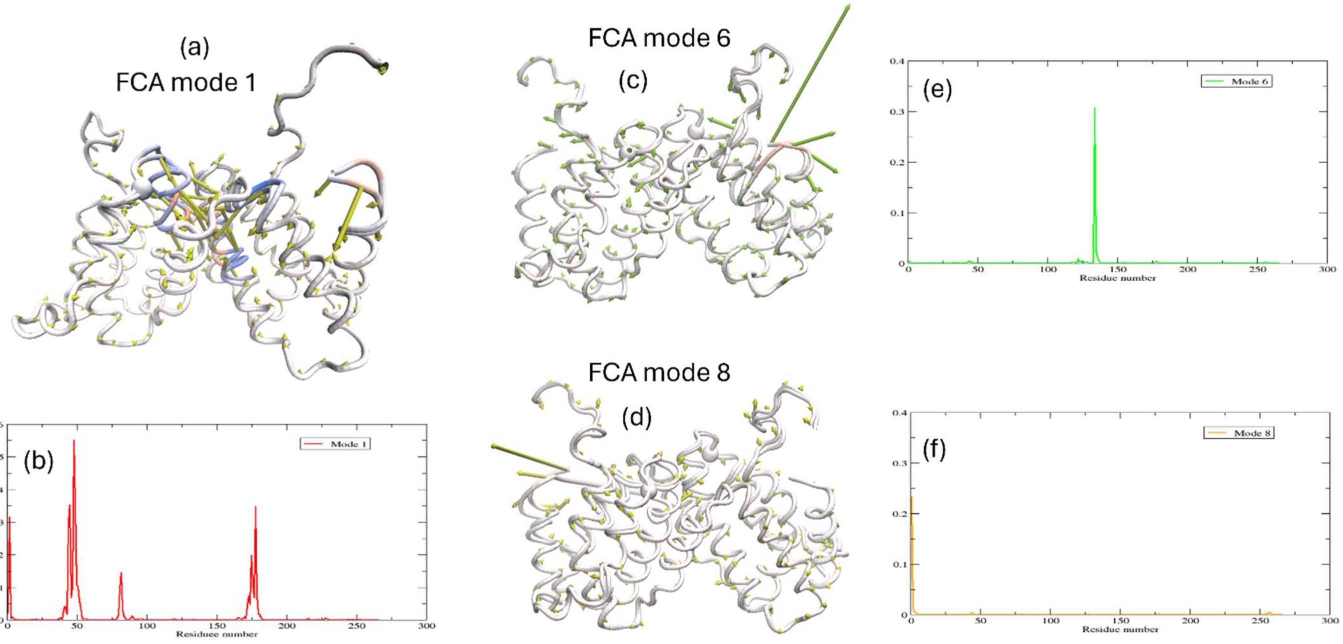


***Supplementary Figure S2:*** **(a)**, **(c)**, **(d)** are the modes in the control protein that are coupled in the FCA. **(b)**, **(e)**, and **(f)** are the root mean square fluctuations (RMSF) of the C-α residues in the respective mode.

| | M1 | M2 | M3 | M4 | M5 | M6 | M7 | M8 |
|---|---|---|---|---|---|---|---|---|
| M1 | 0 | 0.00002 | -0.00022 | -0.00004 | 0.00036 | 0.00601 | 0.00005 | 0.00155 |
| M2 | 0.00002 | 0 | 0.00017 | 0.0002 | 0.00001 | 0.00014 | -0.00016 | -0.00009 |
| M3 | 0.00022 | 0.00017 | 0 | -0.00003 | 0.00018 | -0.00023 | -0.0002 | 0.00016 |
| M4 | -0.00004 | 0.0002 | -0.00003 | 0 | 0.00023 | -0.00004 | 0.00011 | 0.00011 |
| M5 | -0.00036 | 0.00001 | 0.00018 | 0.00023 | 0 | -0.00005 | -0.00004 | 0.00025 |
| M6 | 0.00601 | 0.00014 | -0.00023 | -0.00004 | -0.00005 | 0 | -0.00015 | 0.00007 |
| M7 | 0.00005 | -0.00016 | -0.0002 | 0.00011 | -0.00004 | -0.00015 | 0 | -0.00005 |
| M8 | 0.00155 | -0.00009 | 0.00016 | 0.00011 | 0.00025 | 0.00007 | -0.00005 | 0 |

***Supplementary Table SI:*** The mutual information (MI) of the control protein from the MD simulations.

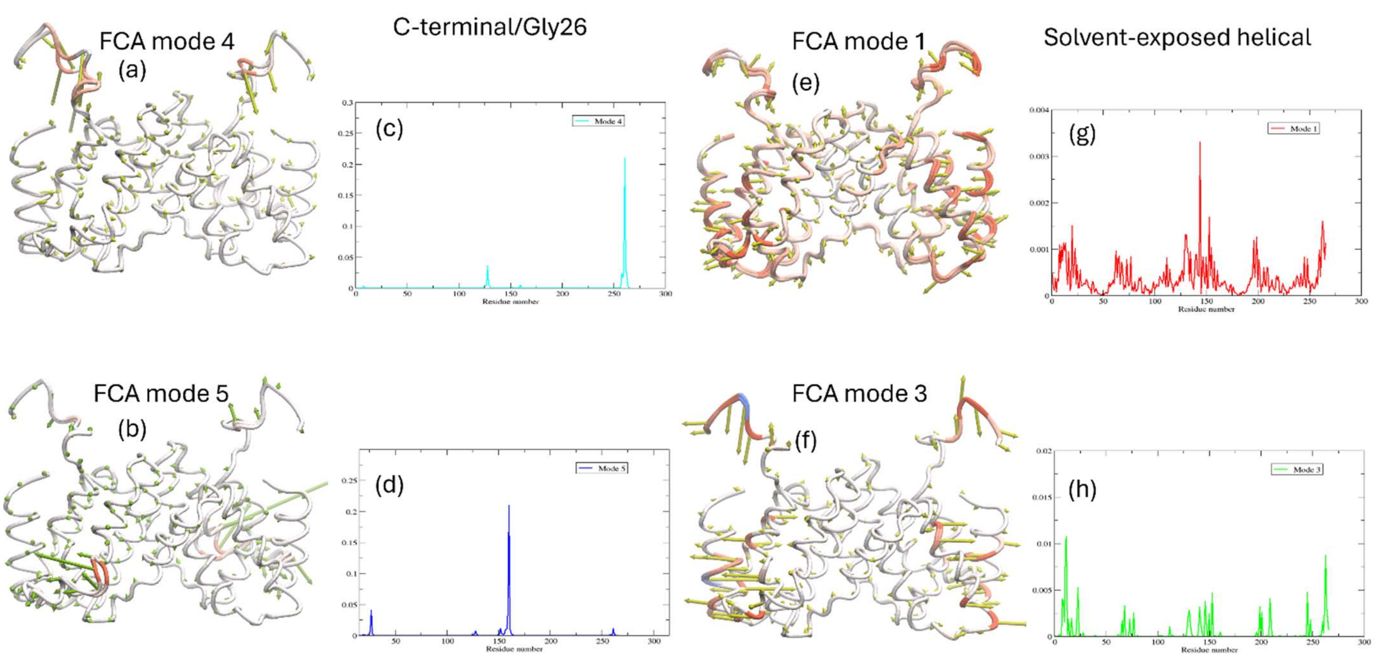


***Supplementary Figure S3*****:** **(a), (b), (e)** and **(f)** are depiction of the C-α dynamics of the modes in the 150 GHz MD simulation that are coupled in the FCA. **(c), (d), (g)** and **(h)** are RMSF of the C-α residues in the respective mode.

|  | M1 | M2 | M3 | M4 | M5 | M6 | M7 | M8 |
|---|---|---|---|---|---|---|---|---|
| M1 | 0 | 0.00005 | 0.00024 | 0.00011 | 0.00017 | 0.00013 | -0.00027 | -0.00007 |
| M2 | 0.00005 | 0 | -0.00008 | 0.00009 | 0.00001 | 0.00011 | 0.00006 | -0.00026 |
| M3 | 0.00024 | -0.00008 | 0 | -0.00005 | -0.00024 | -0.00005 | -0.00008 | 0.00008 |
| M4 | 0.00011 | 0.00009 | -0.00005 | 0 | 0.00030 | 0 | 0.00013 | 0.00002 |
| M5 | 0.00017 | 0.00001 | -0.00024 | 0.00030 | 0 | -0.00025 | 0.00014 | 0.00018 |
| M6 | 0.00013 | 0.00011 | -0.00005 | 0 | -0.00025 | 0 | -0.00008 | 0.00015 |
| M7 | -0.00027 | 0.00006 | -0.00008 | 0.00013 | 0.00014 | -0.00008 | 0 | -0.00016 |
| M8 | -0.00007 | -0.00026 | 0.00008 | 0.00002 | 0.00018 | 0.00015 | -0.00016 | 0 |

***Supplementary Table SII:*** The MI matrix from the IFN-γ simulation with a **150 GHz** applied electric field.

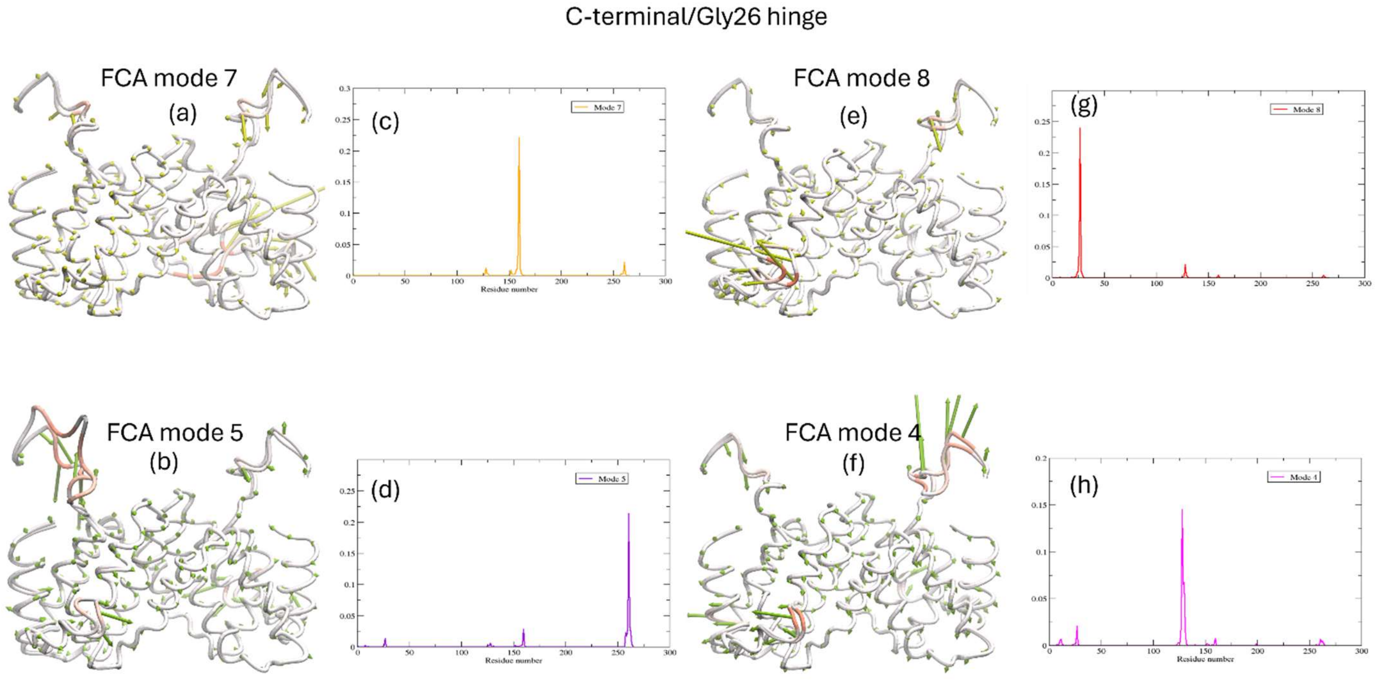


***Supplementary Figure S4:*** **(a)**, **(b)**, **(e)** and **(f)** are a depiction of the C-α dynamics of the modes in the **7.5 GHz** MD simulation that are coupled in the FCA. **(c)**, **(d)**, **(g)** and **(h)** are the RMSF of the C-α residues in the respective mode.

|  | M1 | M2 | M3 | M4 | M5 | M6 | M7 | M8 |
|---|---|---|---|---|---|---|---|---|
| M1 | 0.00000 | 0.00019 | -0.00033 | -0.00008 | 0.00001 | -0.00005 | 0.00007 | 0.00014 |
| M2 | 0.00019 | 0.00000 | 0.00011 | -0.00014 | 0.00014 | -0.00034 | -0.00006 | -0.00019 |
| M3 | -0.00033 | 0.00011 | 0.00000 | 0.00016 | -0.00018 | 0.00000 | 0.00004 | -0.00033 |
| M4 | -0.00008 | -0.00014 | 0.00016 | 0.00000 | 0.00020 | -0.00016 | 0.00001 | 0.00027 |
| M5 | 0.00001 | 0.00014 | -0.00018 | 0.00020 | 0.00000 | -0.00014 | 0.00031 | 0.00027 |
| M6 | -0.00005 | -0.00034 | 0.00000 | -0.00016 | -0.00014 | 0.00000 | -0.00002 | 0.00006 |
| M7 | 0.00007 | -0.00006 | 0.00004 | 0.00001 | 0.00031 | -0.00002 | 0.00000 | -0.00032 |
| M8 | 0.00014 | -0.00019 | -0.00033 | 0.00027 | 0.00027 | 0.00006 | -0.00032 | 0.00000 |

***Supplementary Table SIII:*** The MI matrix from the IFN-γ simulation with a **7.5 GHz** applied electric field

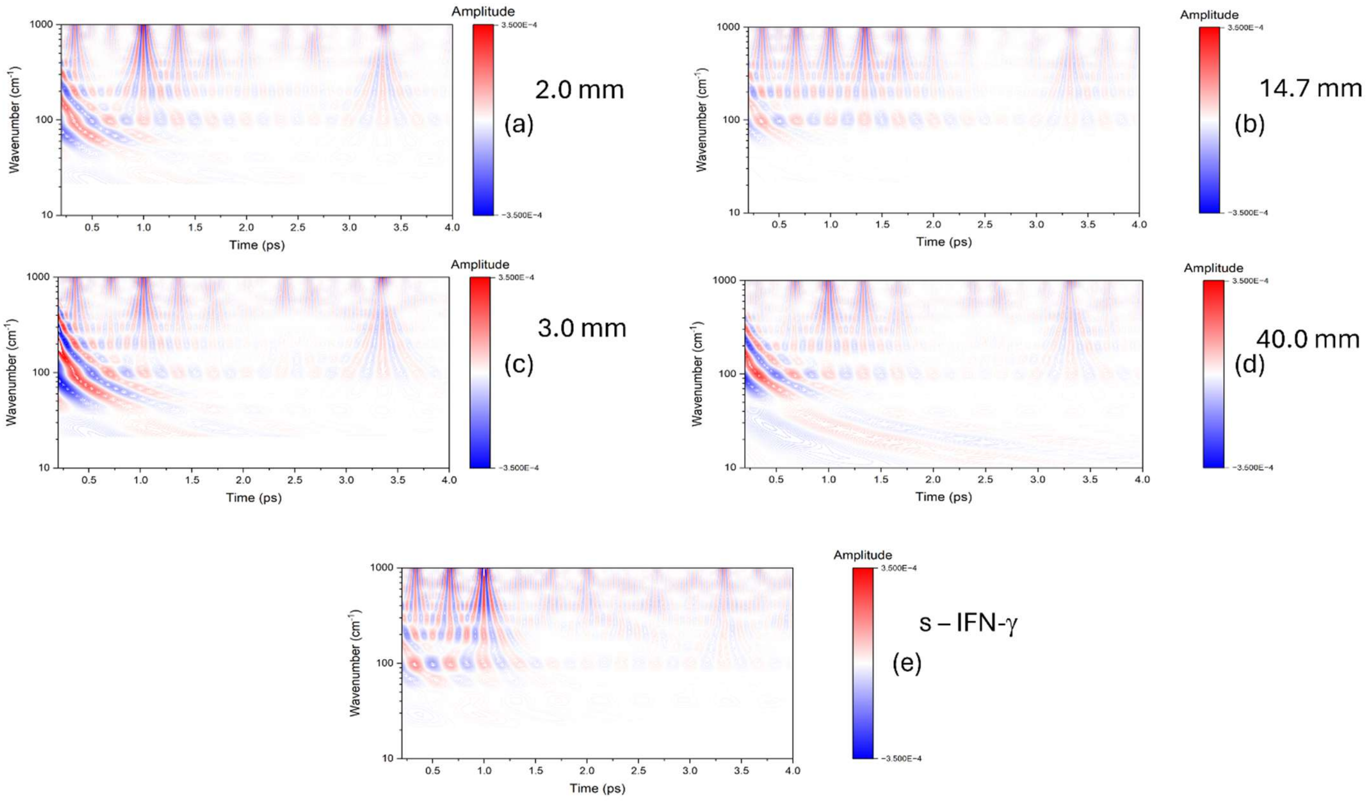


***Supplementary Figure S5:*** Plot of the UB-TDS phase spectral map in the 0.2 – 4.0 ps time interval and in the 10 – 1000 $cm^{-1}$ spectral region for **(a)** the 2.0 mm distance sample, **(b)** the 14.7 mm distance sample, **(c)** the 3.0 mm distance sample, **(d)**, 40.0 mm distance sample and **(e)** the s-IFN-γ sample. The spectral maps accentuate the coherent vibrational oscillations in the IFN-γ sample as a function of time.

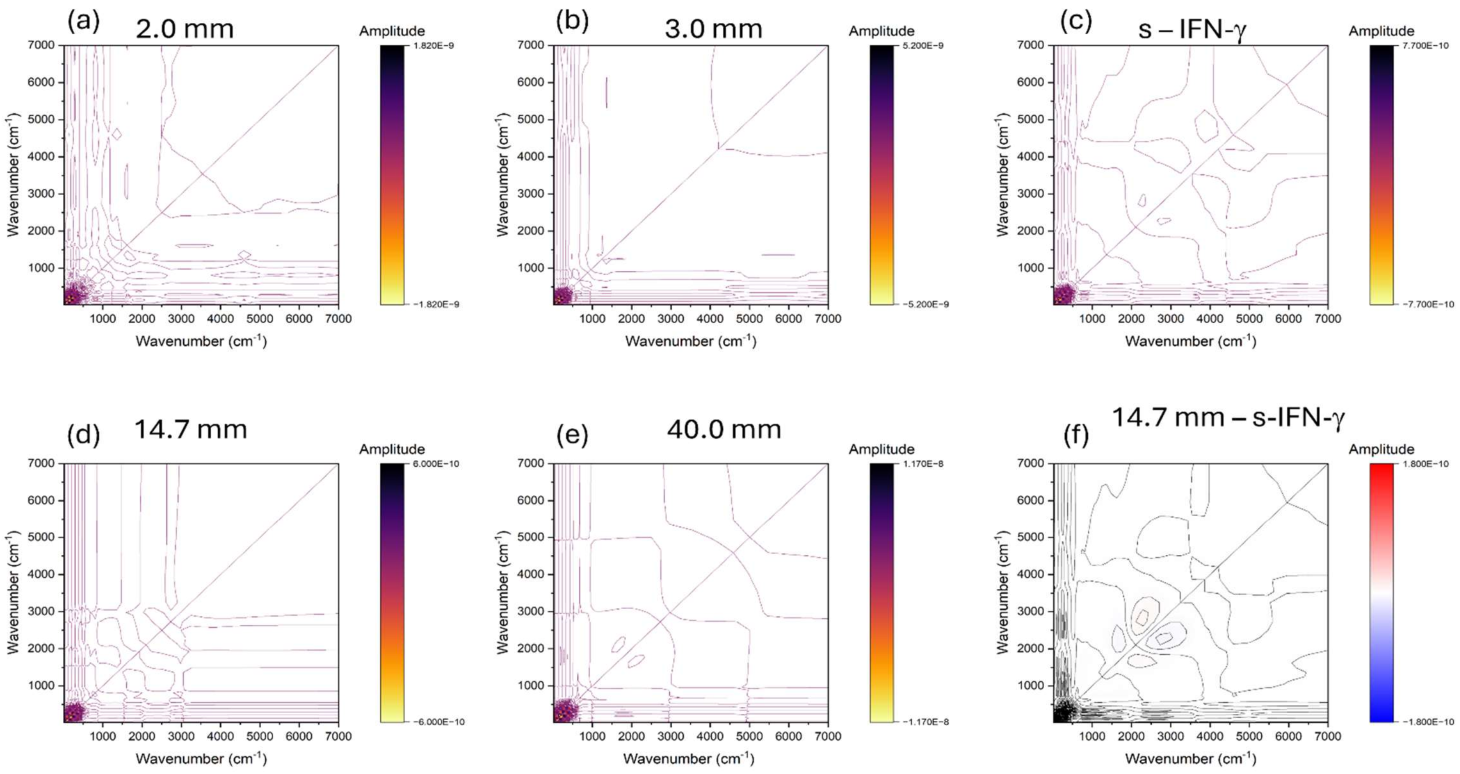


***Supplementary Figure S6:*** Two-dimensional (2-D) cross correlation spectral maps of the UB-TDS phase spectra from **(a)** the 2.0 mm distance sample, **(b)** the 3.0 mm distance sample, **(c)** the s-IFN-γ, **(d)** the 14.7 mm distance sample, **(e)** the 40.0 mm distance sample and **(f)** the subtraction of the 14.7 mm distance sample and the s – IFN-γ sample.

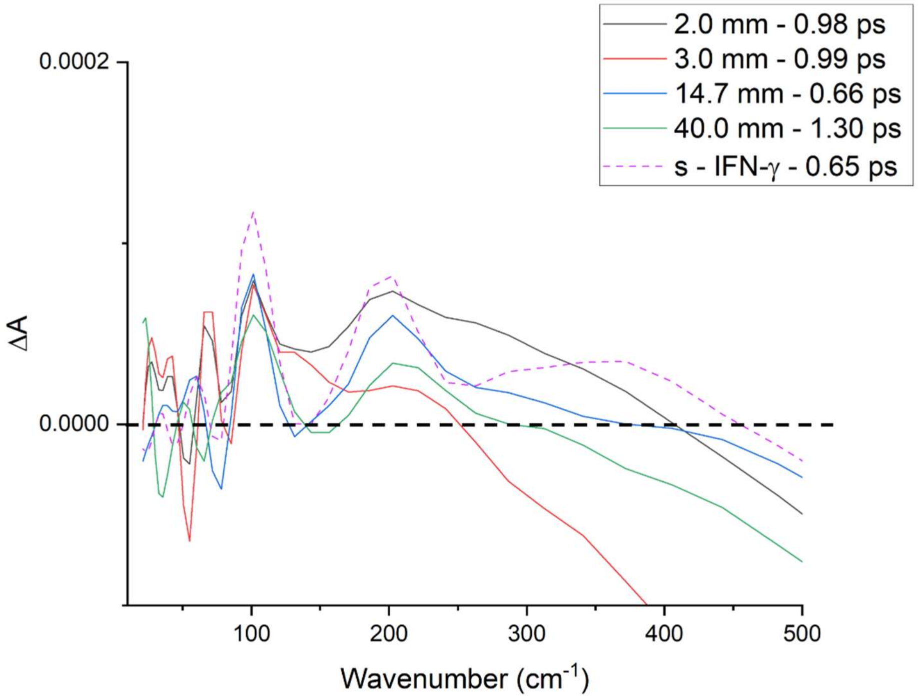


***Supplementary Figure S7:*** The 10 – 500 cm$^{-1}$ spectral region of the UB-TDS phase spectrum of the second time constant from Figure 3a

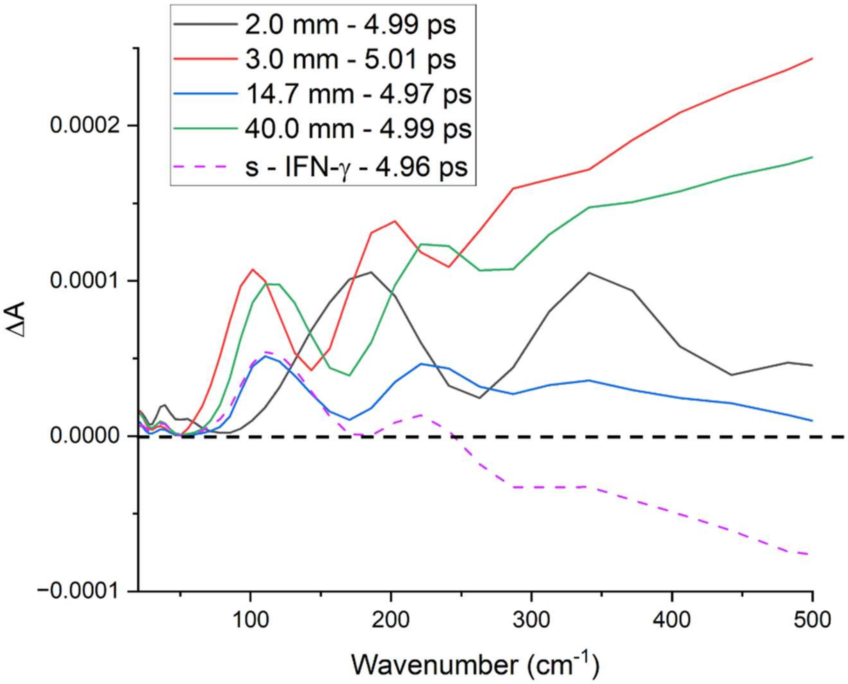


**Supplementary Figure S8:** The 10 – 500 cm$^{-1}$ spectral region of the UB-TDS phase spectrum of the fifth time constant from Figure 6a.

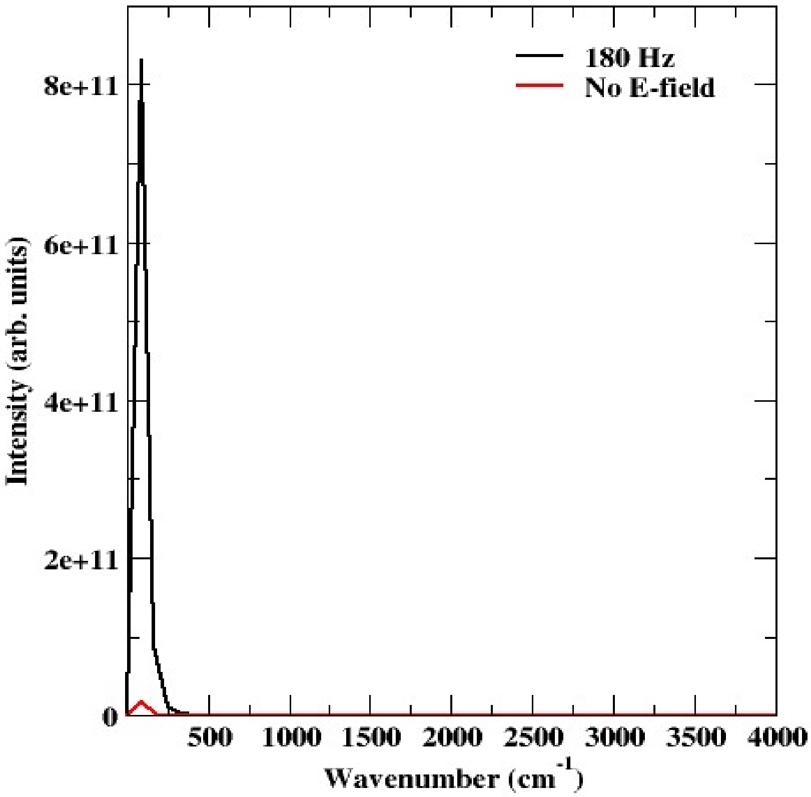


***Supplementary Figure S9:*** The infrared absorption spectrum computed from the Fourier transform of the quantum-mechanical total dipole autocorrelation function (DACF) tracked over the 350 fs QM/MM trajectories. The control protein (no electric field) is shown with a red trace, while the field-driven trajectory (180 Hz) is shown with a black trace. In this plot, the full amplitude of the ~85 cm$^{-1}$ mode is shown for both samples.

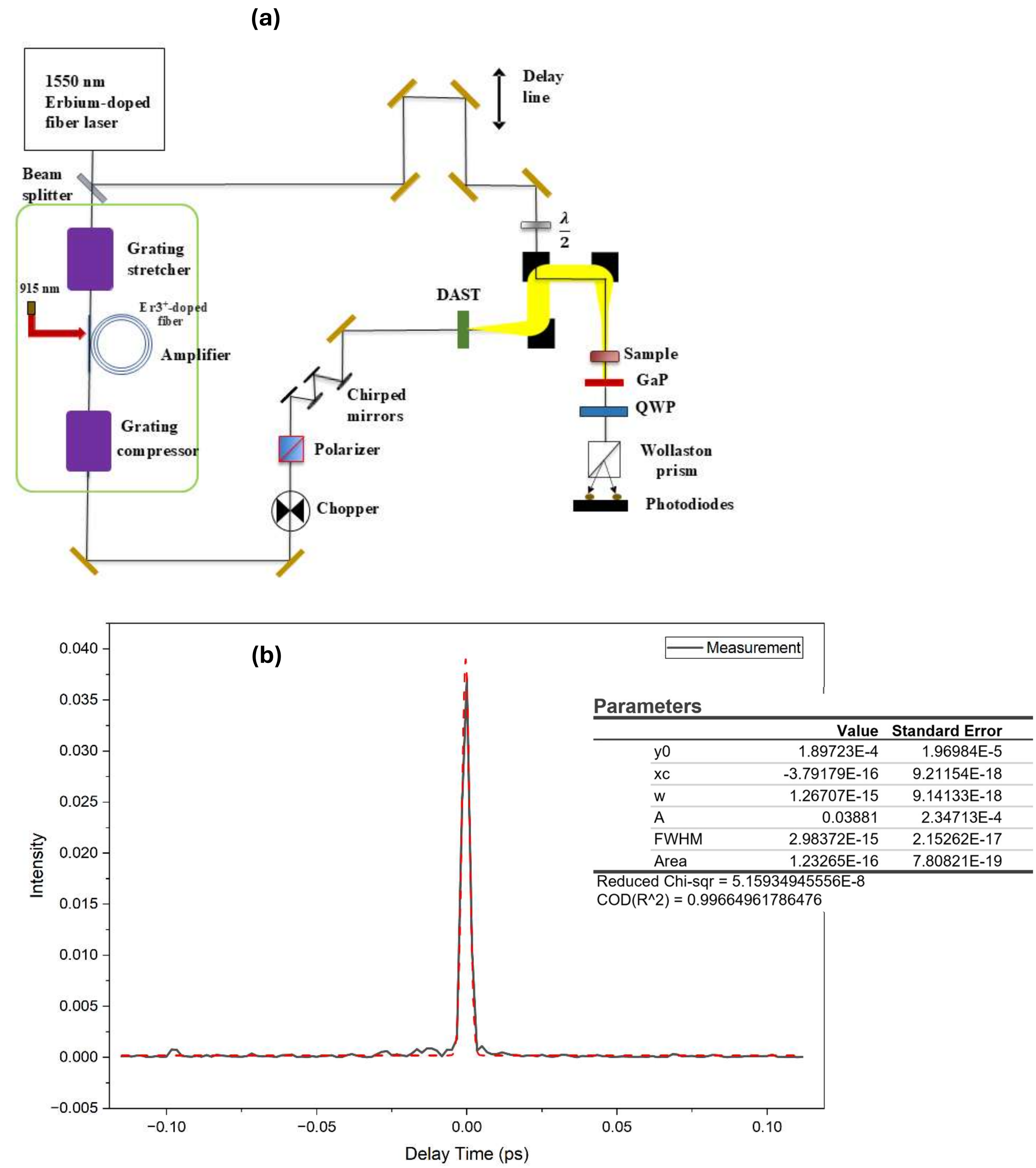


***Supplementary Figure S10***: A schematic of the optical set-up used for the time-resolved spectroscopy (UB-TDS) experiments described in this work. **(b)** Non-resonant cross-correlation instrument response function (IRF) amplitude profile of the experimentally resolved pulse duration from the time-resolved spectroscopy experiments.

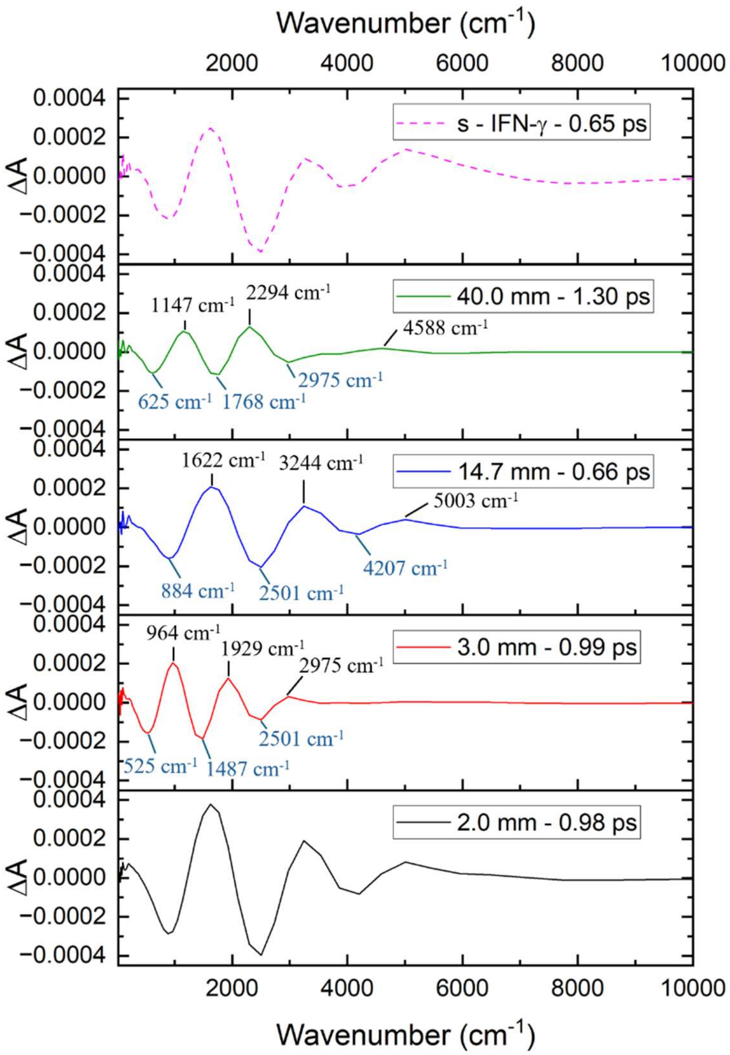


***Supplementary Figure S11:*** **Figure 3a** from the manuscript. The spectra are presented as a vertically stacked plot sharing common X- and Y- axes. The spectral features discussed in the text of the manuscript are explicitly labeled.

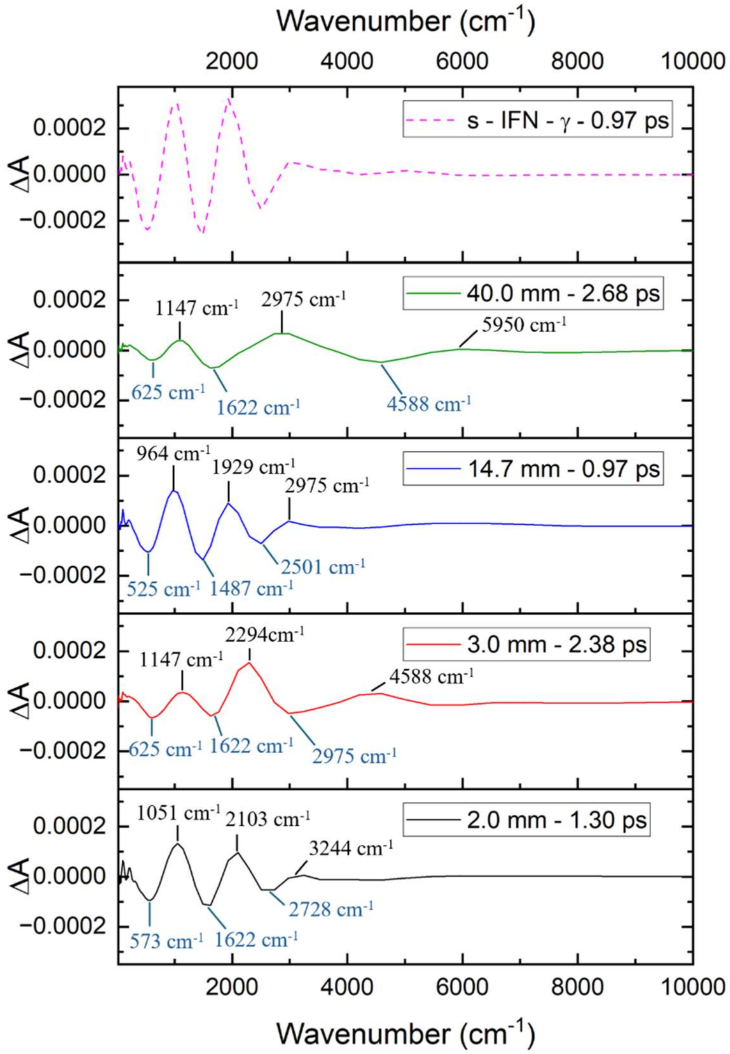


***Supplementary Figure S12:*** **Figure 4a** from the manuscript. The spectra are presented as a vertically stacked plot sharing common X- and Y- axes. The spectral features discussed in the text of the manuscript are explicitly labeled.

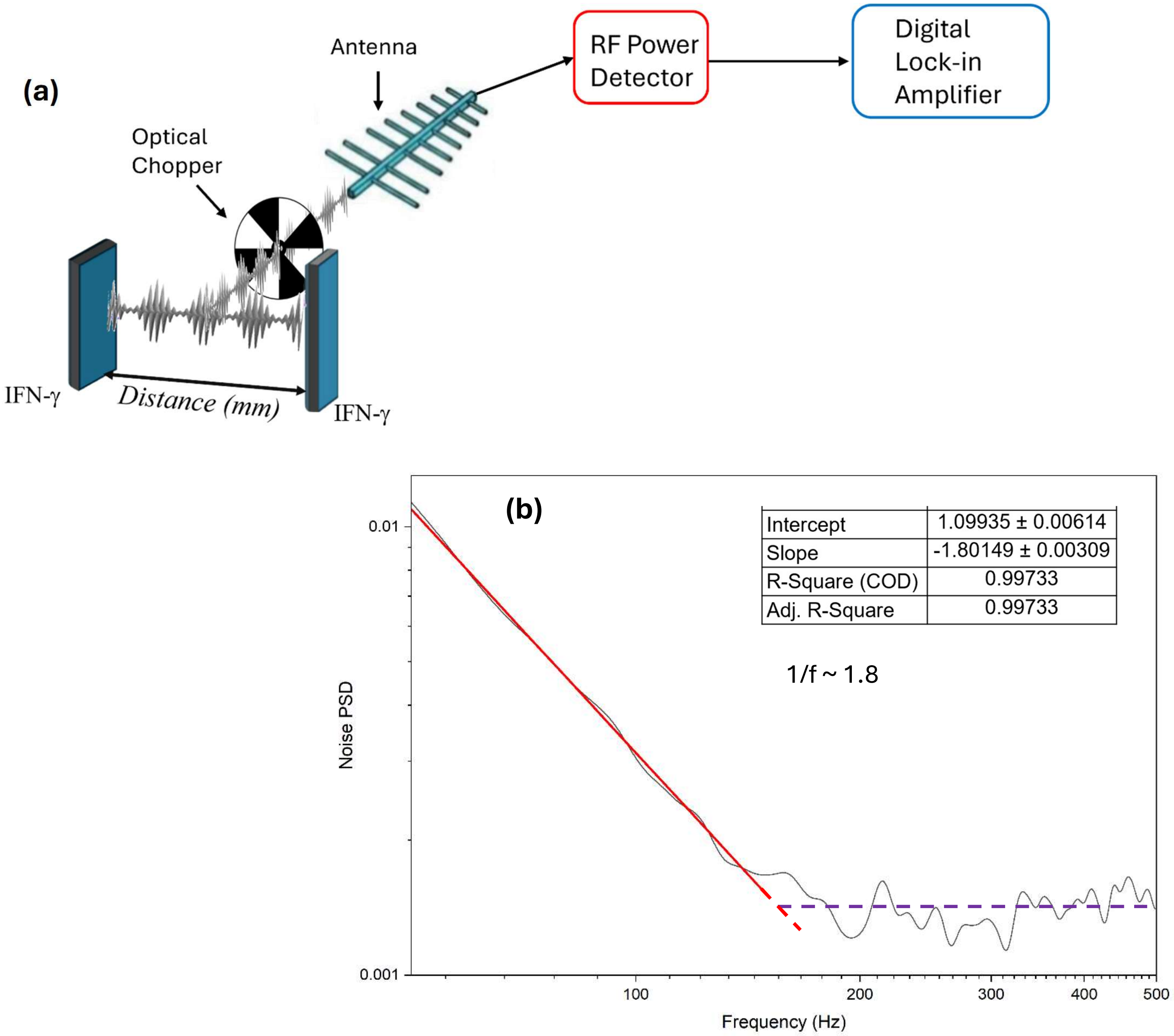


***Supplementary Figure S13*****:** **(a)** A schematic illustration of the heterodyne lock-in detection method used to measure the time-dependent phase velocity ($\Theta(t)$) and amplitude ($R(t)$) fields of the interference generated by the emission of the IFN–γ samples. **(b)** To establish the baseline sensitivity of the detection system, a noise spectrum measurement was performed within the RF enclosure. The 1 kHz chopper remained operational to account for any synchronous microphonic or electromagnetic artifacts from the motor, while the physical signal was isolated using aluminum foil placed between the chopper and the antenna. The resulting demodulated noise spectrum up to 150 Hz exhibited a positive linear slope of approximately 1.8. This rising noise profile is attributed to the internal phase noise dynamics of the RF detector system and localized frequency fluctuations, defining the absolute minimum detectable signal floor for the experiment at the 1 kHz operating frequency.
